# Genome-wide scan identifies novel genetic loci regulating salivary metabolite levels

**DOI:** 10.1101/687350

**Authors:** Abhishek Nag, Yuko Kurushima, Ruth C. E. Bowyer, Philippa M. Wells, Stefan Weiss, Maik Pietzner, Thomas Kocher, Johannes Raffler, Uwe Völker, Massimo Mangino, Timothy D. Spector, Michael V. Milburn, Gabi Kastenmüller, Robert P. Mohney, Karsten Suhre, Cristina Menni, Claire J. Steves

**Author notes:** **Corresponding author**, Dr. Claire J. Steves, Department of Twin Research and Genetic Epidemiology, King’s College London, London, UK.

## Abstract

Saliva, as a biofluid, is inexpensive and non-invasive to obtain, and provides a vital tool to investigate oral health and its interaction with systemic health conditions. There is growing interest in salivary biomarkers for systemic diseases, notably cardiovascular disease. Whereas hundreds of genetic loci have been shown to be involved in the regulation of blood metabolites leading to unprecedented insights into the pathogenesis of complex human diseases, little is known about the impact of host genetics on salivary metabolites. Here we report the first genome-wide association study exploring 476 salivary metabolites in 1,419 subjects of European ancestry from the TwinsUK cohort (discovery phase). A total of 14 salivary metabolites were significantly associated (p<10^−10^) with genetic variants that mapped to 11 distinct loci, most of which replicated in the Study of Health in Pomerania (SHIP-2) cohort. Interestingly, while only a limited number of the loci that are known to regulate blood metabolites were also associated with salivary metabolites in our study, we identified several novel saliva-specific locus-metabolite associations, including associations for the *AGMAT* (with the metabolites 4-guanidinobutanoate and beta-guanidinopropanoate), *ATP13A5* (with the metabolite creatinine) and *DPYS* (with the metabolites 3-ureidopropionate and 3-ureidoisobutyrate) loci. Our study suggests that there are biological pathways which are specific to the regulation of the salivary metabolome. In addition, some of our findings may have clinical relevance, such as the utility of the pyrimidine (uracil) degradation metabolites in predicting 5-fluorouracil toxicity and the role of the agmatine pathway metabolites as biomarkers of oral health.

## Introduction

Metabolic reactions pervade every aspect of human physiology, abnormalities in which underlie a plethora of human diseases^1^. Investigating the genetic underpinnings of population-wide variation of metabolites can offer novel insights into human metabolism and diseases, in addition to providing potential therapeutic targets to modulate metabolite levels. Large-scale genetic association studies have so far identified hundreds of loci that regulate the levels of metabolites in blood^2–6^, and to a lesser extent in other biospecimens as well^7–9^. Previous studies have shown that genetic variants on average explain a greater proportion of trait variance for metabolites compared to what is generally observed for complex traits^3,4^, highlighting the utility of metabolites as intermediate traits for dissecting the genetics of complex diseases.

Saliva is an abundantly produced biofluid, and it can be obtained in an inexpensive and non-invasive manner, without the need for healthcare professionals. It is mainly composed of water (>99%) and several other minor constituents such as mucous, digestive enzymes, cytokines, immunoglobulins, antibacterial peptides and low molecular weight metabolites^10^.

Recent advances in metabolomic profiling allow quantification of hundreds of metabolites belonging to diverse biochemical pathways in large population samples^4,11^. In 2015, the Human Metabolome Database (HMDB) incorporated data on the ‘salivary metabolome’ which included 853 salivary metabolites that were systematically characterized using a multiplatform approach^12^. Since saliva is separated from the systemic circulation by just a thin layer of cells, which allows passive and active exchange of substances^13^, it provides a reflection of not just oral health but the functioning of other organ systems as well^14^. Indeed, a number of studies have previously reported associations between oral health and systemic conditions such as cardiovascular diseases, diabetes, autoimmune diseases, mental health disorders, dementia, among others^15–19^. Therefore, investigation of salivary metabolites could not only provide novel biomarkers but also further our understanding of biological pathways underlying oral as well as general health conditions.

Here we report a genome-wide association analysis (based on 1000 Genomes imputed data) for 476 salivary metabolites in the population-based TwinsUK study, followed by replication in the population-based Study of Health in Pomerania (SHIP-2) cohort.

## Material and methods

### (I) Discovery phase

#### Study population

The discovery phase of the study was conducted in the TwinsUK cohort, an adult twin registry comprising of healthy volunteers, based at St. Thomas’ Hospital in London^20^. Twins gave fully informed consent under a protocol reviewed by the St. Thomas’ Hospital Local Research Ethics Committee. Subjects of European ancestry with available genotype data and for whom salivary metabolite profiling was done on a fasting state sample were included in our study (N = 1,419; mean age = 62.2 years; *%* females = 92.7).

#### Genotyping, imputation and QC

Subjects were genotyped in two different batches of approximately the same size, using two genotyping platforms from Illumina: 300K Duo and HumanHap610-Quad arrays. Whole genome imputation of the genotypes was performed using the 1000 Genomes reference haplotypes^21^, further details of which are provided in Moayyeri *et al* (2013). Stringent QC measures, including minimum genotyping success rate (>95%), Hardy-Weinberg equilibrium (p>10^−6^), minimum MAF (>0.5%) and imputation quality score (INFO>0.5), retained ~9.6 million variants for genome-wide analysis.

#### Saliva sample collection

Saliva samples were obtained by asking the fasted volunteer to spit as much saliva as possible into an empty sterile pot over a period of 10 minutes. The saliva samples were immediately refrigerated and then frozen at −80°C (usually within 4 hours of sample collection) before further processing. Following that, the samples were shipped on dry ice for metabolite profiling at Metabolon Inc., Durham, USA (See **Supplemental methods (I)** for further details on sample processing).

#### Metabolic profiling of saliva samples

Metabolite concentrations in the saliva samples were estimated using Ultrahigh Performance Liquid Chromatography-Tandem Mass Spectroscopy (UPLC-MS) i.e. chromatographic separation, followed by full-scan mass spectroscopy, to record all detectable ions in the samples (see **Supplemental methods (II)** for further details). Based on their unique ion signatures (chromatographic and mass spectral), 997 distinct metabolites were identified, of which 823 had known chemical identity at the time of analysis. The 823 known metabolites were broadly classified into 8 metabolic groups (amino acids, peptides, carbohydrates, energy, lipids, nucleotides, cofactors and vitamins, and xenobiotics) as described in the KEGG (Kyoto Encyclopedia of Genes and Genomes) database^22^. The 8 metabolic groups were further subdivided into 99 distinct biochemical pathways.

Raw metabolite values were normalised for the volume and osmolality measurement of the saliva samples. The normalised metabolite values were then log-transformed, and scaled to uniform mean 0 and standard deviation 1. Of the 823 known metabolites, 476 were retained for analysis based on presence of measurement in more than 80% samples. For the samples with missing values for these 476 metabolites, data were imputed using the run day minimum value for the metabolite. The resulting imputed dataset of the 476 metabolites was used for further analysis.

#### Genome-wide association analysis of salivary metabolites

##### (i) Primary genome-wide association analysis

For each of the 476 metabolites, a linear mixed-model was fitted to test the association between the metabolite (dependent variable) and genome-wide variants (independent variable). Age, sex and time of saliva sample collection were included as covariates in the model. The score test implemented in GEMMA^23^, which utilises a sample kinship matrix (estimated using a subset of ~500,000 variants) to account for the twin structure or relatedness in the TwinsUK data, was used to assess significance of the associations. A genome-wide and metabolome-wide significance cut-off of p<10^−10^ (corresponding to the conventional genome-wide significance threshold of 5×10^−8^, corrected for 476 metabolites) was used to identify significant variant-metabolite associations. For each locus that was significantly associated with a metabolite, we reported the variant with the lowest association p-value.

##### (ii) Testing loci identified in the primary analysis for additional metabolite associations

Next, we focused just on the loci that were identified in the primary stage of association testing to look for additional variant-metabolite associations for those loci. For each locus that was identified in the primary analysis, we clumped all variants located within a 100 Mb block and with LD (r^2^) > 0.2, and checked for additional metabolite associations at a significance threshold of p<10^−6^ (corresponding to p=0.05, corrected for 476 metabolites and a prior assumption of about 100 associated loci).

##### (iii) Testing the significantly associated loci using metabolite data from other biospecimens

For each locus-metabolite association, we further assessed the most significant variant-metabolite pair by using measurements for the respective metabolite in serum^4^ and faecal samples^7^ of the TwinsUK subjects (provided the metabolite was measured in that biospecimen). We tested only those serum and faecal samples that overlapped with the ones in saliva and were collected within 5 years of the saliva samples (in order to ensure that, for a given individual, samples from the different biospecimens being tested were obtained within a certain period of one another). Association testing for serum and faecal metabolites was done using an identical model to that described for the analysis of salivary metabolites.

##### (iv) Conditional analysis for the significantly associated loci

###### (a) Detection of secondary association signals

We used approximate conditional analysis, as implemented on GCTA^24^, to test whether any of the associated loci had multiple distinct i.e. secondary association signals, at a “locus-wide” significance threshold of p<10^−5^. For each associated locus, all variants that surpassed the study-wide significance threshold (p<10^−10^) were conditioned on the most significantly associated variant at that locus (using the association summary statistics). For the conditional analysis, we used genotype data from the complete TwinsUK dataset (N=5,654) to model LD patterns between variants.

###### (b) Adjustment for periodontal disease status

Since salivary metabolites are known to be associated with periodontal disease (PD)^11^, for each locus-metabolite association, we additionally adjusted the most significant variant-metabolite pair for PD status. Self-reported gingival bleeding, and a history of gum disease or tooth mobility were used as indicators of PD in TwinsUK^25^ (270 PD cases; 1,083 controls).

#### Expression quantitative trait locus (eQTL) analysis

We used the version 7 data release of the Genotype-Tissue Expression (GTEx) project (accessed 15 April 2019), which was based on RNA-Seq data obtained from 48 non-diseased tissue sites across ~1,000 individuals, to test whether the most significant variant at each associated locus had an eQTL effect on transcripts located within a 1 Mb window of the variant.

#### Annotation of associations using reference databases

We searched the NHGRI GWAS catalogue (accessed 15 April 2019) for previous disease associations for the significantly associated loci that were identified in our study. We also searched the OMIM database (accessed 15 April 2019) to check the candidate genes at the associated loci for a causal link with inborn errors of metabolism. Moreover, we also queried the HMDB^12^ and KEGG^22^ databases to identify biochemical pathways and known disease associations for the associated metabolites.

### (II) Replication phase

The replication phase was performed in the Study of Health in Pomerania (SHIP-2), a population-based study comprising of European ancestry subjects, conducted in the northeastern area of Germany. Further details of SHIP-2, including cohort details, genotyping and imputation, and saliva sample collection are provided in **Supplemental methods (III)**. Metabolic profiling for SHIP-2 saliva samples (N=1,000) was performed using an identical process to that described for TwinsUK.

Since the method of saliva sample collection in SHIP-2 (chewing on a piece of cotton) meant that the sample thus obtained represented stimulated saliva, normalisation of the metabolite measurements for sample osmolality was not considered necessary. The fact that the salivary osmolality values in SHIP-2 had a much narrower distribution compared to that in TwinsUK verified our rationale (**Figure S1**).

For each locus-metabolite association that was identified in the discovery phase, we tested the most significantly associated variant using a linear regression model was fitted on R (version 3.5.2). Covariates used in the association model were similar to those used for the discovery phase.

### (III) Testing the significantly associated salivary metabolites with phenotypes of interest

We wanted to test how salivary metabolites that were regulated by genetic loci related to relevant phenotypes. For that, we selected the metabolites that were uniquely associated in saliva i.e. the ones for which a genetic association had not been previously reported in blood, and tested them with phenotypes (diseases / traits / adverse drug effects) relating to the metabolite or its associated biochemical pathway. We obtained the relevant phenotype information from the TwinsUK database, selecting one twin per pair (N=1,426). The phenotype association analysis was performed on R (version 3.5.2) by fitting a linear regression model to test the association between the salivary metabolite and the disease / trait / adverse drug effect (adjusted for age and sex).

## Results

### Identification of novel genetic loci regulating salivary metabolite levels

Primary genome-wide discovery analysis in TwinsUK identified 13 metabolites that were significantly associated with genetic loci after correcting for multiple testing (p<10^−10^). Furthermore, when we narrowed our analysis to just the loci that were identified in the primary stage, one additional metabolite was found associated (p<10^−6^). Consequently, a total of 14 distinct locus-metabolite associations (hereafter, referred to as ‘mQTLs’) were identified in the discovery phase of our study. The set of significantly associated variants mapped to 11 distinct genetic loci, which have been referred to by the name(s) of the overlapping or the nearest gene(s) (**Figure 1**). Three of those loci (*AGMAT, SLC2A9* and *DPYS*) were associated with two metabolites each (**Table 1**). In all three instances, the two metabolites regulated by the same locus were correlated (Pearson’s r^2^ for the metabolite pairs ranged between 0.42 - 0.84) (**Figure S2**). On the other hand, none of the 14 metabolites that was associated in our study had more than one significant locus. Quantile-quantile (QQ) plots for the significantly associated metabolites are provided in **Figure S3**.

**Figure 1:**
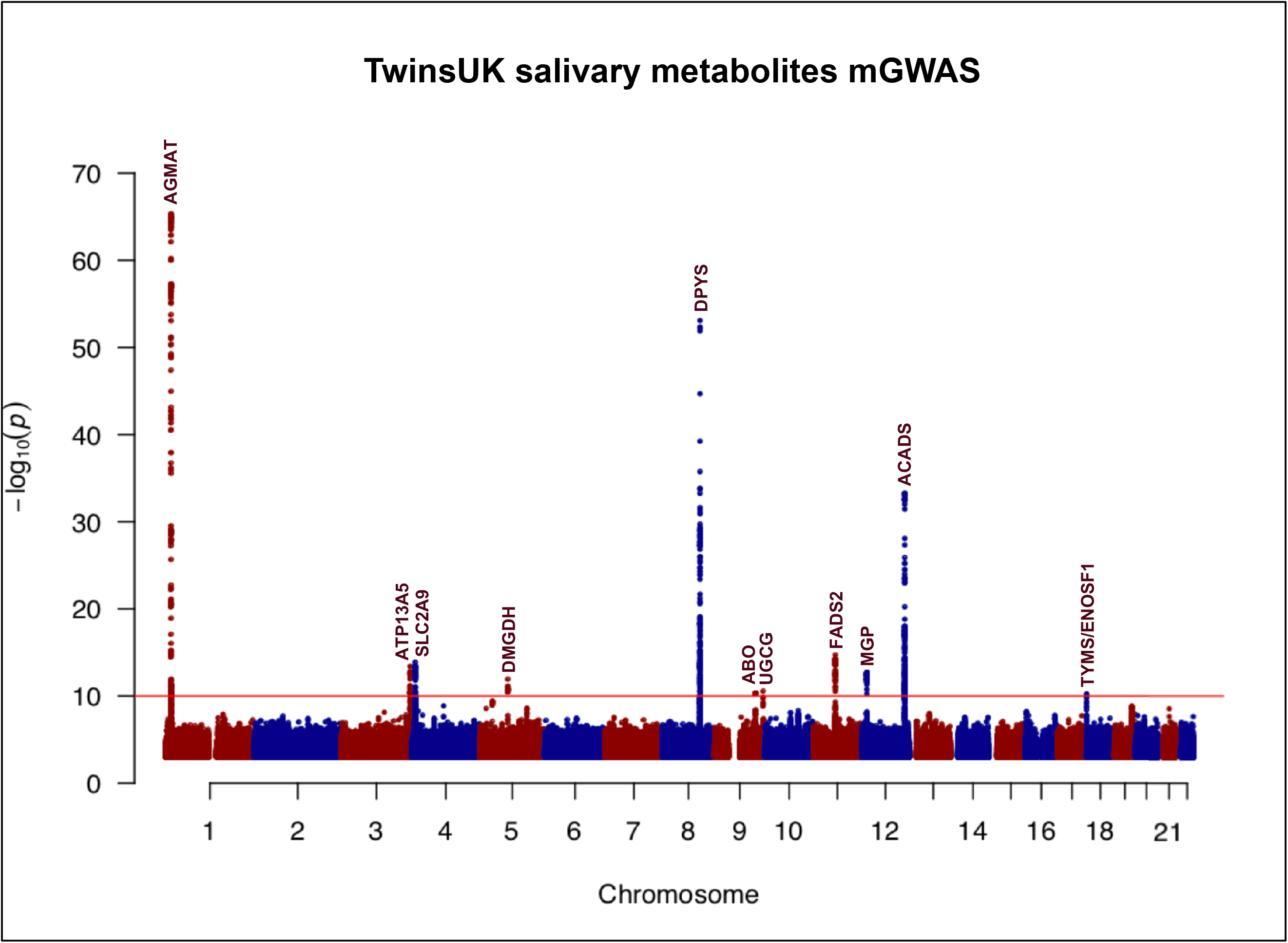
Manhattan plot illustrating the findings of the discovery phase (TwinsUK) of the genome-wide association study for salivary metabolites (mGWAS) The red horizontal line demarcates the study-wide significance threshold of 10^−10^. Eleven loci surpassed the study-wide significance threshold. The loci are referred to by the name(s) of the overlapping or the nearest gene(s).

**Table 1:** Summary of genetic loci that were significantly associated with salivary metabolite(s) in the discovery phase (TwinsUK)

| Locus | Index variant | Chr | Position (build 37 bp) | Biochemical (Metabolite) | EA | EAF | Beta | SE | P-value |
| --- | --- | --- | --- | --- | --- | --- | --- | --- | --- |
| AGMAT | rs10927806 | 1 | 15,909,239 | 4-guanidinobutanoate | C | 0.45 | 0.252 | 0.037 | 4.4 x 10<sup>-11</sup> |
|  | rs6690813 | 1 | 15,861,073 | beta-guanidinopropanoate | C | 0.45 | 0.708 | 0.034 | 4.6 x 10<sup>-66</sup> |
| ATP13A5 | rs55918334 | 3 | 193,086,314 | creatinine | G | 0.53 | 0.305 | 0.039 | 3.9 x 10<sup>-14</sup> |
| SLC2A9 | rs13129697 | 4 | 9,926,967 | urate | T | 0.72 | 0.337 | 0.042 | 1.3 x 10<sup>-14</sup> |
|  | *rs7675964 | 4 | 9,941,434 | allantoin | C | 0.73 | 0.239 | 0.043 | 4.1 x 10<sup>-8</sup> |
| DMGDH | rs248386 | 5 | 78,330,227 | dimethylglycine | C | 0.81 | -0.349 | 0.048 | 1.1 x 10<sup>-12</sup> |
| DPYS | rs80274300 | 8 | 105,470,882 | 3-ureidopropionate | C | 0.81 | -0.764 | 0.043 | 8.1 x 10<sup>-54</sup> |
|  | rs80274300 | 8 | 105,470,882 | 3-ureidoisobutyrate | C | 0.81 | -0.482 | 0.048 | 5.8 x 10<sup>-22</sup> |
| ABO | rs201298979 | 9 | 136,145,424 | N-acetylglucosamine/<br>N-acetylgalactosamine | AC | 0.77 | -0.314 | 0.046 | 2.6 x 10<sup>-11</sup> |
| UGCG | rs10981098 | 9 | 114,591,623 | glycosyl-N-stearoyl-sphinganine | T | 0.79 | -0.308 | 0.046 | 4.6 x 10<sup>-11</sup> |
| FADS2 | rs174564 | 11 | 61,588,305 | 1-(1-enyl-palmitoyl)-2-arachidonoyl-GPC | A | 0.67 | 0.324 | 0.039 | 1.9 x 10<sup>-15</sup> |
| ACADS | rs34673751 | 12 | 121,171,891 | ethylmalonate | G | 0.74 | -0.539 | 0.041 | 5.1 x 10<sup>-34</sup> |
| MGP | rs5796614 | 12 | 15,053,094 | gamma-carboxyglutamate | G | 0.6 | -0.29 | 0.038 | 1.9 x 10<sup>-13</sup> |
| TYMS/<br>ENOSF1 | rs2790 | 18 | 673,086 | ribonate | A | 0.81 | 0.325 | 0.049 | 5.8 x 10<sup>-11</sup> |

(Chr = Chromosome; EA = Effect Allele; EAF = Effect Allele Frequency; SE = Standard Error)
For the primary stage of association analysis in the discovery phase (TwinsUK), locus-metabolite associations (mQTLs) were identified using a genome-wide and metabolome-wide significance threshold of $p < 10^{-10}$ .
\*This mQTL was identified when the analysis was restricted to only the loci that were identified in the primary stage of association testing (significance threshold of $p < 10^{-6}$ )

Of the 11 genetic loci that were identified in the discovery phase, four of them *(SLC2A9, DMGDH, FADS2* and *ACADS)* have previously been reported in association with blood metabolites^2,4,5^; the remaining seven loci were novel i.e. there were no previously known associations for them with metabolites in blood or any other biospecimens.

The *AGMAT* locus, one of the novel loci identified, was associated with the metabolites 4-guanidinobutanoate and beta-guanidinopropanoate. These metabolites are generated as intermediate products in the polyamine synthesis pathway, the main site of action for the enzyme agmatinase (encoded by the *AGMAT* gene)^26^. The most significantly associated variants for the two metabolites i.e. rs10927806 and rs6690813, respectively, are in high LD with one another (r^2^=0.99), suggesting a shared underlying genetic regulation for the two metabolites by the *AGMAT* locus. Likewise, the association between the *ATP13A5* locus and creatinine (a widely used measure of renal function), observed in our study, has also not been reported previously. Another interesting novel association that we identified pertained to the metabolism of the pyrimidine uracil - the *DPYS* locus (encodes for the enzyme dihydropyrimidinase, involved in uracil degradation) was associated with the metabolites 3-ureidopropionate and 3-ureidoisobutyrate (breakdown products of uracil metabolism). The novel association for the *TYMS/ENOSF1* locus with ribonate is also intriguing, since it has previously been shown that ribonate is one of the substrates for the catalytic activity of reverse thymidylate synthase (rTS), the protein product of *ENOSF1^27^*. Therefore, it appears that *ENOSF1*, which is the source of anti-sense RNA of *TYMS*, is probably the functional gene mediating the observed association of the *TYMS/ENOSF1* locus with salivary ribonate. The associations of the *SLC2A9* locus with allantoin, the *ABO* locus with N-acetylglucosamine/N-acetylgalactosamine, the *UGCG* locus with glycosyl-N-stearoyl-sphinganine, and the *MGP* locus with gamma-carboxyglutamate were the remaining mQTLs observed in our study that have not been reported previously.

Of the 14 distinct mQTLs that were identified in our study, it was possible to test the most significant variant-metabolite pair for nine mQTLs in both serum and faecal metabolite data in TwinsUK (metabolites corresponding to the remaining five mQTLs were not present in the serum and faecal metabolite datasets). Of them, the associations for the *ATP13A5* and *DPYS* loci did not replicate in serum (p>0.05); while, none of the associations, barring the one for the *ACADS* locus, replicated in the faecal data (p>0.05) (**Table S1**). Thus, a comparison across all three biospecimens (for the significantly associated salivary metabolites which were also measured in serum and faecal samples in TwinsUK) demonstrates that the effects of the *ATP13A5* and *DPYS* loci were specific to saliva (**Figure S4**).

For none of the mQTLs that was identified, did we find any additional independent association signals after conditioning for the most significant variant at the locus (conditional p>10^−5^ for all variants tested at each locus), a finding which was verified by the regional association plots (**Figure S5**). The observation that a single genetic association signal underlies each of the significantly associated metabolites might partly be due to our lack of power to detect secondary signals at these loci.

Moreover, the strength of association for the most significant variant-metabolite pair did not change much on adjusting for PD status, for any of the mQTLs (**Table S2**). Hence, it does not appear that the presence of underlying PD has a significant effect on the associations observed in our study.

For 8 of the 11 associated loci, it was observed that the most significant variant demonstrated an eQTL effect on at least one transcript in one of the tissues in the GTEx database (no significant eQTL effects were observed for the *ATP13A5, SLCA2A9* and *ABO* loci) (**Table S3**). In case of 7 of those 8 loci (except *DPYS)*, the significant eQTL effect was observed for the overlapping or the nearest gene transcript; and for 5 of those 7 loci *(AGMAT, FADS2, DMGDH, TYMS/ENOSF1* and *UGCG)*, the eQTL effect was observed in one of the gut-related tissues. While eQTL data for transcripts assayed in the minor salivary glands was available for a small number of donors (N=97) in the GTEx database, none of the significant eQTL effects that we observed was in the salivary tissue.

### Replication of the discovery phase findings

In SHIP-2, we could attempt replication for 9 of the 14 mQTLs that were identified in the discovery phase (metabolites corresponding to the remaining five mQTLs were not measured in SHIP-2). In the initial replication analysis, which was performed using salivary metabolite data that was not normalised for sample osmolality, 8 of the 9 discovery phase associations were replicated (p<0.05), with the direction of effect consistent with that observed in TwinsUK (**Table 2**). The association between the *ABO* locus and N-acetylglucosamine/N-acetylgalactosamine was the only finding from the discovery phase that did not replicate in SHIP-2. When we repeated the replication analysis in SHIP-2 with metabolite data that was normalised for the sample osmolality, the strength of all the associations was comparatively reduced (**Table S4**).

**Table 2:** Summary of the discovery phase associations that were tested in the replication study (SHIP-2)

| Locus | Chr | Index variant | Biochemical (Metabolite) | EA | Beta | SE | P-value |
| --- | --- | --- | --- | --- | --- | --- | --- |
| AGMAT | 1 | rs10927806 | 4-guanidinobutanoate | C | 0.115 | 0.044 | $8.9 \times 10^{-3}$ |
| ATP13A5 | 3 | rs55918334 | creatinine | G | 0.239 | 0.04 | $2.8 \times 10^{-9}$ |
| SLC2A9 | 4 | rs13129697 | urate | T | 0.576 | 0.045 | $7.7 \times 10^{-35}$ |
| | 4 | rs7675964 | allantoin | C | 0.209 | 0.049 | $2.3 \times 10^{-5}$ |
| DMGDH | 5 | rs248386 | dimethylglycine | C | -0.336 | 0.055 | $1.4 \times 10^{-9}$ |
| DPYS | 8 | rs80274300 | 3-ureidopropionate | C | -1.035 | 0.049 | $3.1 \times 10^{-82}$ |
| ABO<sup>†</sup> | 9 | rs9411378 | N-acetylglucosamine/<br>N-acetylgalactosamine | A | 0.032 | 0.048 | 0.5 |
| ACADS | 12 | rs34673751 | ethylmalonate | G | -0.132 | 0.049 | $7.3 \times 10^{-3}$ |
| TYMS/<br>ENOSF1 | 18 | rs2790 | ribonate | A | 0.137 | 0.055 | 0.013 |

(Chr = Chromosome; EA = Effect Allele; SE = Standard Error)
<sup>†</sup>Since information for the index variant (rs201298979) at the *ABO* locus was not available in the replication study, a proxy variant (rs9411378) that was in high LD ( $r^2=0.95$ ) with the index variant was used for the replication analysis.
Replication in SHIP-2 could be attempted for 9 of the 14 mQTLs that were identified in the discovery phase (metabolites corresponding to the remaining five mQTLs were not measured in SHIP-2). For each of these nine mQTLs, the most significant variant-metabolite pair identified in the discovery phase was tested in SHIP-2 using salivary metabolite data that was not normalised for osmolality (since SHIP-2 saliva samples represented stimulated saliva).

### Phenotypic associations for salivary metabolites

Majority of the loci that were associated in our study have been reported in relation with GWAS traits, inborn errors of metabolism and / or clinically relevant biochemical pathways (**Table 3**). We further tested the salivary metabolites associated with the *DPYS, AGMAT* and *ATP13A5* loci in relation with specific phenotypes using information available in the TwinsUK database (**Table 4**).

**Table 3:**
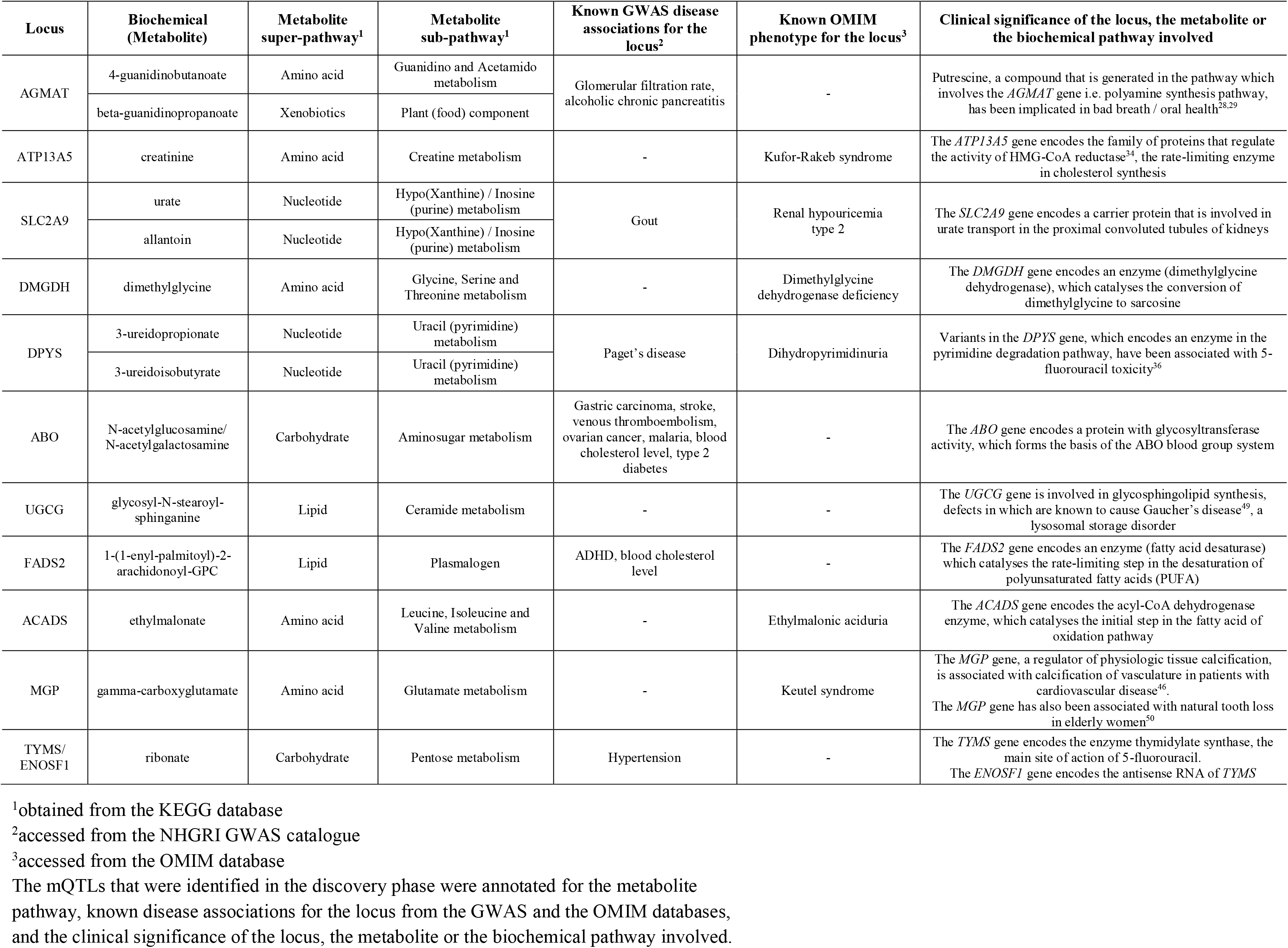
Annotation of the genetic loci and the corresponding metabolites that were significantly associated in the discovery phase.

**Table 4:** Summary of the phenotypes that were tested in relation with specific salivary metabolites.

| Locus | Chr | Biochemical (Metabolite) | Phenotypic association(s) tested |
| --- | --- | --- | --- |
| AGMAT | 1 | 4-guanidinobutanoate | 1. Periodontal disease<br>2. Clinical depression or anxiety disorder |
|  |  | beta-guanidinopropanoate |  |
| ATP13A5 | 3 | creatinine | 1. eGFR (measure of renal function)<br>2. Grip strength (measure of muscle strength)<br>3. Statin usage |
| DPYS | 8 | 3-ureidopropionate | 1. Irritable bowel syndrome (based on the ROME-III criteria<sup>36</sup>) |
|  |  | 3-ureidoisobutyrate |  |

(Chr = Chromosome; eGFR = effective Glomerular Filtration Rate)
The metabolites that were uniquely associated in saliva i.e. the ones for which a genetic association had not been previously reported in blood, were tested with phenotypes (diseases / traits / adverse drug effects) relating to the metabolite or its associated biochemical pathway, using information from the TwinsUK database.

The *AGMAT*-associated metabolites (4-guanidinobutanoate and beta-guanidinopropanoate) are generated in the polyamine synthesis pathway (https://www.genome.jp/kegg-bin/show_pathway?hsa00330). This pathway also produces the compound putrescine, which has been implicated in poor oral health and foul breath^28,29^. Consequently, we evaluated the significance of the *AGMAT*-associated metabolites in oral health by testing them with PD status - the levels of both 4-guanidinobutanoate and beta-guanidinopropanoate were significantly higher in PD cases compared to controls (p=0.0003 and p=0.0006, respectively). Since eGFR (estimated glomerular filtration rate) is calculated on the basis of serum creatinine, and these two commonly used measures of renal function are negatively correlated, we wanted to investigate the association between salivary creatinine and eGFR. We observed a similar strong negative relationship between salivary creatinine and eGFR (p=1.6 x 10^−11^), which was indicative of a homeostasis between creatinine concentrations in serum and saliva. Furthermore, creatinine is also known to be a marker of muscle mass and strength^30^. We, therefore, tested salivary creatinine in relation with grip strength (a measure of muscle strength), which suggested a positive correlation between them (p=0.01).

Apart from these findings, the remaining phenotypic associations that we tested were largely negative, as follows:

The enzyme agmatinase (encoded by the *AGMAT* gene), which acts on the substrate agmatine, has been implicated in the pathophysiology of mood disorders^31^. Moreover, studies have also proposed agmatine as a novel neuromodulator^32^. We, therefore, tested the *AGMAT*-associated metabolites in relation with a diagnosis of clinical depression or anxiety disorder, and responses (on a ordinal scale) to questions in the Hospital Anxiety and Depression Scale (HADS) questionnaire^33^. But, neither analysis showed any significant association (**Table S5**).

*ATP13A5*, the locus that was associated with salivary creatinine, belongs to the family of ATPases that regulate the activity of HMG-CoA reductase^34^, the main site of action of the cholesterol-lowering class of drugs called statins. Statins are known to cause muscle dysfunction (myopathy) in a small fraction of patients^35^. Given that *ATP13A5* and statins both act in the same biochemical pathway, we assessed whether salivary creatinine is also associated with statin usage, and could therefore be used as a biomarker for statin-induced myopathy. There was, however, no association between salivary creatinine and statin usage (p=0.22).

5-Fluorouracil (5-FU) is a pyrimidine analogue that is a commonly used anticancer drug. It is eliminated from the body via the pyrimidine degradation pathway, hence variants in genes coding for the pyrimidine degradation enzymes (for instance, *DPYS*) are known to be associated with the development of 5-FU toxicity^36^, which mainly manifests as gastrointestinal side effects. Since we could not directly assess the DPYS-associated metabolites (3-ureidopropionate and 3-ureidoisobutyrate) in relation to the gastrointestinal side effects of 5-FU toxicity, we tested the metabolites with a commonly observed phenotype of gastrointestinal dysfunction, irritable bowel syndrome or IBS (ascertained using the ROME-III questionnaire^37^). For both metabolites, we observed that the levels were not significantly different in IBS “cases” compared to “controls” (p>0.05, for both metabolites). However, this negative finding does not negate the possibility that these metabolites could be of clinical use in predicting 5-FU toxicity.

## Discussion

Here we report a genome-wide association analysis of 476 metabolites measured in saliva samples of healthy population-based studies of European descent. We identified a total of 11 distinct genetic loci that regulate the level of 14 salivary metabolites, of which three loci were associated with more than one metabolite each.

The fact that saliva is reflective of the concentration of biochemicals in blood forms the basis for certain clinical applications of saliva that others have proposed previously such as therapeutic monitoring of drugs^38^, cortisol measurement^39^ and renal function monitoring^40^. Using salivary metabolite data, we replicated the associations for certain well-established genetic loci that are known to regulate the level of blood metabolites. Thus, our findings add further credence to the notion that, as biofluids, a certain degree of homeostasis exists between blood and saliva.

Additionally, we identified some novel associations in our study, which have expanded our knowledge of genetic influences on human metabolites. In particular, the association between the *DPYS* locus and pyrimidine metabolites is intriguing because of its clinical relevance. Mutations in the pyrimidine catabolism genes such as *DPYD* (encodes dihydropyrimidine dehydrogenase) and *DPYS* (encodes dihydropyrimidinase) have been linked to inborn errors of metabolism^41,42^ [MIM:274270 and 222748, respectively] as well as development of severe toxicity to the chemotherapeutic agent 5-FU^36,43,44^. Studies have previously demonstrated the applicability of salivary measurement of certain pyrimidine pathway metabolites (uracil and dihydrouracil) for evaluating 5-FU toxicity due to deficient *DPYD* activity^45^. In our study, variants in the *DPYS* gene correlated with the levels of specific pyrimidine metabolites (3-ureidopropionate and 3-ureidoisobutyrate). Therefore, studies to explore the utility of salivary measurements of these metabolites as non-invasive tools for predicting 5-FU toxicity resulting from mutations that affect *DPYS* activity are warranted. While we could not test 3-ureidopropionate and 3-ureidoisobutyrate in relation to the gastrointestinal side effects of 5-FU toxicity, we did not find any association between these metabolites and a phenotype relevant to gut dysfunction (IBS phenotype). Similarly, the observed association between salivary ribonate and the *ENOSF1* locus, overexpression of which has been shown to cause resistance of tumour cell lines to 5-FU^27^, was further indicative of the possible role of saliva as a tool for drug monitoring. The other novel finding of note was the association between the *AGMAT* locus and the metabolites 4-guanidinobutanoate and beta-guanidinopropanoate. The *AGMAT*-associated metabolites are produced in the polyamine synthesis pathway, which also generates the compound putrescine that has been implicated in oral health. Moreover, in our study, the *AGMAT*-associated metabolites were correlated with PD status, a disease related to poor oral health. Together, these findings suggest that the *AGMAT*-associated metabolites might serve as potential biomarkers for oral health. On the other hand, though the agmatine pathway has been previously implicated in mood disorders, we did not find any association between the *AGMAT*-associated metabolites and either symptoms of or a prior diagnosis of clinical depression or anxiety disorder. Thus, evidence based on the *AGMAT*-associated metabolites does not lend support to the hypothesis regarding a potential link between pathways involved in maintaining oral health and regulation of mood^15,18^. There were a few other novel associations of interest in our study, such as the metabolite gamma-carboxyglutamate (associated with the *MGP* locus, which is known to cause abnormal vascular calcification in patients with cardiovascular disease^46^), for which we could not assess the clinical significance since specific phenotypic information was not available in sufficient numbers in those with salivary metabolite data.

Given that we identified some novel genetic associations for salivary metabolites i.e. genetic loci that had not been reported in association with blood metabolites, it is indicative of the presence of regulatory pathways that are specific to the salivary metabolome. For the novel genetic loci, in most cases, the overlapping or the nearest gene transcript was expressed in one or more gut-related tissues, including salivary glands. However, for these loci, we did not find much evidence for cis-eQTL effects specific to salivary or other gut tissues, whereby we could attribute their association with the respective salivary metabolite(s) to transcriptional regulation of overlapping / neighbouring genes. Interestingly, there is growing evidence to suggest that the human metabolome is a reflection of an interaction between the host and the gut microbiome^7,47,48^ – it will, therefore, be worth exploring whether the salivary metabolites that were associated in our study correlate with the composition of the salivary microbiome. Attempting to elucidate, in this manner, biological processes that might mediate the observed associations between genetic loci and salivary metabolites will help broaden our understanding of the underlying pathways.

In summary, our study has provided an initial map of genetic loci that regulate the salivary metabolome. Based on what has been observed for other complex human traits, future studies with larger sample sizes are expected to uncover additional genetic loci with much smaller effects on salivary metabolites. Furthermore, our study also offered insights into hitherto unknown biological pathways involved in maintaining the levels of salivary metabolites. While we did explore, to an extent, the clinical relevance of a few salivary metabolites of interest, a more comprehensive analysis with a wider range of phenotypic domains will help broaden our understanding of the relationship between the salivary metabolome and systemic health conditions.

## Supplemental Data

Supplemental data includes supplemental materials and methods, five supplemental tables and six supplemental figures.

## Author Contributions

Conceived and designed the experiments: C.J.S., C.M., T.D.S., G.K., R.P.M., M.V.M, J.R., U.V.

Performed the experiments: R.P.M.

Analysed the data: A.N., K.S., Y.K., P.M.W., R.C.E.B., T.K., M.P., S.W., M.M.

Wrote the manuscript: A.N., C.J.S., C.M., K.S., R.P.M.

All authors revised the manuscript

## Declaration of Interests

R.P.M. is an employee of Metabolon, Inc. and, as such, has affiliations with or financial involvement with Metabolon, Inc.

## Acknowledgements

C.J.S. acknowledges funding from the Chronic Disease Research Foundation and the Wellcome Trust (grant WT081878MA). K.S. was supported by the Biomedical Research Program at Weill Cornell Medicine in Qatar, a program funded by the Qatar Foundation.

TwinsUK is funded by the Wellcome Trust, Medical Research Council, European Union, the National Institute for Health Research (NIHR)-funded BioResource, Clinical Research Facility and Biomedical Research Centre based at Guy’s and St. Thomas’ NHS Foundation Trust in partnership with King’s College London.

The SHIP cohort study is part of the Community Medicine Research Net (http://www.medizin.uni-greifswald.de/cm) of the University of Greifswald, Germany, which is funded by the German Federal Ministry of Education and Research (BMBF, grant no. 01ZZ96030, 01ZZ0701); the Ministry for Cultural Affairs and the Ministry for Social Affairs of the Federal State of Mecklenburg-West Pomerania (SHIP; http://www.medizin.uni-greifswald.de/cm).

## Web Resources

1000 Genomes project: http://www.internationalgenome.org/

Metabolon: https://www.metabolon.com/

GEMMA: http://www.xzlab.org/software.html

KEGG: https://www.genome.jp/kegg/

GCTA: http://cnsgenomics.com/software/gcta/#COJO

GTEx: https://gtexportal.org/home/

NHGRI GWAS catalogue: https://www.ebi.ac.uk/gwas/

OMIM: http://omim.org/

HMDB: http://www.hmdb.ca/

LocusZoom: http://locuszoom.org/

---

**Figure S1: Comparison of the distribution of saliva sample osmolality in TwinsUK and SHIP-2**

The distribution of saliva sample osmolality observed in SHIP-2 was much narrower in comparison to TwinsUK, indicative of the fact that, in contrast to TwinsUK, the saliva samples in SHIP-2 represented stimulated saliva.

**Figure S2: Plots demonstrating the correlation between the pairs of metabolites that were associated with the same locus in the discovery phase**

The figure demonstrates the correlation between the pairs of metabolites which were regulated by the same locus, as follows (i-iii): beta-guanidinopropanoate and 4-guanidinobutanoate (*AGMAT*); urate and allantoin (*SLC2A9*); 3-ureidopropionate and 3-ureidoisobutyrate (*DPYS*).

**Figure S3: Quantile-quantile (QQ) plots of the genome-wide association analysis for the 14 salivary metabolites that were significantly associated in the discovery phase**

The figure demonstrates the QQ plots for the significantly associated salivary metabolites, as follows (i-xiv): 4-guanidinobutanoate; beta-guanidinopropanoate; creatinine; urate; allantoin; dimethylglycine; 3-ureidopropionate; 3-ureidoisobutyrate; N-acetylglucosamine/N-acetylgalactosamine; glycosyl-N-stearoyl-sphinganine; 1-(1-enyl-palmitoyl)-2-arachidonoyl-GPC; ethylmalonate; gamma-carboxyglutamte; ribonate.

**Figure S4: A comparison of the effect of the index variant (from the salivary mGWAS) using measurements of the corresponding metabolite in faecal, serum and saliva samples of the TwinsUK subjects**

The plots demonstrate a comparison of the effect of the index variant (from the salivary mGWAS) for the significantly associated salivary metabolites that were measured in all three biospecimens (faeces, serum and saliva), as follows (i-vii): 4-guanidinobutanoate; creatinine; urate; dimethylglycine; 3-ureidopropionate; ethylmalonate; ribonate.

**Figure S5: Regional association plots for the genetic loci that were significantly associated with a salivary metabolite in the discovery phase**

Regional association plots were created using the LocusZoom tool for the locus-metabolite associations (mQTLs) that were identified in the discovery phase, as follows (i-xiii): *AGMAT* and 4-guanidinobutanoate; *AGMAT* and beta-guanidinopropanoate; *ATP13A5* and creatinine; *SLC2A9* and urate; *SLC2A9* and allantoin; *DMGDH* and dimethylglycine; *DPYS* and 3-ureidopropinoate; *DPYS* and 3-ureidoisobutyrate; *ABO* and N-acetylglucosamine/N-acetylgalactosamine; *UGCG* and glycosyl-N-stearoyl-sphinganine; *FADS2* and 1-(1-enyl-palmitoyl)-2-arachidonoyl-GPC; *ACADS* and ethylmalonate; *TYMS/ENOSF1* and ribonate *The regional association plot for the *MGP* locus has not been included since LD information for variants at that locus was not well characterised in the 1000 Genomes dataset.

**Figure S6: Boxplots demonstrating the distribution of the significantly associated salivary metabolites, stratified by the genotypes of the corresponding index variant**

The 14 salivary metabolites that were identified in the discovery phase of our study mapped to 11 distinct genetic loci: of which, three loci (*AGMAT, SLC2A9* and *DPYS*) were associated with two metabolites each; and the remaining eight loci were each associated with one metabolite. The plots demonstrate the distribution of the significantly associated salivary metabolite with respect to the genotypes of the corresponding index variant (for each index variant, the number of observations for each genotype is provided), as follows (i-xiv): 4-guanidinobutanoate; beta-guanidinopropanoate; creatinine; urate; allantoin; dimethylglycine; 3-ureidopropionate; 3-ureidoisobutyrate; N-acetylglucosamine/N-acetylgalactosamine; glycosyl-N-stearoyl-sphinganine; 1-(1-enyl-palmitoyl)-2-arachidonoyl-GPC; ethylmalonate; gamma-carboxyglutamte; ribonate.

## References

1. DeBerardinis, R.J., and Thompson, C.B. (2012). Cellular metabolism and disease: what do metabolic outliers teach us? Cell 148, 1132–1144.

2. Illig, T., Gieger, C., Zhai, G., Romisch-Margl, W., Wang-Sattler, R., Prehn, C., Altmaier, E., Kastenmuller, G., Kato, B.S., Mewes, H.-W., et al. (2010). A genome-wide perspective of genetic variation in human metabolism. Nat. Genet. 42, 137–141.

3. Kettunen, J., Tukiainen, T., Sarin, A.-P., Ortega-Alonso, A., Tikkanen, E., Lyytikainen, L.-P., Kangas, A.J., Soininen, P., Wurtz, P., Silander, K., et al. (2012). Genome-wide association study identifies multiple loci influencing human serum metabolite levels. Nat. Genet. 44, 269–276.

4. Shin, S.-Y., Fauman, E.B., Petersen, A.-K., Krumsiek, J., Santos, R., Huang, J., Arnold, M., Erte, I., Forgetta, V., Yang, T.-P., et al. (2014). An atlas of genetic influences on human blood metabolites. Nat. Genet. 46, 543–550.

5. Long, T., Hicks, M., Yu, H.-C., Biggs, W.H., Kirkness, E.F., Menni, C., Zierer, J., Small, K.S., Mangino, M., Messier, H., et al. (2017). Whole-genome sequencing identifies common-to-rare variants associated with human blood metabolites. Nat. Genet. 49, 568–578.

6. Gieger, C., Geistlinger, L., Altmaier, E., Hrabe de Angelis, M., Kronenberg, F., Meitinger, T., Mewes, H.-W., Wichmann, H.-E., Weinberger, K.M., Adamski, J., et al. (2008). Genetics meets metabolomics: a genome-wide association study of metabolite profiles in human serum. PLoS Genet. 4, e1000282.

7. Zierer, J., Jackson, M.A., Kastenmuller, G., Mangino, M., Long, T., Telenti, A., Mohney, R.P., Small, K.S., Bell, J.T., Steves, C.J., et al. (2018). The fecal metabolome as a functional readout of the gut microbiome. Nat. Genet. 50, 790–795.

8. Suhre, K., Wallaschofski, H., Raffler, J., Friedrich, N., Haring, R., Michael, K., Wasner, C., Krebs, A., Kronenberg, F., Chang, D., et al. (2011). A genome-wide association study of metabolic traits in human urine. Nat. Genet. 43, 565–569.

9. Luykx, J.J., Bakker, S.C., Lentjes, E., Neeleman, M., Strengman, E., Mentink, L., DeYoung, J., de Jong, S., Sul, J.H., Eskin, E., et al. (2014). Genome-wide association study of monoamine metabolite levels in human cerebrospinal fluid. Mol. Psychiatry 19, 228–234.

10. Soini, H.A., Klouckova, I., Wiesler, D., Oberzaucher, E., Grammer, K., Dixon, S.J., Xu, Y., Brereton, R.G., Penn, D.J., and Novotny, M. V (2010). Analysis of volatile organic compounds in human saliva by a static sorptive extraction method and gas chromatography-mass spectrometry. J. Chem. Ecol. 36, 1035–1042.

11. Barnes, V.M., Kennedy, A.D., Panagakos, F., Devizio, W., Trivedi, H.M., Jonsson, T., Guo, L., Cervi, S., and Scannapieco, F.A. (2014). Global metabolomic analysis of human saliva and plasma from healthy and diabetic subjects, with and without periodontal disease. PLoS One 9, e105181.

12. Wishart, D.S., Feunang, Y.D., Marcu, A., Guo, A.C., Liang, K., Vazquez-Fresno, R., Sajed, T., Johnson, D., Li, C., Karu, N., et al. (2018). HMDB 4.0: the human metabolome database for 2018. Nucleic Acids Res. 46, D608–D617.

13. Spielmann, N., and Wong, D.T. (2011). Saliva: diagnostics and therapeutic perspectives. Oral Dis. 17, 345–354.

14. Chiappin, S., Antonelli, G., Gatti, R., and De Palo, E.F. (2007). Saliva specimen: a new laboratory tool for diagnostic and basic investigation. Clin. Chim. Acta. 383, 30–40.

15. Kisely, S., Baghaie, H., Lalloo, R., Siskind, D., and Johnson, N.W. (2015). A systematic review and meta-analysis of the association between poor oral health and severe mental illness. Psychosom. Med. 77, 83–92.

16. Kaufman, E., and Lamster, I.B. (2002). The diagnostic applications of saliva--a review. Crit. Rev. Oral Biol. Med. 13, 197–212.

17. de Oliveira, C., Watt, R., and Hamer, M. (2010). Toothbrushing, inflammation, and risk of cardiovascular disease: results from Scottish Health Survey. BMJ 340, c2451.

18. O’Neil, A., Berk, M., Venugopal, K., Kim, S.-W., Williams, L.J., and Jacka, F.N. (2014). The association between poor dental health and depression: findings from a large-scale, population-based study (the NHANES study). Gen. Hosp. Psychiatry 36, 266–270.

19. Noble, J.M., Borrell, L.N., Papapanou, P.N., Elkind, M.S. V, Scarmeas, N., and Wright, C.B. (2009). Periodontitis is associated with cognitive impairment among older adults: analysis of NHANES-III. J. Neurol. Neurosurg. Psychiatry 80, 1206–1211.

20. Moayyeri, A., Hammond, C.J., Valdes, A.M., and Spector, T.D. (2013). Cohort Profile: TwinsUK and healthy ageing twin study. Int. J. Epidemiol. 42, 76–85.

21. Auton, A., Brooks, L.D., Durbin, R.M., Garrison, E.P., Kang, H.M., Korbel, J.O., Marchini, J.L., McCarthy, S., McVean, G.A., and Abecasis, G.R. (2015). A global reference for human genetic variation. Nature 526, 68–74.

22. Kanehisa, M., Goto, S., Sato, Y., Furumichi, M., and Tanabe, M. (2012). KEGG for integration and interpretation of large-scale molecular data sets. Nucleic Acids Res. 40, D109–D114.

23. Zhou, X., and Stephens, M. (2012). Genome-wide efficient mixed-model analysis for association studies. Nat. Genet. 44, 821–824.

24. Yang, J., Ferreira, T., Morris, A.P., Medland, S.E., Madden, P.A.F., Heath, A.C., Martin, N.G., Montgomery, G.W., Weedon, M.N., Loos, R.J., et al. (2012). Conditional and joint multiple-SNP analysis of GWAS summary statistics identifies additional variants influencing complex traits. Nat. Genet. 44, 369–375, S1-S3.

25. Kurushima, Y., Tsai, P.-C., Castillo-Fernandez, J., Couto Alves, A., El-Sayed Moustafa, J.S., Le Roy, C., Spector, T.D., Ide, M., Hughes, F.J., Small, K.S., et al. (2019). Epigenetic findings in periodontitis in UK twins: a cross-sectional study. Clin. Epigenetics 11, 27.

26. Dudkowska, M., Lai, J., Gardini, G., Stachurska, A., Grzelakowska-Sztabert, B., Colombatto, S., and Manteuffel-Cymborowska, M. (2003). Agmatine modulates the in vivo biosynthesis and interconversion of polyamines and cell proliferation. Biochim. Biophys. Acta 1619, 159–166.

27. Wichelecki, D.J., Froese, D.S., Kopec, J., Muniz, J.R.C., Yue, W.W., and Gerlt, J.A. (2014). Enzymatic and structural characterization of rTSgamma provides insights into the function of rTSbeta. Biochemistry 53, 2732–2738.

28. Goldberg, S., Kozlovsky, A., Gordon, D., Gelernter, I., Sintov, A., and Rosenberg, M. (1994). Cadaverine as a putative component of oral malodor. J. Dent. Res. 73, 1168–1172.

29. Porter, S.R., and Scully, C. (2006). Oral malodour (halitosis). BMJ 333, 632–635.

30. Patel, S.S., Molnar, M.Z., Tayek, J.A., Ix, J.H., Noori, N., Benner, D., Heymsfield, S., Kopple, J.D., Kovesdy, C.P., and Kalantar-Zadeh, K. (2013). Serum creatinine as a marker of muscle mass in chronic kidney disease: results of a cross-sectional study and review of literature. J. Cachexia. Sarcopenia Muscle 4, 19–29.

31. Bernstein, H.-G., Stich, C., Jager, K., Dobrowolny, H., Wick, M., Steiner, J., Veh, R., Bogerts, B., and Laube, G. (2012). Agmatinase, an inactivator of the putative endogenous antidepressant agmatine, is strongly upregulated in hippocampal interneurons of subjects with mood disorders. Neuropharmacology 62, 237–246.

32. Regunathan, S. (2006). Agmatine: biological role and therapeutic potentials in morphine analgesia and dependence. AAPS J. 8, E479–E484.

33. Stern, A.F. (2014). The hospital anxiety and depression scale. Occup. Med. (Lond). 64, 393–394.

34. Roitelman, J., and Simoni, R.D. (1992). Distinct sterol and nonsterol signals for the regulated degradation of 3-hydroxy-3-methylglutaryl-CoA reductase. J. Biol. Chem. 267, 25264–25273.

35. Sathasivam, S., and Lecky, B. (2008). Statin induced myopathy. BMJ 337, a2286.

36. Akai, F., Hosono, H., Hirasawa, N., and Hiratsuka, M. (2015). Novel single nucleotide polymorphisms of the dihydropyrimidinase gene (DPYS) in Japanese individuals. Drug Metab. Pharmacokinet. 30, 127–129.

37. Longstreth, G.F., Thompson, W.G., Chey, W.D., Houghton, L.A., Mearin, F., and Spiller, R.C. (2006). Functional bowel disorders. Gastroenterology 130, 1480–1491.

38. Drobitch, R.K., and Svensson, C.K. (1992). Therapeutic drug monitoring in saliva. An update. Clin. Pharmacokinet. 23, 365–379.

39. Aardal-Eriksson, E., Karlberg, B.E., and Holm, A.C. (1998). Salivary cortisol--an alternative to serum cortisol determinations in dynamic function tests. Clin. Chem. Lab. Med. 36, 215–222.

40. Goll, R.D., and Mookerjee, B.K. (1978). Correlation of biochemical parameters in serum and saliva in chronic azotemic patients and patients on chronic hemodialysis. J. Dial. 2, 344–399.

41. Hamajima, N., Kouwaki, M., Vreken, P., Matsuda, K., Sumi, S., Imaeda, M., Ohba, S., Kidouchi, K., Nonaka, M., Sasaki, M., et al. (1998). Dihydropyrimidinase deficiency: structural organization, chromosomal localization, and mutation analysis of the human dihydropyrimidinase gene. Am. J. Hum. Genet. 63, 717–726.

42. Berger, R., Stoker-de Vries, S.A., Wadman, S.K., Duran, M., Beemer, F.A., de Bree, P.K., Weits-Binnerts, J.J., Penders, T.J., and van der Woude, J.K. (1984). Dihydropyrimidine dehydrogenase deficiency leading to thymine-uraciluria. An inborn error of pyrimidine metabolism. Clin. Chim. Acta. 141, 227–234.

43. Wei, X., McLeod, H.L., McMurrough, J., Gonzalez, F.J., and Fernandez-Salguero, P. (1996). Molecular basis of the human dihydropyrimidine dehydrogenase deficiency and 5-fluorouracil toxicity. J. Clin. Invest. 98, 610–615.

44. van Kuilenburg, A.B., Haasjes, J., Richel, D.J., Zoetekouw, L., Van Lenthe, H., De Abreu, R.A., Maring, J.G., Vreken, P., and van Gennip, A.H. (2000). Clinical implications of dihydropyrimidine dehydrogenase (DPD) deficiency in patients with severe 5-fluorouracil-associated toxicity: identification of new mutations in the DPD gene. Clin. Cancer Res. 6, 4705–4712.

45. Neto, O.V., Raymundo, S., Franzoi, M.A., do Carmo Artmann, A., Tegner, M., Muller, V.V., Hahn, R.Z., Alves, G.V., Schwartsmann, G., Linden, R., et al. (2018). DPD functional tests in plasma, fresh saliva and dried saliva samples as predictors of 5-fluorouracil exposure and occurrence of drug-related severe toxicity. Clin. Biochem. 56, 18–25.

46. Mizobuchi, M., Towler, D., and Slatopolsky, E. (2009). Vascular calcification: the killer of patients with chronic kidney disease. J. Am. Soc. Nephrol. 20, 1453–1464.

47. Chen, M.X., Wang, S.-Y., Kuo, C.-H., and Tsai, I.-L. (2019). Metabolome analysis for investigating host-gut microbiota interactions. J. Formos. Med. Assoc. 118 Suppl 1, S10–S22.

48. Peisl, B.Y.L., Schymanski, E.L., and Wilmes, P. (2018). Dark matter in host-microbiome metabolomics: Tackling the unknowns-A review. Anal. Chim. Acta 1037, 13–27.

49. Alfonso, P., Navascues, J., Navarro, S., Medina, P., Bolado-Carrancio, A., Andreu, V., Irun, P., Rodriguez-Rey, J.C., Pocovi, M., Espana, F., et al. (2013). Characterization of variants in the glucosylceramide synthase gene and their association with type 1 Gaucher disease severity. Hum. Mutat. 34, 1396–1403.

50. Hirano, H., Ezura, Y., Ishiyama, N., Yamaguchi, M., Nasu, I., Yoshida, H., Suzuki, T., Hosoi, T., and Emi, M. (2003). Association of natural tooth loss with genetic variation at the human matrix Gla protein locus in elderly women. J. Hum. Genet. 48, 288–292.

