## Supplementary material for "Genome-wide scan identifies novel genetic loci regulating salivary metabolite levels": Supllemental material

### Supplemental Materials and Methods

#### (I) Sample preparation for global metabolomics profiling (TwinsUK and SHIP-2)

Metabolic profiling was conducted by Metabolon, Inc., as previously described for various specimens, particularly blood and stool[1,2]. All samples were stored at -80°C until processed. Saliva samples were extracted at a constant volume. Briefly, recovery standards were added prior to the first step in the extraction process for quality control purposes. To remove protein, dissociate small molecules bound to protein or trapped in the precipitated protein matrix, and to recover chemically diverse metabolites, proteins were precipitated with methanol under vigorous shaking for 2 min (Glen Mills Genogrinder 2000) followed by centrifugation. The resulting extract was divided into fractions and vacuum dried. For each sample, dried extracts were dissolved in injection solvent containing eight or more injection standards at fixed concentrations, depending on the platform, to assure injection and chromatographic consistency. Each sample was analyzed by four ultra-high performance liquid chromatography-tandem mass spectrometry (UPLC-MS/MS) methods: 1) One aliquot was analyzed using acidic positive ion conditions, chromatographically optimized for more hydrophilic compounds. In this method, the extract was gradient eluted from a C18 column (Waters UPLC BEH C18-2.1x100 mm, 1.7 µm) using water and methanol, containing 0.05% perfluoropentanoic acid (PFPA) and 0.1% formic acid (FA). 2) A second aliquot was also analyzed using acidic positive ion conditions; however, it was chromatographically optimized for more hydrophobic compounds. In this method, the extract was gradient eluted from the same aforementioned C18 column using methanol, acetonitrile, water, 0.05% PFPA and 0.01% FA and was operated at an overall higher organic content. 3) A third aliquot was analyzed using basic negative ion optimized conditions using a separate dedicated C18 column. The basic extracts were gradient eluted from the column using methanol and water, however with 6.5mM Ammonium Bicarbonate at pH 8. 4) The fourth aliquot was analyzed via negative ionization following elution from a HILIC column (Waters UPLC BEH Amide 2.1x150 mm, 1.7 µm) using a gradient consisting of water and acetonitrile with 10mM Ammonium Formate, pH 10.8. 5) A fifth aliquot was reserved for backup.

Three types of controls were analyzed in concert with the experimental samples: samples generated from a pool of human saliva used in the study served as technical replicate throughout the data set; extracted water samples served as process blanks; and a cocktail of standards spiked into every analyzed sample allowed instrument performance monitoring. Instrument variability was determined by calculating the median relative standard deviation (RSD) for the standards that were added to each sample prior to injection into the mass spectrometers (median RSD was 4% in SHIP-2 and 6% in TwinsUK; n=30 standards). Overall process variability was determined by calculating the median RSD for all endogenous metabolites (i.e., non-instrument standards) present in 100% of the pooled human saliva samples (median RSD was 11% in SHIP-2 and 14% in TwinsUK; n=377 metabolites in SHIP-2 and 499 metabolites in TwinsUK). Experimental samples and controls were randomized across the platform run.

All methods utilized a Waters ACQUITY UPLC and a Thermo Scientific Q-Exactive high resolution/accurate mass spectrometer interfaced with a heated electrospray ionization (HESI-II) source and Orbitrap mass analyzer operated at 35,000 mass resolution. Instruments were tuned and calibrated for mass resolution and mass accuracy daily. The MS analysis alternated between MS and data-dependent MSn scans using dynamic exclusion. The scan range varied slighted between methods but covered 70-1,000 m/z.

#### **(II) Metabolite identification and quantification** (TwinsUK and SHIP-2)

Metabolites were identified by automated comparison of the ion features in the experimental samples to a reference library of chemical standard entries that included retention time, molecular weight (m/z), preferred adducts, and in-source fragments as well as associated MS spectra, and curated by visual inspection for quality control using software developed at Metabolon[3]. Identification of known chemical entities was based on comparison to Metabolon's spectral library of >4,500 purified chemical standards. Commercially available purified standard compounds have been acquired and registered into LIMS for distribution to the various UPLC-MS/MS platforms for determination of their detectable characteristics. Known metabolites reported in this study conformed to confidence Level 1 (the highest confidence level of identification) of the Metabolomics Standards Initiative[4,5], unless otherwise denoted with an asterisk. Additional mass spectral entries have been created for structurally unnamed biochemicals (>2,750 in the Metabolon library), which have been identified by virtue of their recurrent nature (both chromatographic and mass spectral). These compounds have the potential to be identified by future acquisition of a matching purified standard or by classical structural analysis and were included in this study.

Peaks were quantified using area-under-the-curve. Raw area counts for each metabolite in each sample were normalized to correct for variation resulting from instrument inter-day tuning differences by the median value for each run-day, therefore, setting the medians to 1.0 for each run. This preserved variation between samples but allowed metabolites of widely different raw peak areas to be compared on a similar graphical scale.

#### (III) Study of Health in Pomerania (SHIP) cohort

The Study of Health in Pomerania (SHIP) is a prospective longitudinal population-based cohort study in Western Pomerania assessing the prevalence and incidence of common diseases and their risk factors[6]. Participants aged 20 to 79 with German citizenship and principal residency in the study area were recruited from a random sample of residents living in the three local cities, 12 towns as well as 17 randomly selected smaller towns. Individuals were randomly selected stratified by age and sex in proportion to population size of the city, town or small towns, respectively. A total of 4,308 participants were recruited between 1997 and 2001 in the SHIP cohort. Individuals were invited to the SHIP study centre for a computer-assisted personal interviews and extensive physical examinations. Follow-up investigations are scheduled in five year intervals. For the present project, data from the second SHIP follow-up (SHIP-2, 2008-2012) were used. The SHIP samples were genotyped using the Genome-Wide Human SNP Array 6.0 (Affymetrix, Santa Clara, USA). Imputation was performed to the 1000 Genomes Phase 3 integrated v5 panel (2014) on the Michigan Imputation Server v1.0.3 (https://imputationserver.sph.umich.edu/index.html).

Collection of saliva samples was performed with the commercially available collection system Salivette® (Sarstedt, Nümbrecht, Germany), as described previously[7]. Study subjects chew on a plain cotton role for 1 minute to stimulate salivation. Afterwards roles with the absorbed saliva were placed into the Salivette® and spun down to extract the collected saliva.

### Supplemental Tables

| Locus | Index variant | Biochemical (Metabolite) | EA | Serum metabolite data (N=546) | | | Faecal metabolite data (N=280) | | |
|---|---|---|---|---|---|---|---|---|---|
| | | | | Beta | SE | P-value | Beta | SE | P-value |
| AGMAT | rs10927806 | 4-guanidinobutanoate | C | 0.378 | 0.05 | $2.7 \times 10^{-12}$ | -0.153 | 0.095 | 0.11 |
| | rs6690813 | beta-guanidinopropanoate | C | NA | NA | NA | 0.142 | 0.097 | 0.14 |
| ATP13A5 | rs55918334 | creatinine | G | 0.017 | 0.068 | 0.8 | 0.036 | 0.096 | 0.71 |
| SLC2A9 | rs13129697 | urate | T | 0.345 | 0.068 | $9.1 \times 10^{-7}$ | 0.006 | 0.1 | 0.95 |
| | rs7675964 | allantoin | C | 0.157 | 0.061 | 0.01 | NA | NA | NA |
| DMGDH | rs248386 | dimethylglycine | C | -0.434 | 0.073 | $1.6 \times 10^{-8}$ | 0.073 | 0.103 | 0.48 |
| DPYS | rs80274300 | 3-ureidopropionate | C | 0.064 | 0.075 | 0.39 | 0.059 | 0.104 | 0.56 |
| ABO | rs201298979 | N-acetylglucosamine/ N-acetylgalactosamine | AC | NA | NA | NA | -0.085 | 0.1 | 0.39 |
| FADS2 | rs174564 | 1-(1-enyl-palmitoyl)-2-arachidonoyl-GPC | A | 0.462 | 0.062 | $2.4 \times 10^{-12}$ | NA | NA | NA |
| ACADS | rs34673751 | ethylmalonate | G | -1.026 | 0.053 | $1.6 \times 10^{-42}$ | -0.23 | 0.114 | 0.04 |
| TYMS/ ENOSF1 | rs2790 | ribonate | A | 0.292 | 0.077 | $1.8 \times 10^{-4}$ | 0.079 | 0.108 | 0.46 |

**Table S1: Association statistics observed in the serum and the faecal metabolite datasets for the significant mQTLs that were identified using salivary metabolite data in the discovery phase**

(EA = Effect Allele; SE = Standard Error; NA = Not available)

The significant mQTLs that were identified using salivary metabolite data were tested using measurements for the respective metabolite in serum and faecal samples of the TwinsUK subjects. Of the 14 distinct mQTLs that were identified using salivary metabolite data, it was possible to test nine of those associations in both serum and faecal metabolite data in the TwinsUK (the remaining metabolites were not present in the serum and faecal metabolite datasets).

| Locus | Index variant | Biochemical (Metabolite) | EA | Initial (unadjusted) analysis | | | On adjustment for periodontal disease status | | |
|---|---|---|---|---|---|---|---|---|---|
| | | | | Beta | SE | P-value | $Beta_{adj}$ | $SE_{adj}$ | $P\text{-value}_{adj}$ |
| AGMAT | rs10927806 | 4-guanidinobutanoate | C | 0.252 | 0.037 | $4.4 \times 10^{-11}$ | 0.245 | 0.038 | $4.7 \times 10^{-10}$ |
| | rs6690813 | beta-guanidinopropanoate | C | 0.708 | 0.034 | $4.6 \times 10^{-66}$ | 0.697 | 0.036 | $2 \times 10^{-59}$ |
| ATP13A5 | rs55918334 | creatinine | G | 0.305 | 0.039 | $3.9 \times 10^{-14}$ | 0.316 | 0.041 | $5.5 \times 10^{-14}$ |
| SLC2A9 | rs13129697 | urate | T | 0.337 | 0.042 | $1.3 \times 10^{-14}$ | 0.358 | 0.045 | $1.1 \times 10^{-14}$ |
| | rs7675964 | allantoin | C | 0.239 | 0.043 | $4.1 \times 10^{-8}$ | 0.250 | 0.050 | $4.2 \times 10^{-8}$ |
| DMGDH | rs248386 | dimethylglycine | C | -0.349 | 0.048 | $1.1 \times 10^{-12}$ | -0.351 | 0.050 | $7.4 \times 10^{-12}$ |
| DPYS | rs80274300 | 3-ureidopropionate | C | -0.764 | 0.043 | $8.1 \times 10^{-54}$ | -0.771 | 0.046 | $5.6 \times 10^{-48}$ |
| | rs80274300 | 3-ureidoisobutyrate | C | -0.482 | 0.048 | $5.8 \times 10^{-22}$ | -0.489 | 0.050 | $1.5 \times 10^{-20}$ |
| ABO | rs201298979 | N-acetylglucosamine/ N-acetylgalactosamine | AC | -0.314 | 0.046 | $2.6 \times 10^{-11}$ | -0.336 | 0.048 | $9.1 \times 10^{-12}$ |
| UGCG | rs10981098 | glycosyl-N-stearoyl-sphinganine | T | -0.308 | 0.046 | $4.6 \times 10^{-11}$ | -0.315 | 0.048 | $1.6 \times 10^{-10}$ |
| FADS2 | rs174564 | 1-(1-enyl-palmitoyl)-2-arachidonoyl-GPC | A | 0.324 | 0.039 | $1.9 \times 10^{-15}$ | 0.350 | 0.042 | $7.2 \times 10^{-16}$ |
| ACADS | rs34673751 | ethylmalonate | G | -0.539 | 0.041 | $5.1 \times 10^{-34}$ | -0.549 | 0.042 | $2.3 \times 10^{-32}$ |
| MGP | rs5796614 | gamma-carboxyglutamate | G | -0.290 | 0.038 | $1.9 \times 10^{-13}$ | -0.291 | 0.040 | $1.8 \times 10^{-12}$ |
| TYMS/ ENOSF1 | rs2790 | ribonate | A | 0.325 | 0.049 | $5.8 \times 10^{-11}$ | 0.363 | 0.051 | $3.3 \times 10^{-12}$ |

**Table S2: Summary of the effect of adjustment for periodontal disease status on the significant mQTLs that were identified in the discovery phase**

(EA = Effect Allele; SE = Standard Error; *adj* refers to the association statistics obtained after adjustment for periodontal disease status)

The significant mQTLs that were identified in the discovery phase were adjusted for periodontal disease status (self-reported gingival bleeding, and history of gum disease or tooth mobility were used as indicators of periodontal disease) in order to test the effect of the presence of underlying periodontal disease on the observed associations.

| Locus | Chr | Index variant | Biochemical (Metabolite) | EA | Salivary metabolite data without normalisation for sample osmolality | | | Salivary metabolite data after normalisation for sample osmolality | | |
|---|---|---|---|---|---|---|---|---|---|---|
| | | | | | Beta | SE | P-value | Beta | SE | P-value |
| AGMAT | 1 | rs10927806 | 4-guanidinobutanoate | C | 0.115 | 0.044 | $8.9 \times 10^{-3}$ | 0.053 | 0.045 | 0.23 |
| ATP13A5 | 3 | rs55918334 | creatinine | G | 0.239 | 0.04 | $2.8 \times 10^{-9}$ | 0.08 | 0.044 | 0.06 |
| SLC2A9 | 4 | rs13129697 | urate | T | 0.576 | 0.045 | $7.7 \times 10^{-35}$ | 0.243 | 0.05 | $1.2 \times 10^{-6}$ |
| | 4 | rs7675964 | allantoin | C | 0.209 | 0.049 | $2.3 \times 10^{-5}$ | 0.135 | 0.05 | $6.7 \times 10^{-3}$ |
| DMGDH | 5 | rs248386 | dimethylglycine | C | -0.336 | 0.055 | $1.4 \times 10^{-9}$ | -0.206 | 0.057 | $2.9 \times 10^{-4}$ |
| DPYS | 8 | rs80274300 | 3-ureidopropionate | C | -1.035 | 0.049 | $3.1 \times 10^{-82}$ | -0.849 | 0.052 | $1.4 \times 10^{-52}$ |
| ABO¶ | 9 | rs9411378 | N-acetylglucosamine/ N-acetylgalactosamine | A | 0.032 | 0.048 | 0.5 | -0.008 | 0.048 | 0.87 |
| ACADS | 12 | rs34673751 | ethylmalonate | G | -0.132 | 0.049 | $7.3 \times 10^{-3}$ | -0.064 | 0.05 | 0.2 |
| TYMS/ ENOSF1 | 18 | rs2790 | ribonate | A | 0.137 | 0.055 | 0.013 | -0.005 | 0.055 | 0.92 |

**Table S4: Comparison of findings in the replication study (SHIP-2) with and without normalisation of the salivary metabolite data for sample osmolality**

(Chr = Chromosome; EA = Effect Allele; SE = Standard Error)

¶Since information for the index variant (rs201298979) at the *ABO* locus was not available in the replication study, a proxy variant (rs9411378) that was in high LD ($r^2$=0.95) with the index variant was used for the replication analysis.

The initial replication analysis in SHIP-2, which was attempted for 9 of the 14 mQTLs that were identified in the discovery phase, was performed using salivary metabolite data that was not normalised for sample osmolality (since SHIP-2 saliva samples represented stimulated saliva). Next, the replication analysis in SHIP-2 was repeated with salivary metabolite data that was normalised for sample osmolality.

| HADS question (related to anxiety or depression) | Association p-value with AGMAT-associated metabolite | |
|---|---|---|
| | 4-guanidinobutanoate | beta-guanidinopropanoate |
| I feel tense or wound up | 0.03 | 0.55 |
| I still enjoy the things I used to enjoy | 0.25 | 0.4 |
| I get a sort of frightened feeling as if something awful is about to happen | 0.04 | 0.29 |
| I can laugh and see the funny side of things | 0.4 | 0.71 |
| Worrying thoughts go through my mind | 0.12 | 0.03 |
| I feel cheerful | 0.81 | 0.55 |
| I can sit at ease and feel relaxed | 0.28 | 0.47 |
| I feel as if I am slowed down | 0.19 | 0.98 |
| I get a sort of frightened feeling like butterflies in my stomach | 0.26 | 0.02 |
| I have lost interest in my appearance | 0.24 | 0.38 |
| I feel restless as if I have to be on the move | 0.38 | 0.5 |
| I look forward with enjoyment to things | 0.23 | 0.81 |
| I get sudden feelings of panic | 0.02 | 0.02 |
| I can enjoy a good book or radio, TV programme | 0.18 | 0.28 |

**Table S5: Summary of association between the AGMAT-associated metabolites (4-guanidinobutanoate and beta-guanidinopropanoate) and responses to questions (related to anxiety or depression) in the HADS (Hospital Anxiety and Depression Scale) questionnaire**

The agmatine pathway, which involves the enzyme agmatinase (encoded by the *AGMAT* gene), has been implicated in mood disorders. We performed an association analysis between the AGMAT-associated metabolites (4-guanidinobutanoate and beta-guanidinopropanoate) and responses to questions in the HADS questionnaire in order to test whether the metabolite levels correlated with specific symptoms related to anxiety or depression.

### Supplemental References

1. Menni, C., Kastenmuller, G., Petersen, A.K., Bell, J.T., Psatha, M., Tsai, P.-C., Gieger, C., Schulz, H., Erte, I., John, S., et al. (2013). Metabolomic markers reveal novel pathways of ageing and early development in human populations. Int. J. Epidemiol. *42*, 1111–1119.

2. Zierer, J., Jackson, M.A., Kastenmuller, G., Mangino, M., Long, T., Telenti, A., Mohney, R.P., Small, K.S., Bell, J.T., Steves, C.J., et al. (2018). The fecal metabolome as a functional readout of the gut microbiome. Nat. Genet. *50*, 790–795.

3. Dehaven, C.D., Evans, A.M., Dai, H., and Lawton, K.A. (2010). Organization of GC/MS and LC/MS metabolomics data into chemical libraries. J. Cheminform. *2*, 9.

4. Sumner, L.W., Amberg, A., Barrett, D., Beale, M.H., Beger, R., Daykin, C.A., Fan, T.W.-M., Fiehn, O., Goodacre, R., Griffin, J.L., et al. (2007). Proposed minimum reporting standards for chemical analysis Chemical Analysis Working Group (CAWG) Metabolomics Standards Initiative (MSI). Metabolomics *3*, 211–221.

5. Schrimpe-Rutledge, A.C., Codreanu, S.G., Sherrod, S.D., and McLean, J.A. (2016). Untargeted Metabolomics Strategies-Challenges and Emerging Directions. J. Am. Soc. Mass Spectrom. *27*, 1897–1905.

6. Volzke, H., Alte, D., Schmidt, C.O., Radke, D., Lorbeer, R., Friedrich, N., Aumann, N., Lau, K., Piontek, M., Born, G., et al. (2011). Cohort profile: the study of health in Pomerania. Int. J. Epidemiol. *40*, 294–307.

7. Salazar, M.G., Jehmlich, N., Murr, A., Dhople, V.M., Holtfreter, B., Hammer, E., Volker, U., and Kocher, T. (2013). Identification of periodontitis associated changes in the proteome of whole human saliva by mass spectrometric analysis. J. Clin. Periodontol. *40*, 825–832.

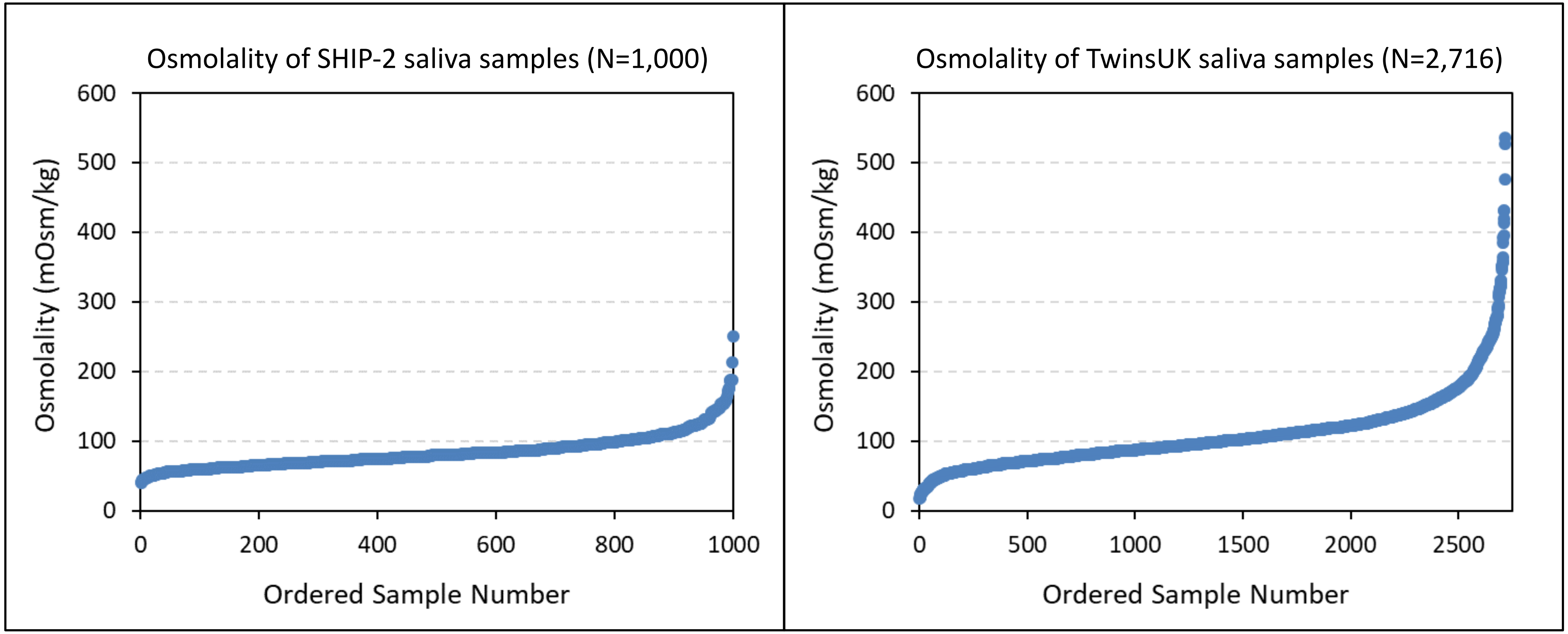

Osmolality of SHIP-2 saliva samples (N=1,000)
Osmolality (mOsm/kg)
600
500
400
300
200
100
0
0
200
400
600
800
1000
Ordered Sample Number
Osmolality of TwinsUK saliva samples (N=2,716)
Osmolality (mOsm/kg)
600
500
400
300
200
100
0
0
500
1000
1500
2000
2500
Ordered Sample Number

(i) beta-guanidinopropanoate and 4-guanidinobutanoate

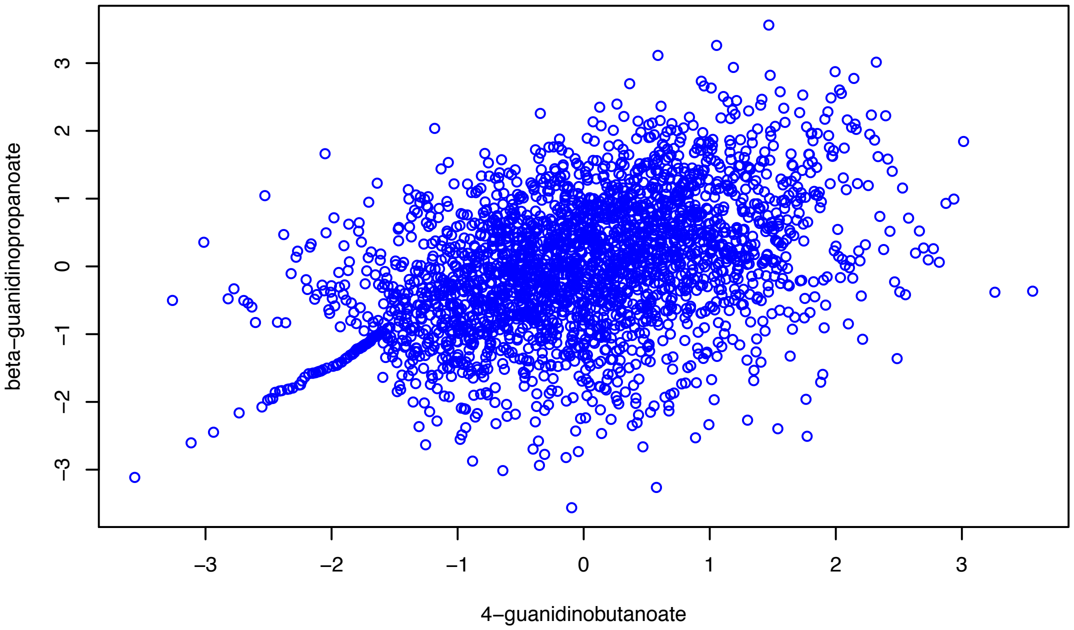


(ii) urate and allantoin

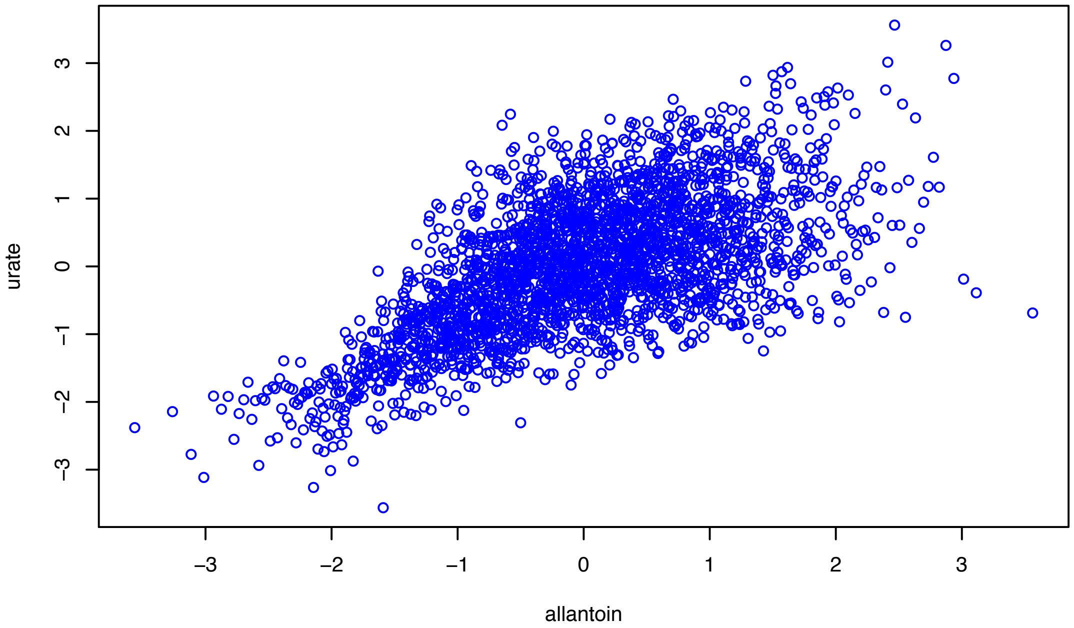


(iii) 3-ureidopropionate and 3-ureidoisobutyrate

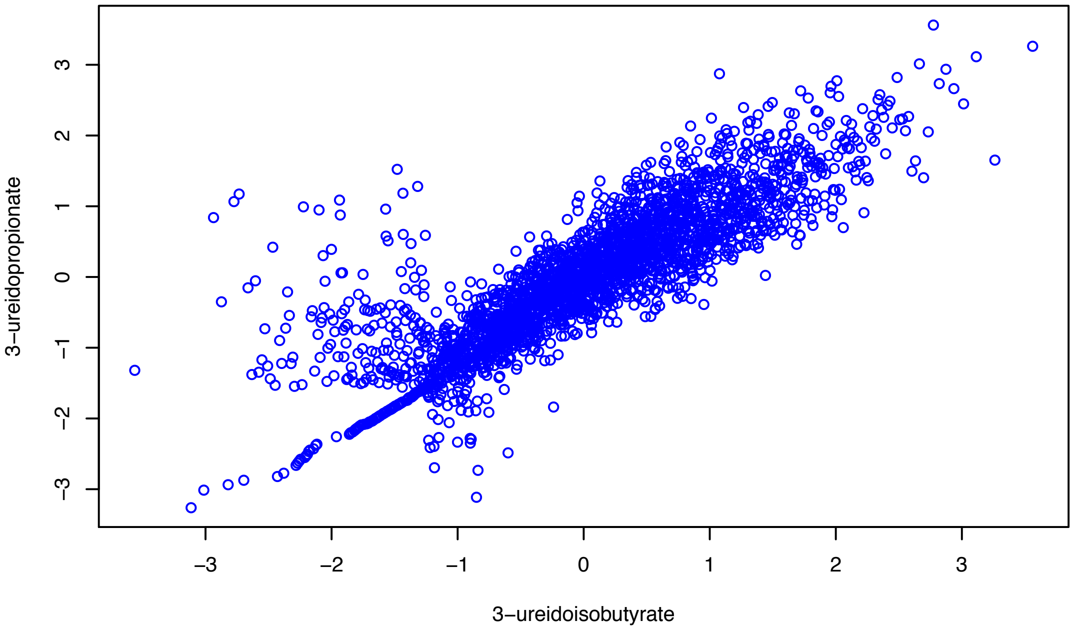

#### (i) 4-guanidinobutanoate

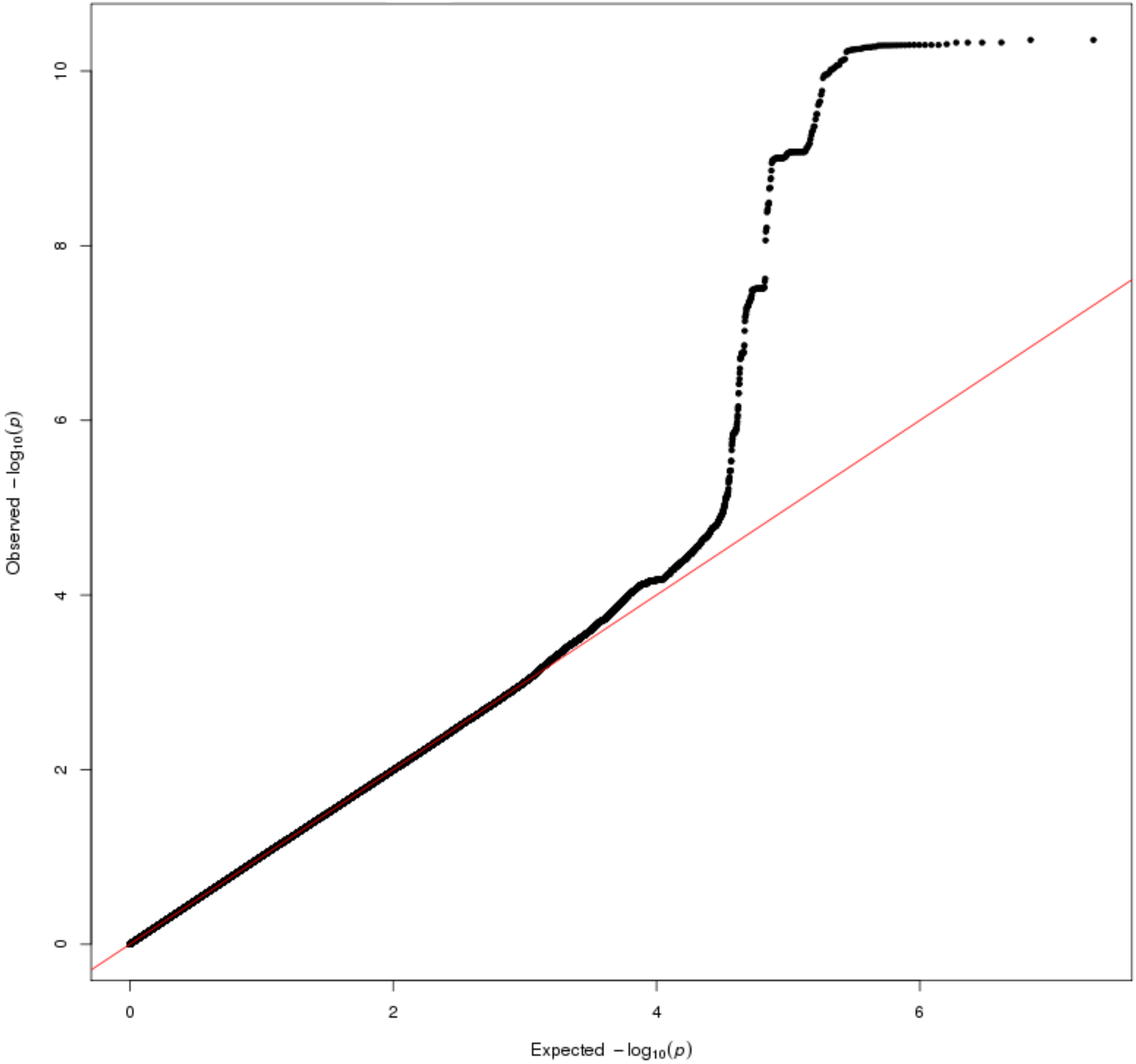

**(ii) beta-guanidinopropanoate**

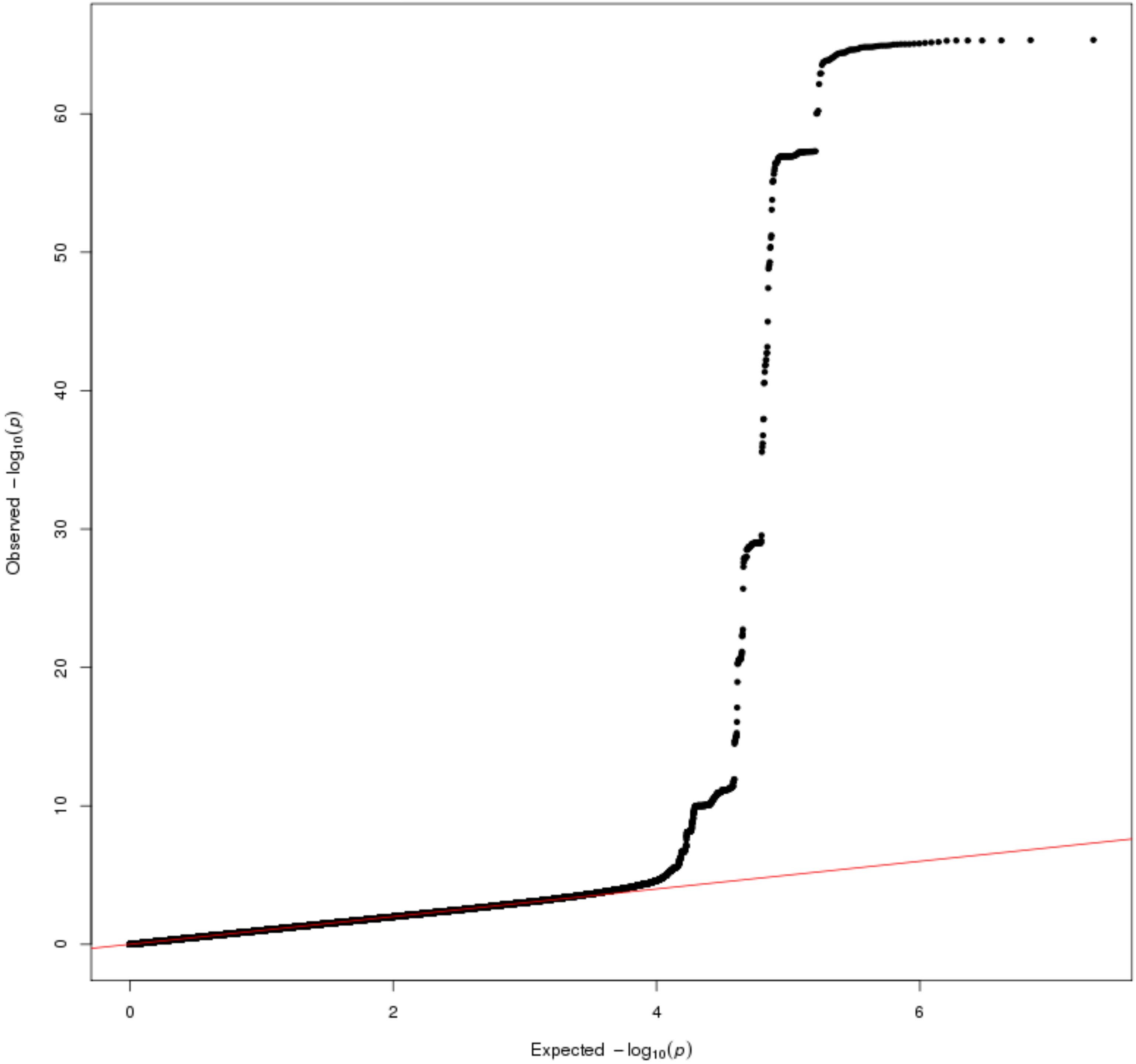

#### (iii) creatinine

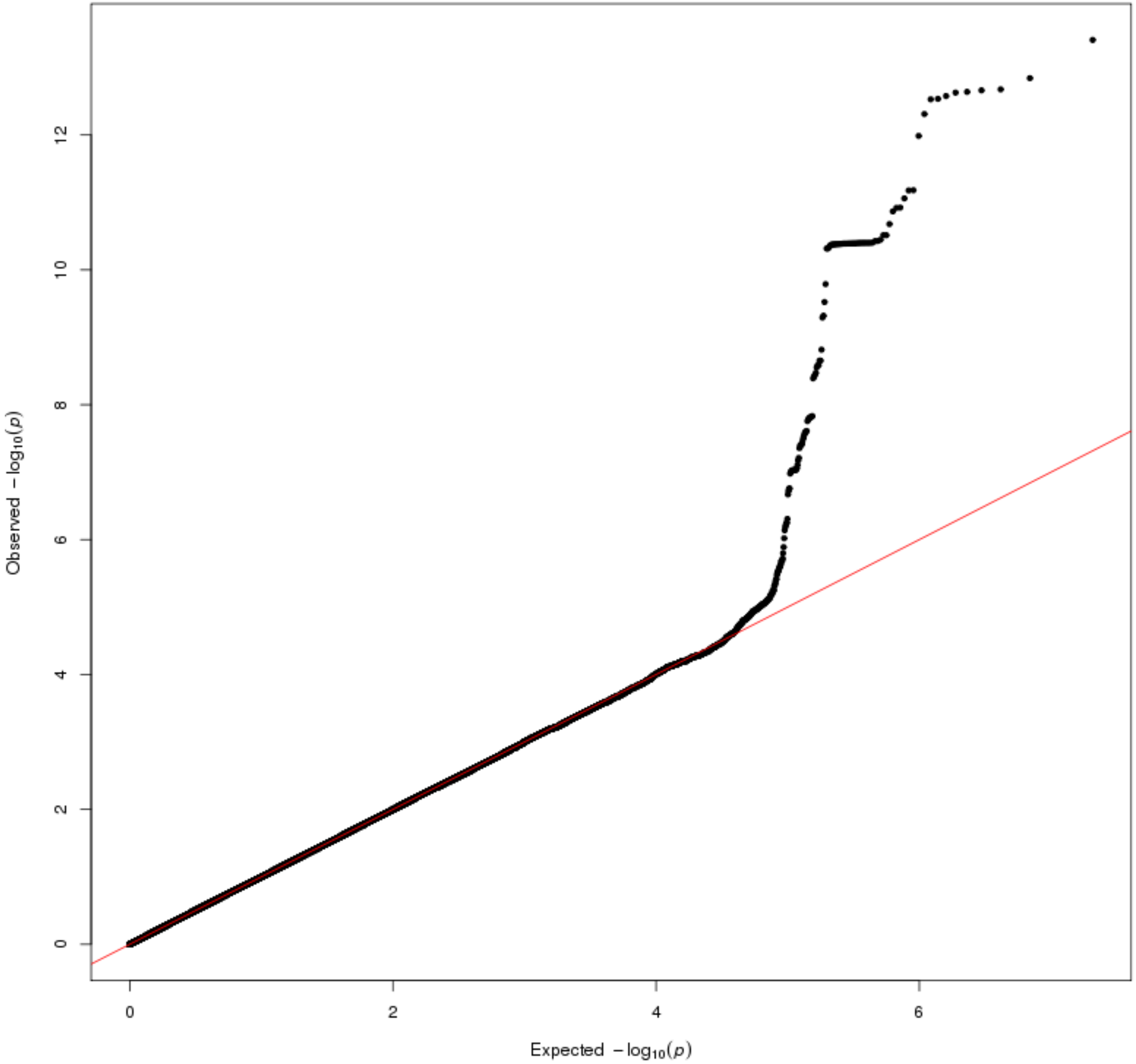

#### (iv) urate

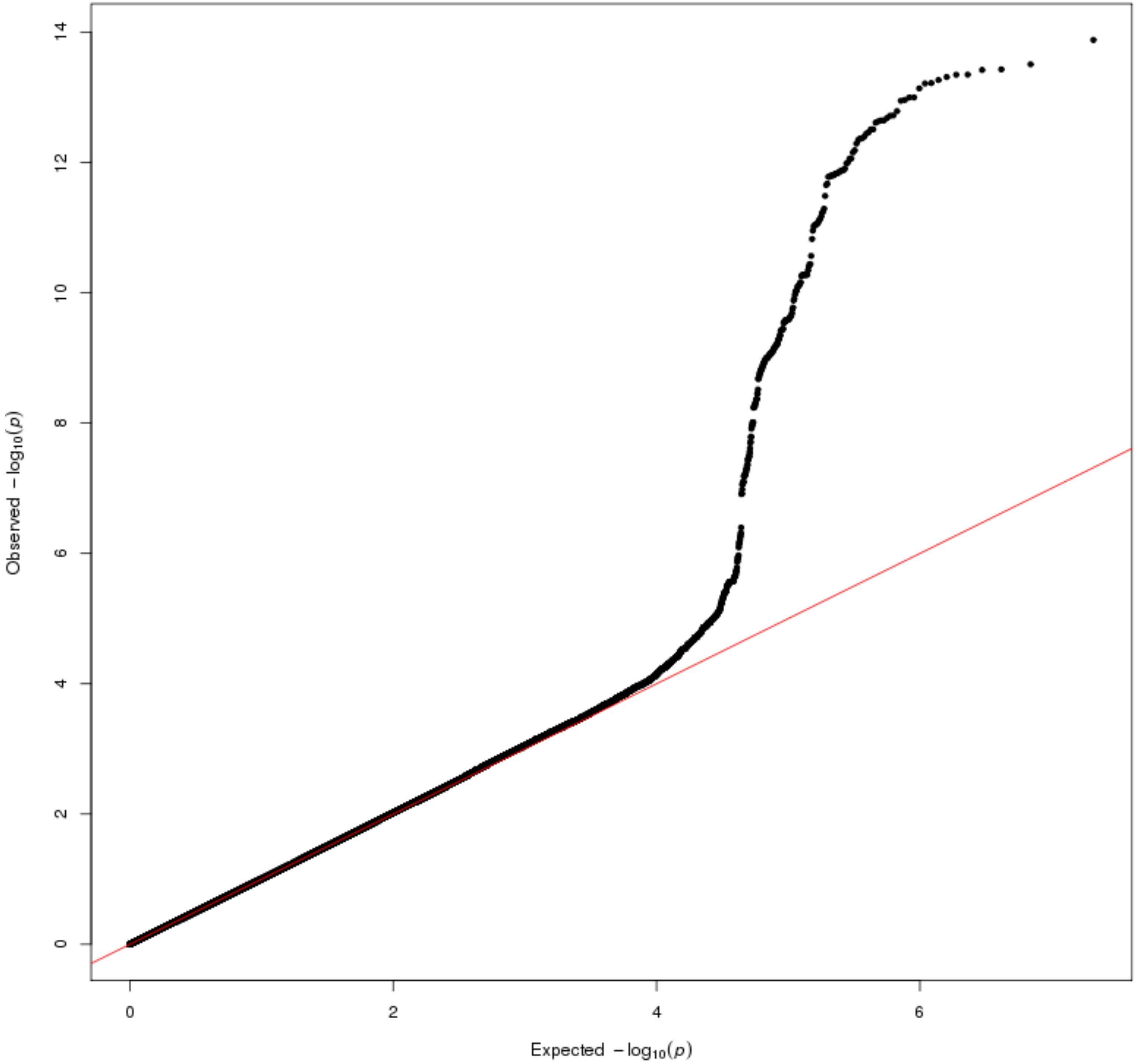

#### (v) allantoin

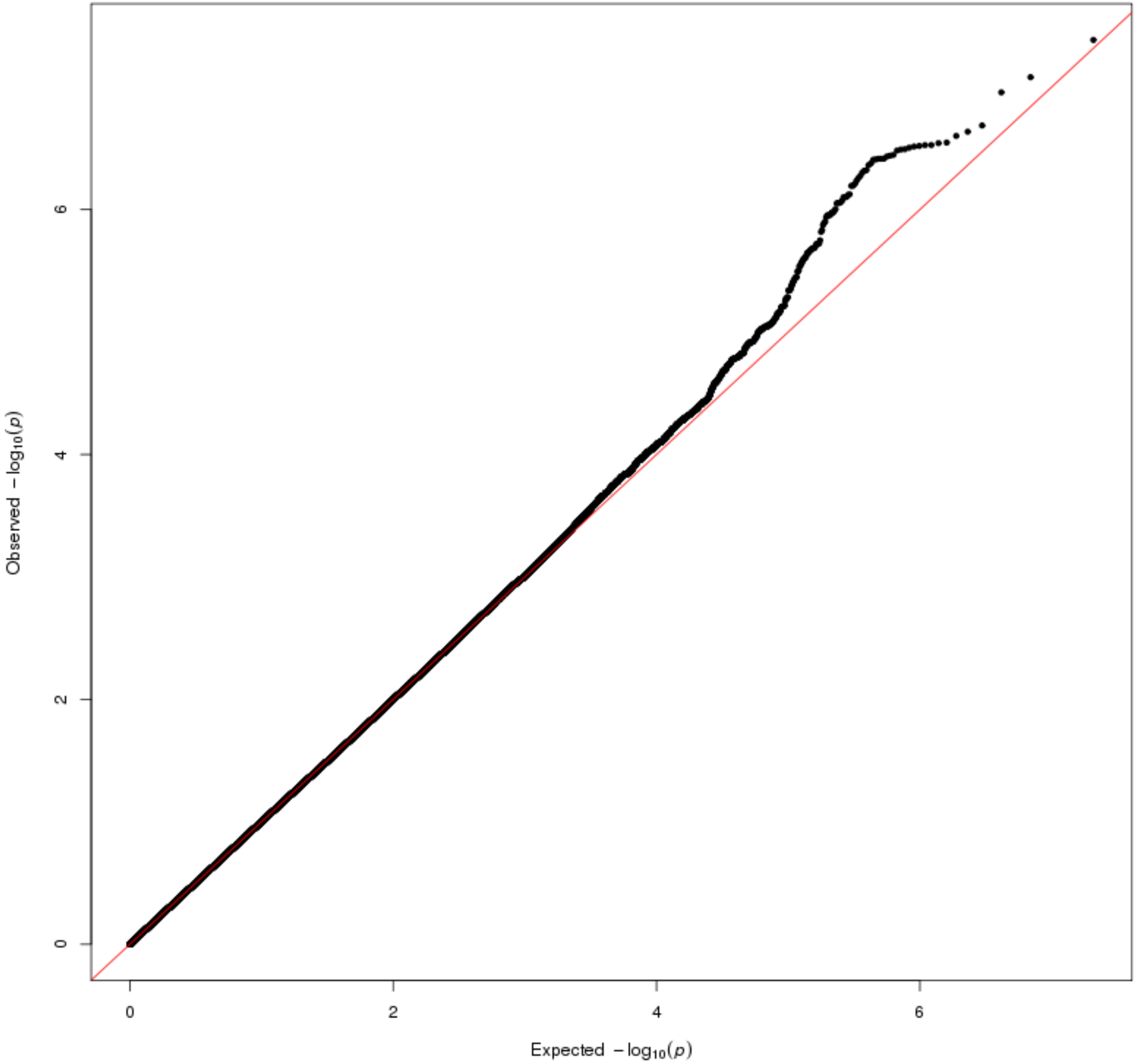

#### (vi) dimethylglycine

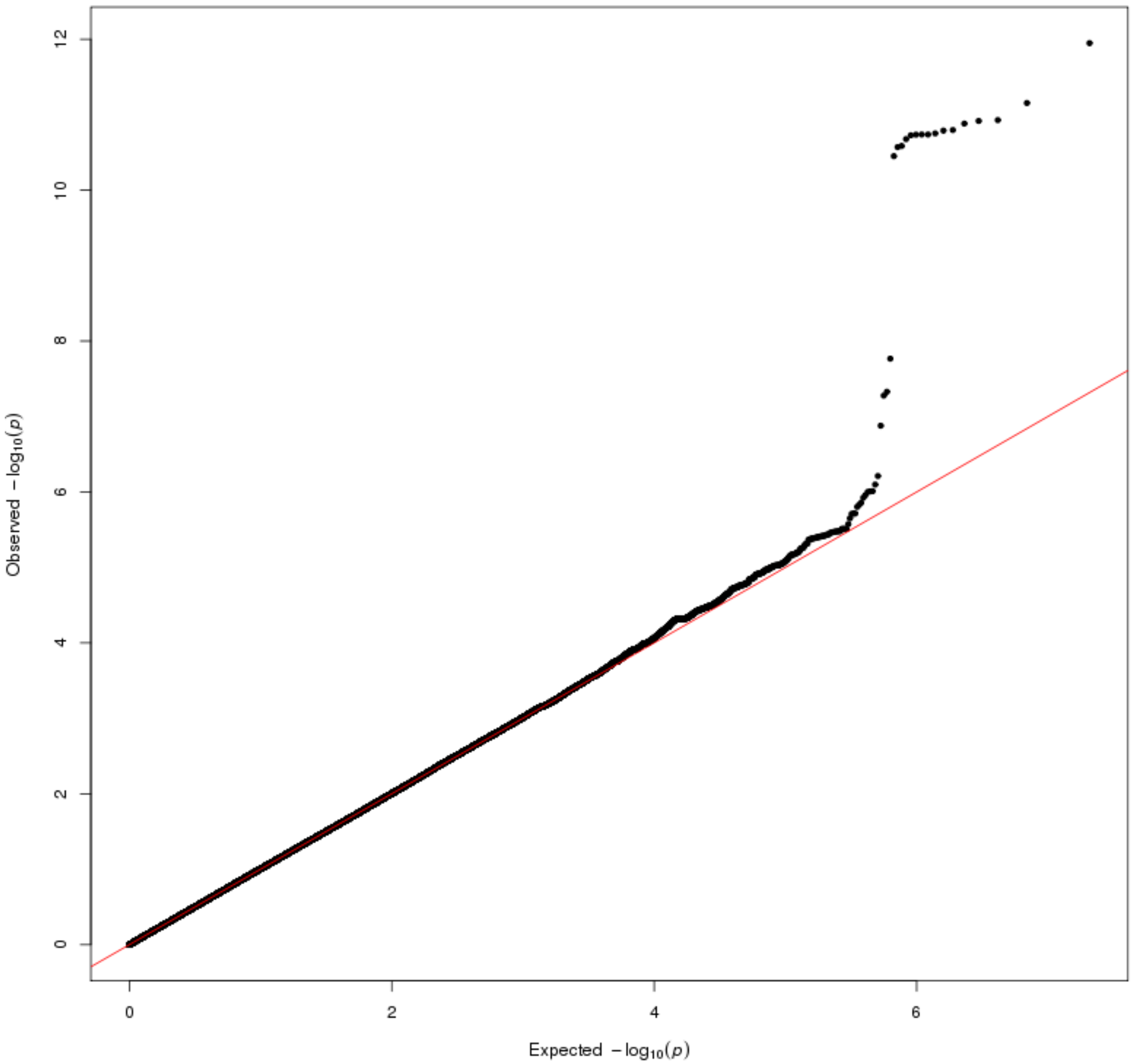

#### (vii) 3-ureidopropionate

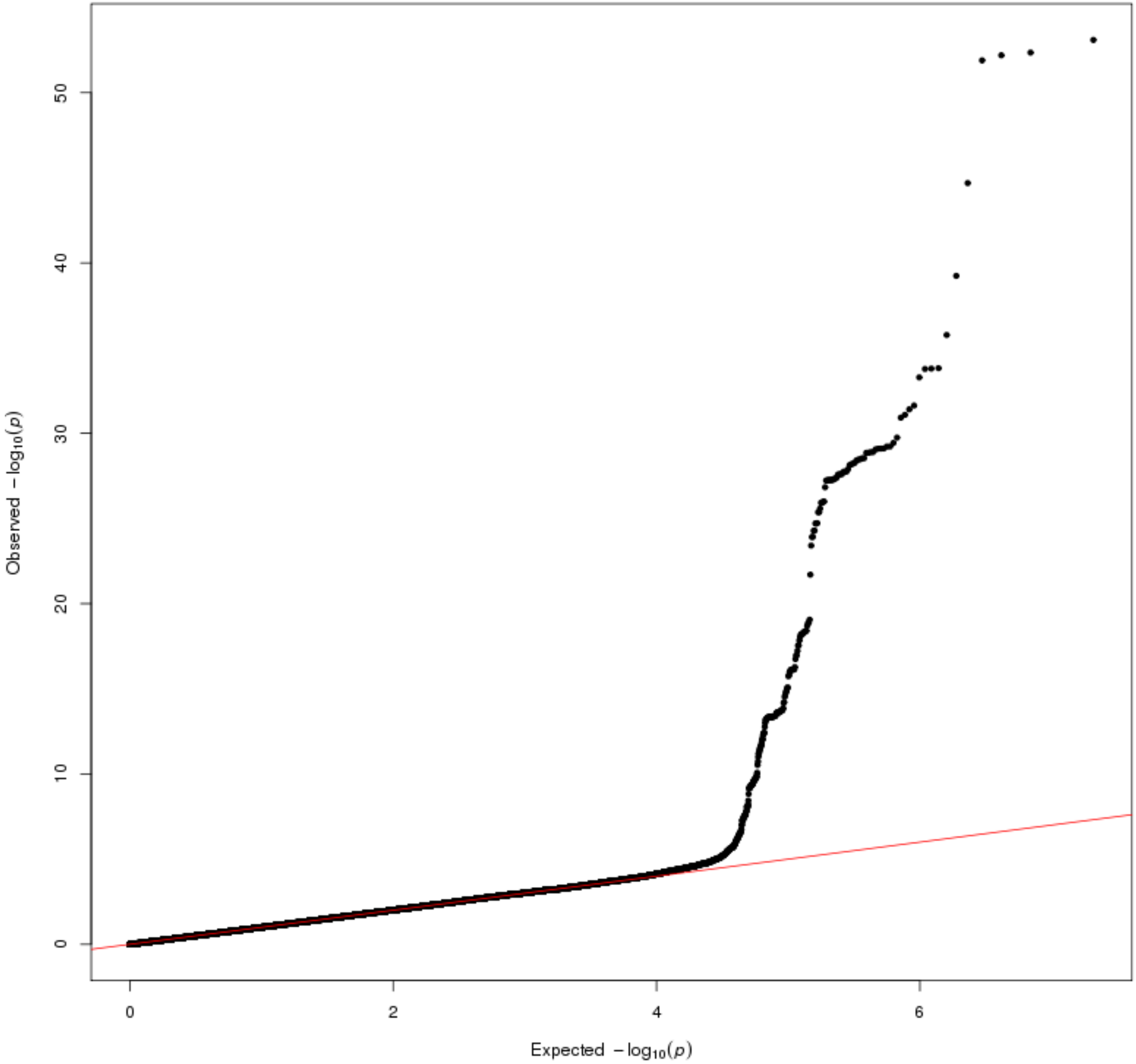

#### (viii) 3-ureidoisobutyrate

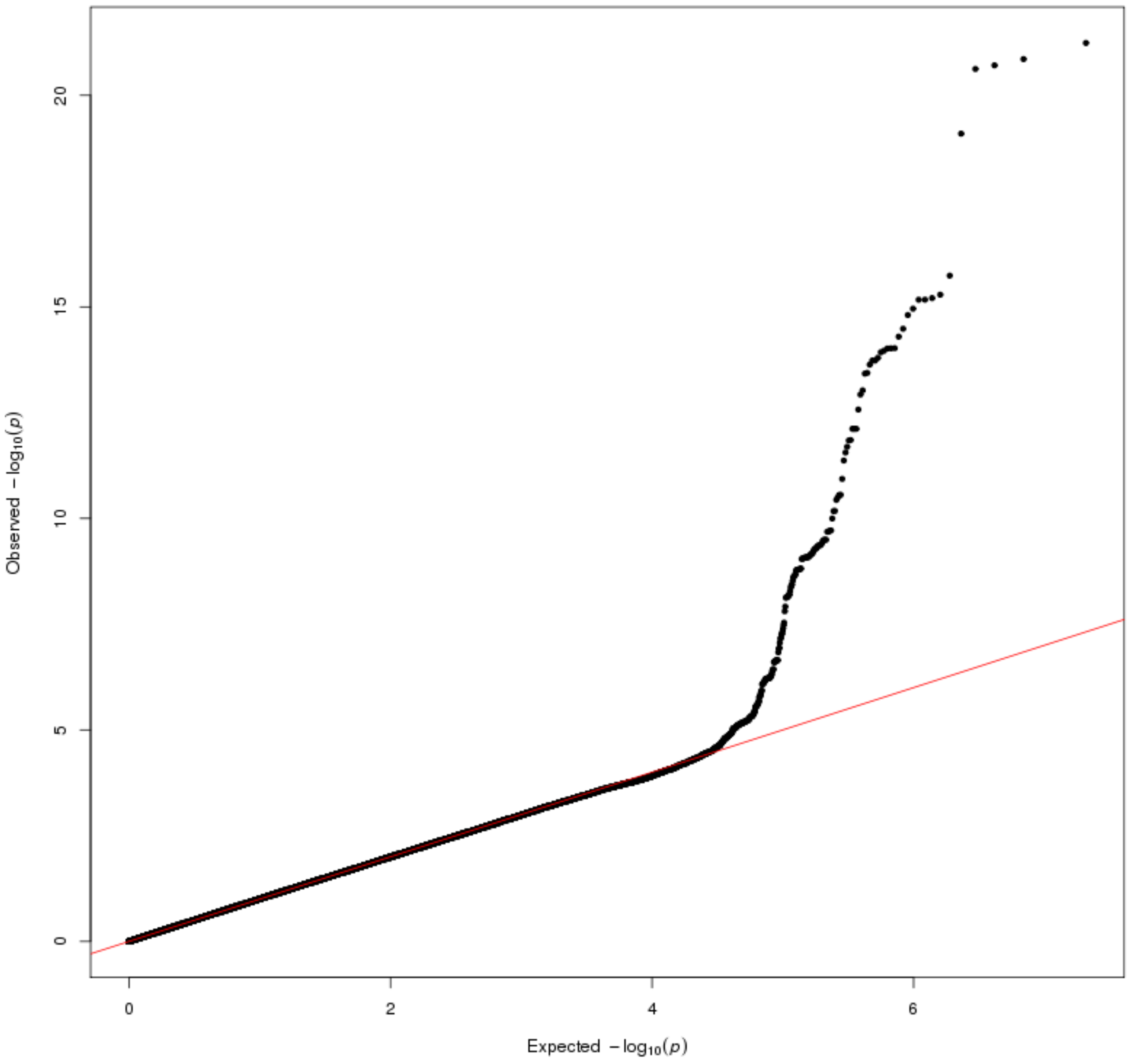

#### (ix) N-acetylglucosamine/N-acetylgalactosamine

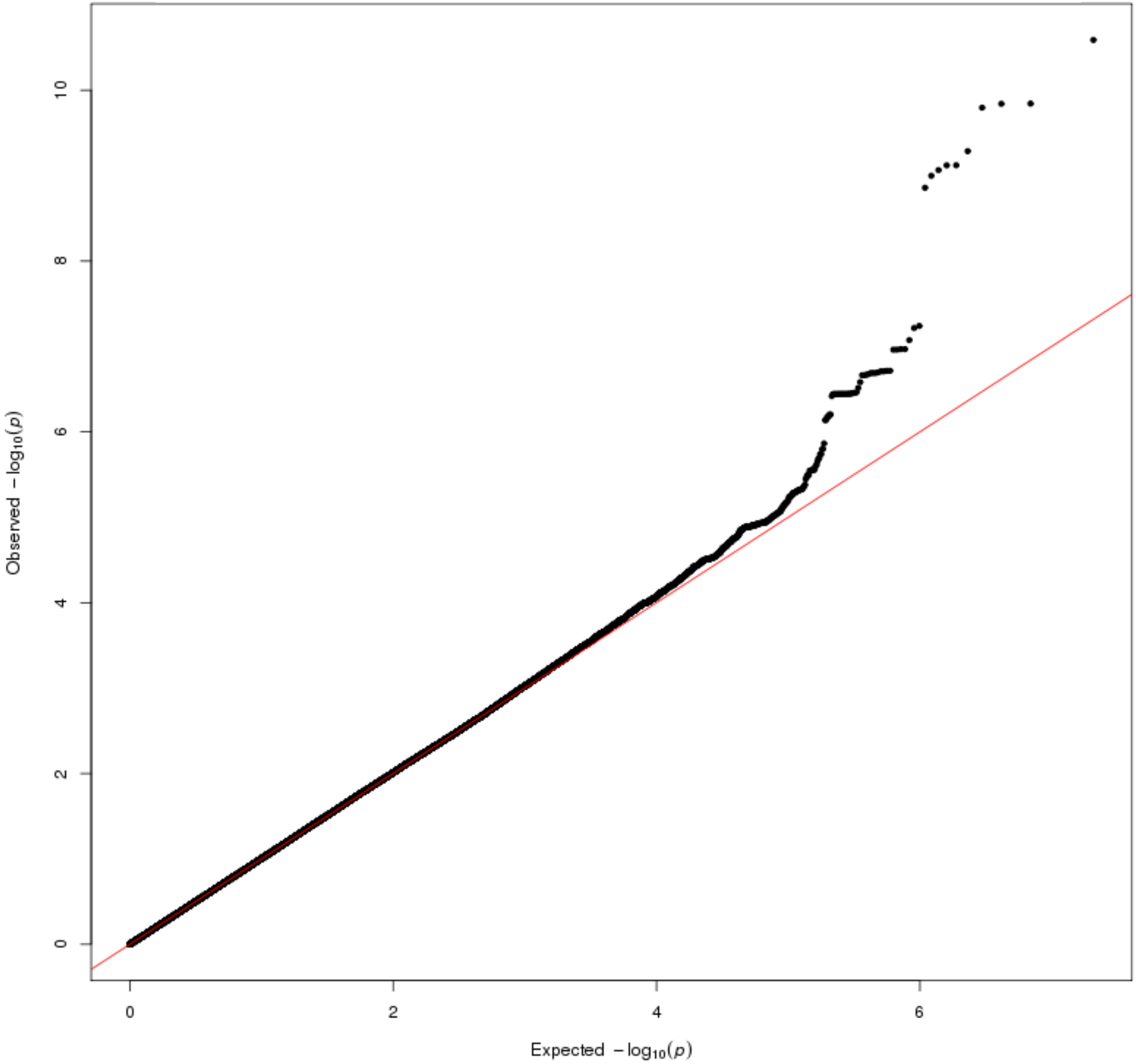

#### (x) glycosyl-N-stearoyl-sphinganine

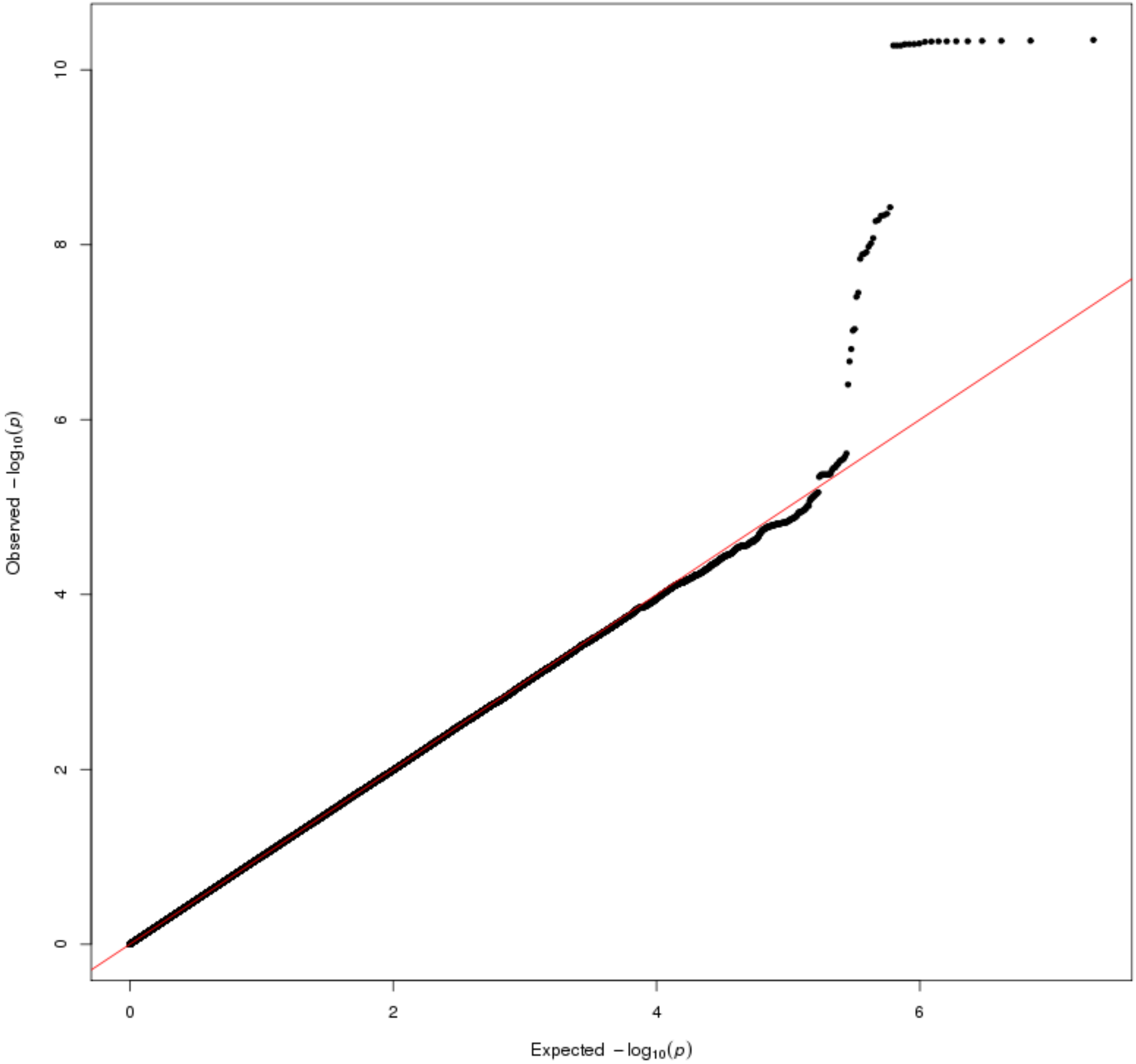

#### (xi) 1-(1-enyl-palmitoyl)-2-arachidonoyl-GPC

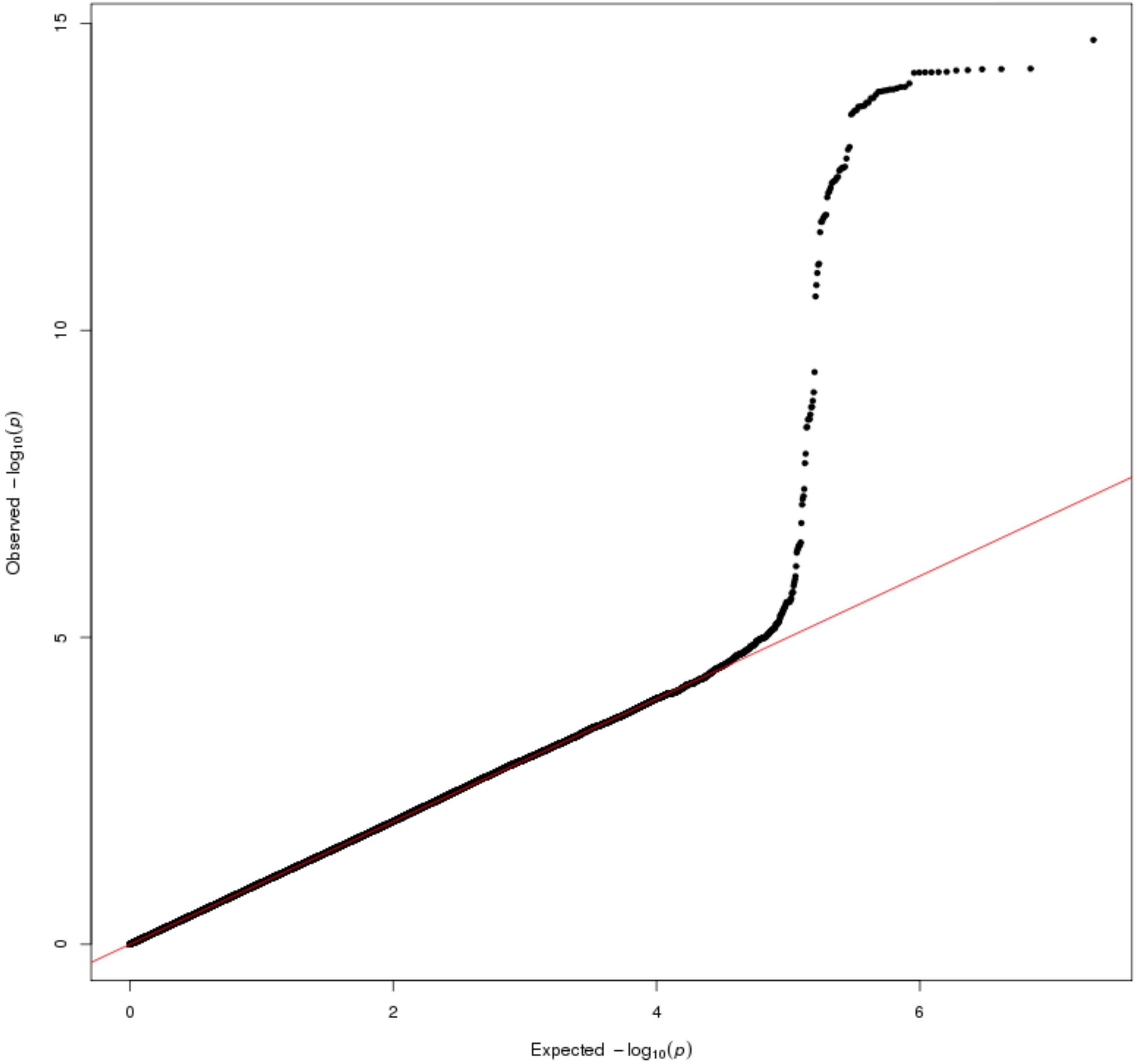

### (xii) ethylmalonate

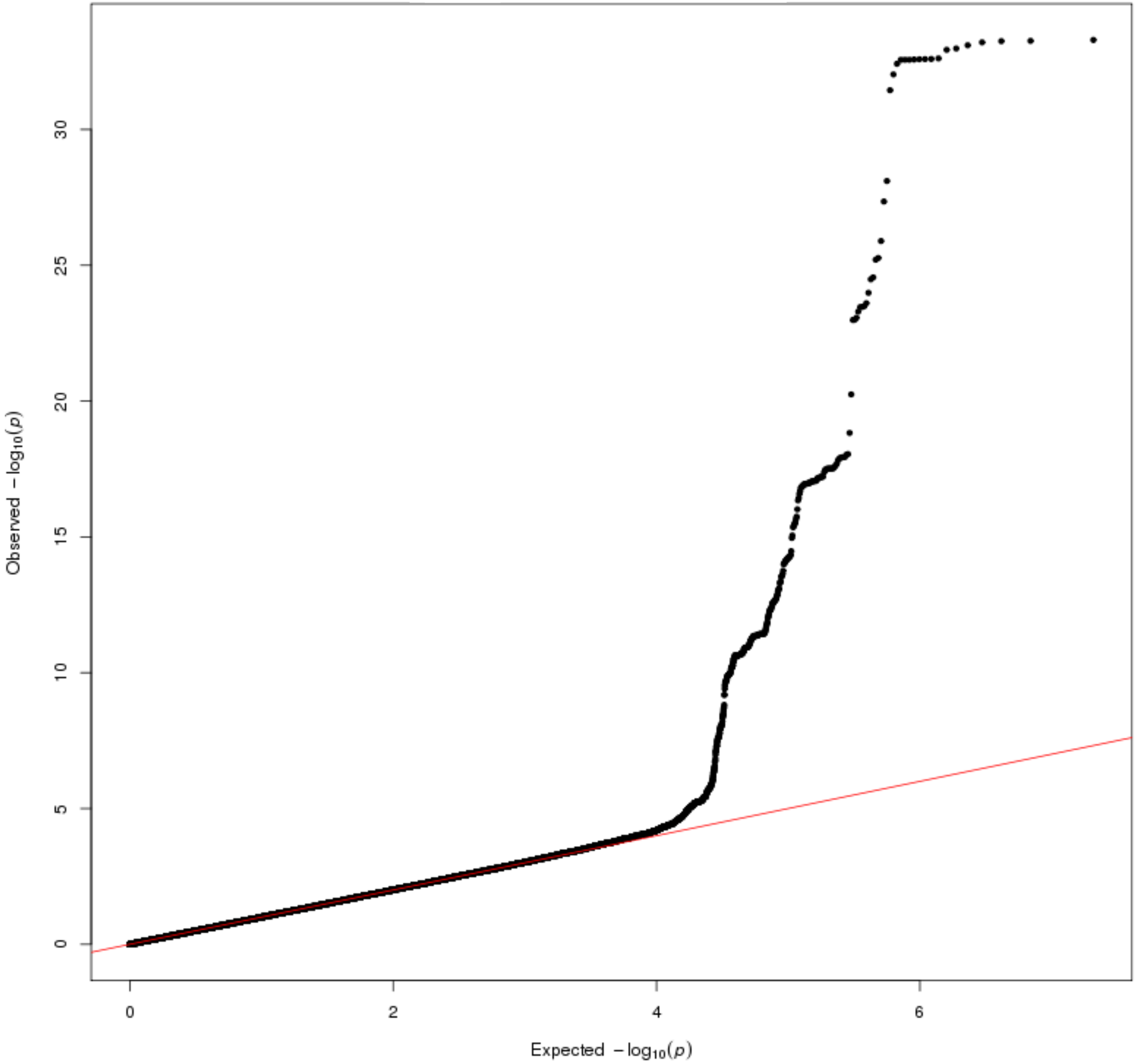

### (xiii) gamma-carboxyglutamate

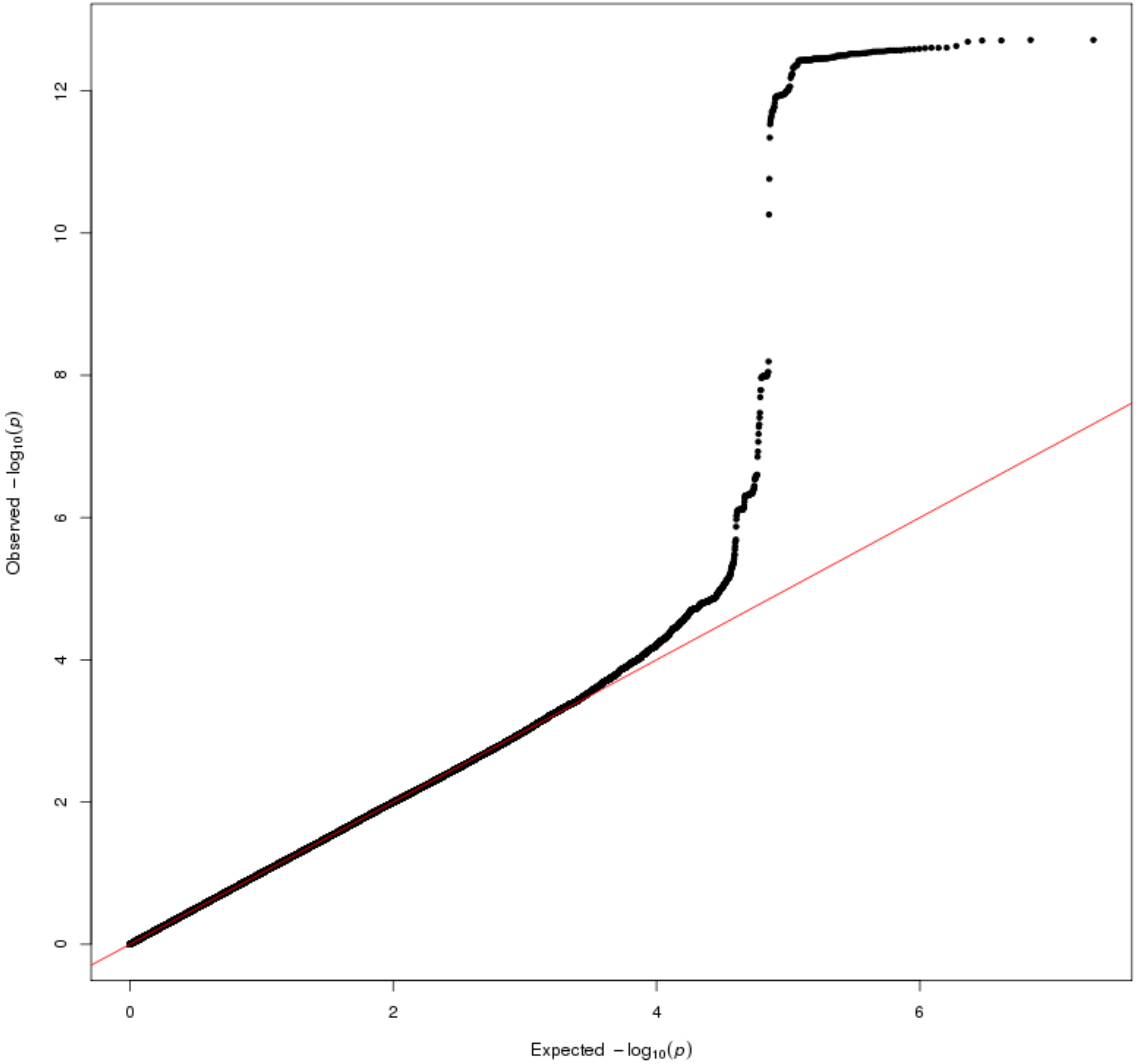

#### (xiv) ribonate

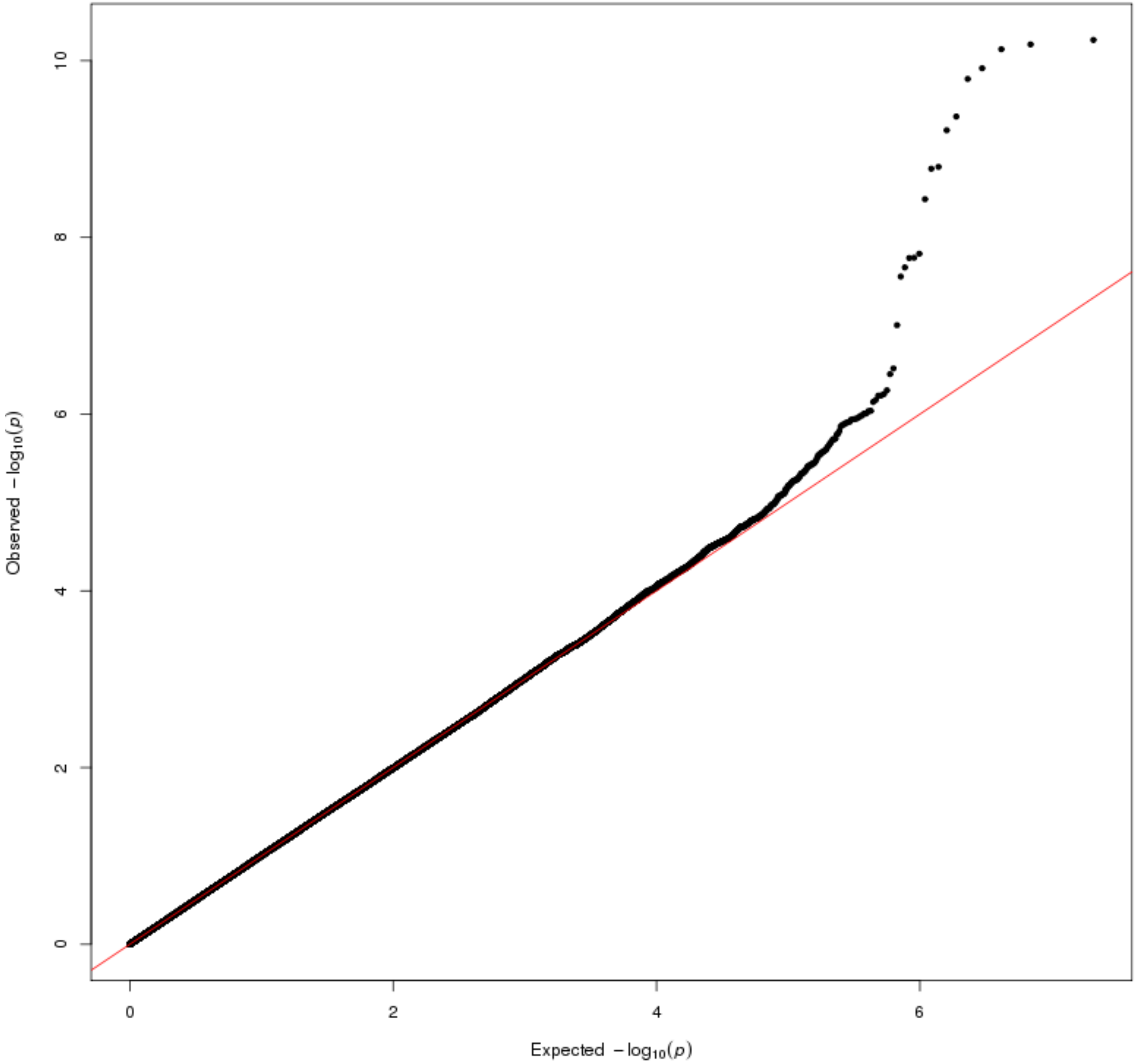

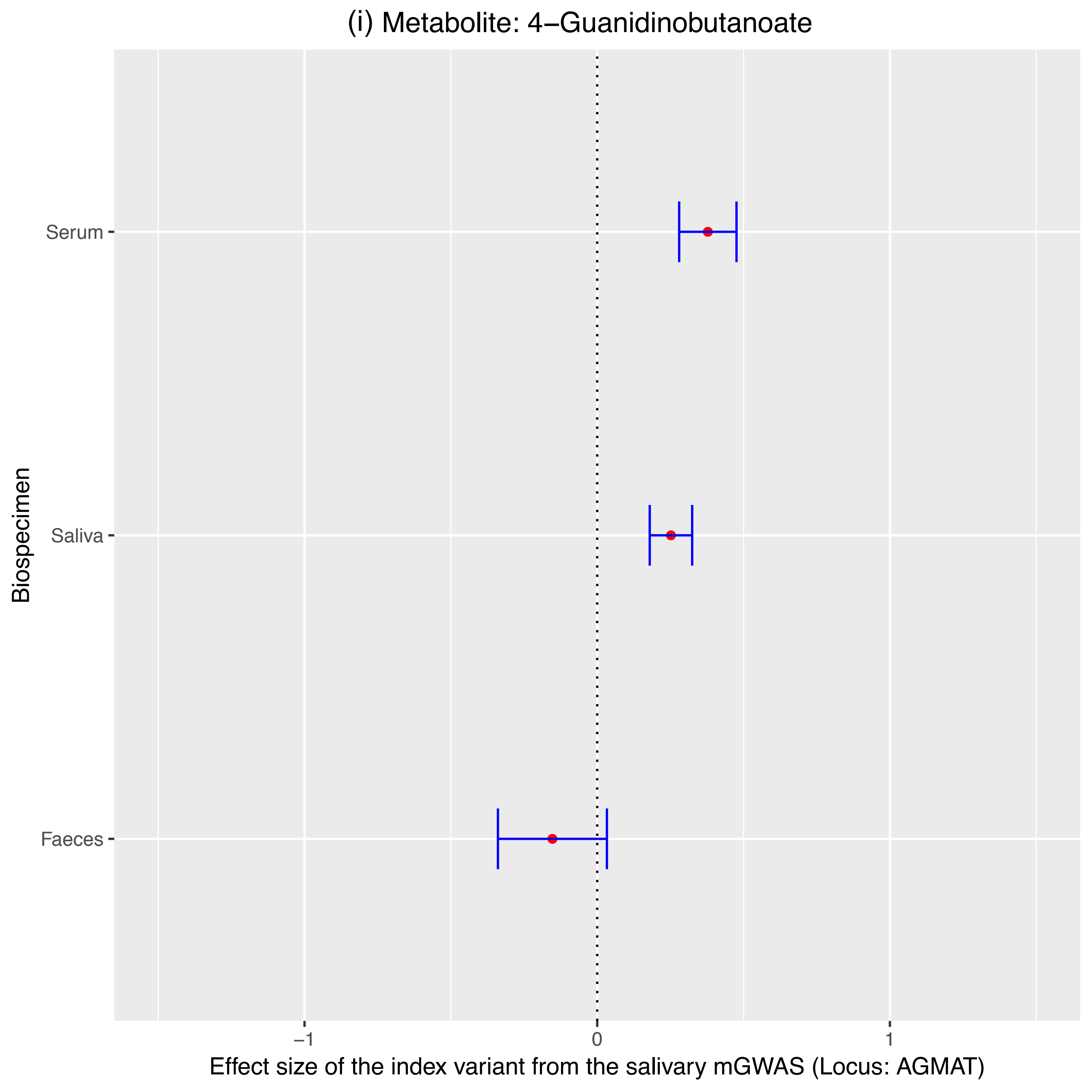
(i) Metabolite: 4−Guanidinobutanoate
Biospecimen
Serum
Saliva
Faeces
−1
0
1
Effect size of the index variant from the salivary mGWAS (Locus: AGMAT)

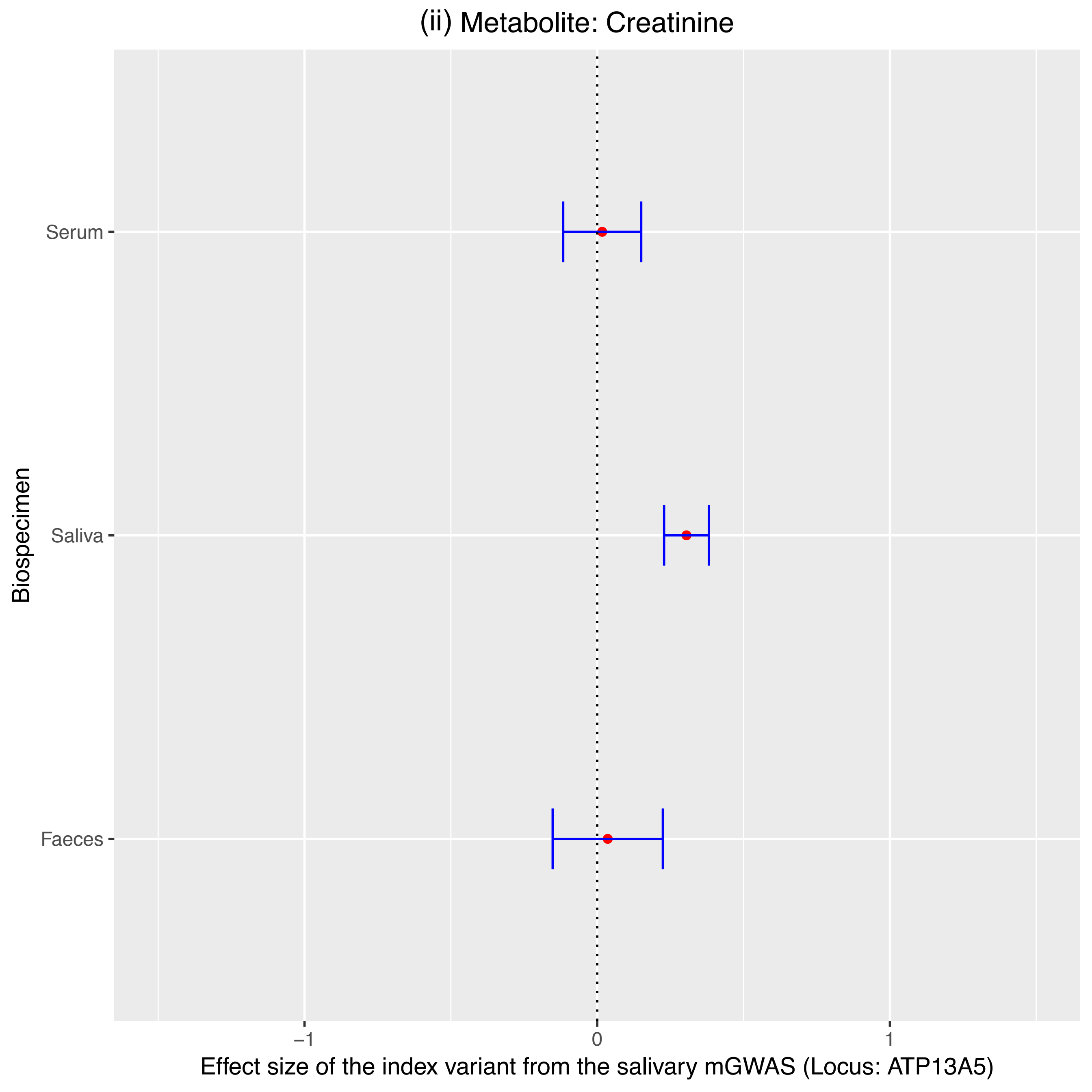
(ii) Metabolite: Creatinine
Serum
Saliva
Faeces
Biospecimen
−1
0
1
Effect size of the index variant from the salivary mGWAS (Locus: ATP13A5)

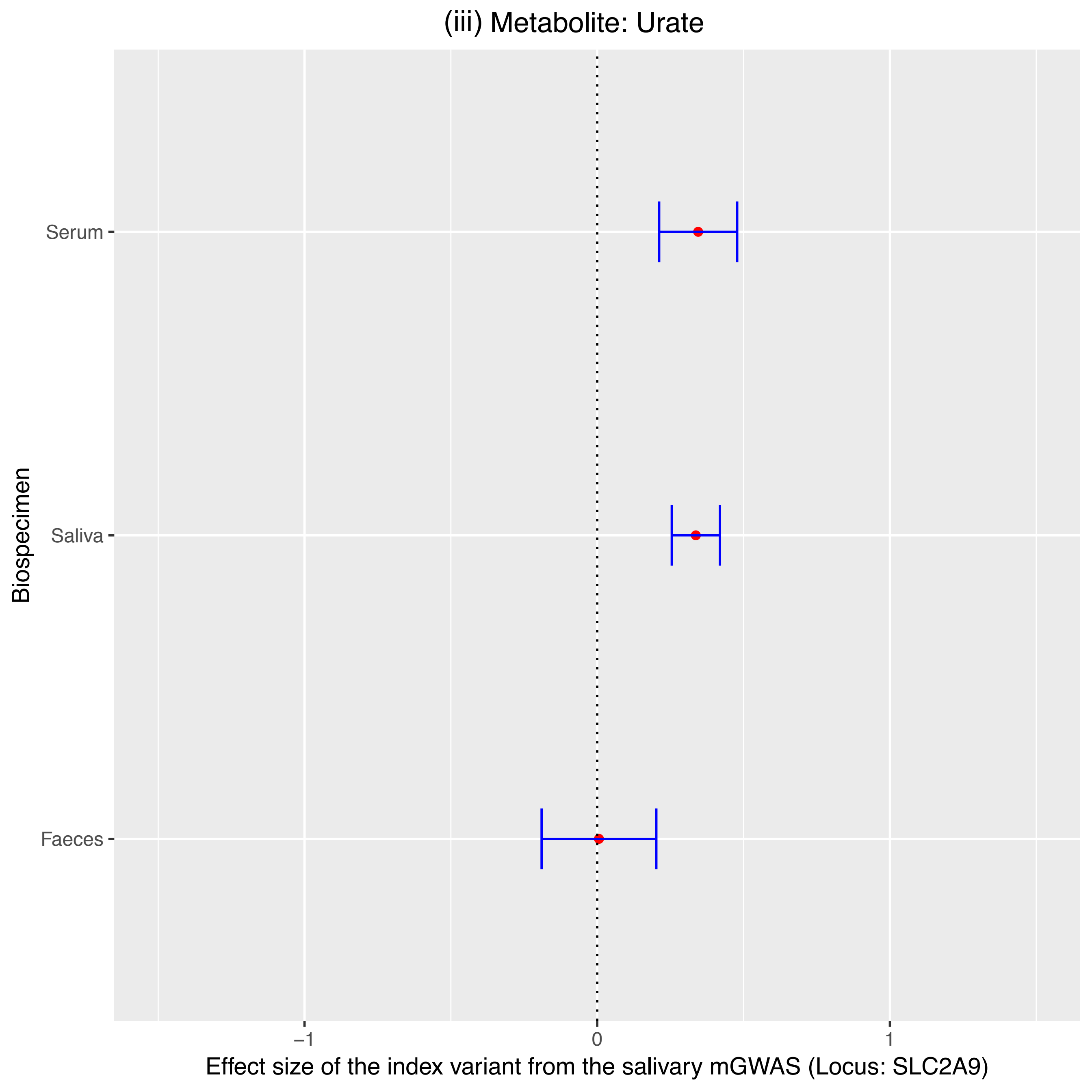
(iii) Metabolite: Urate
Serum
Saliva
Faeces
Biospecimen
−1
0
1
Effect size of the index variant from the salivary mGWAS (Locus: SLC2A9)

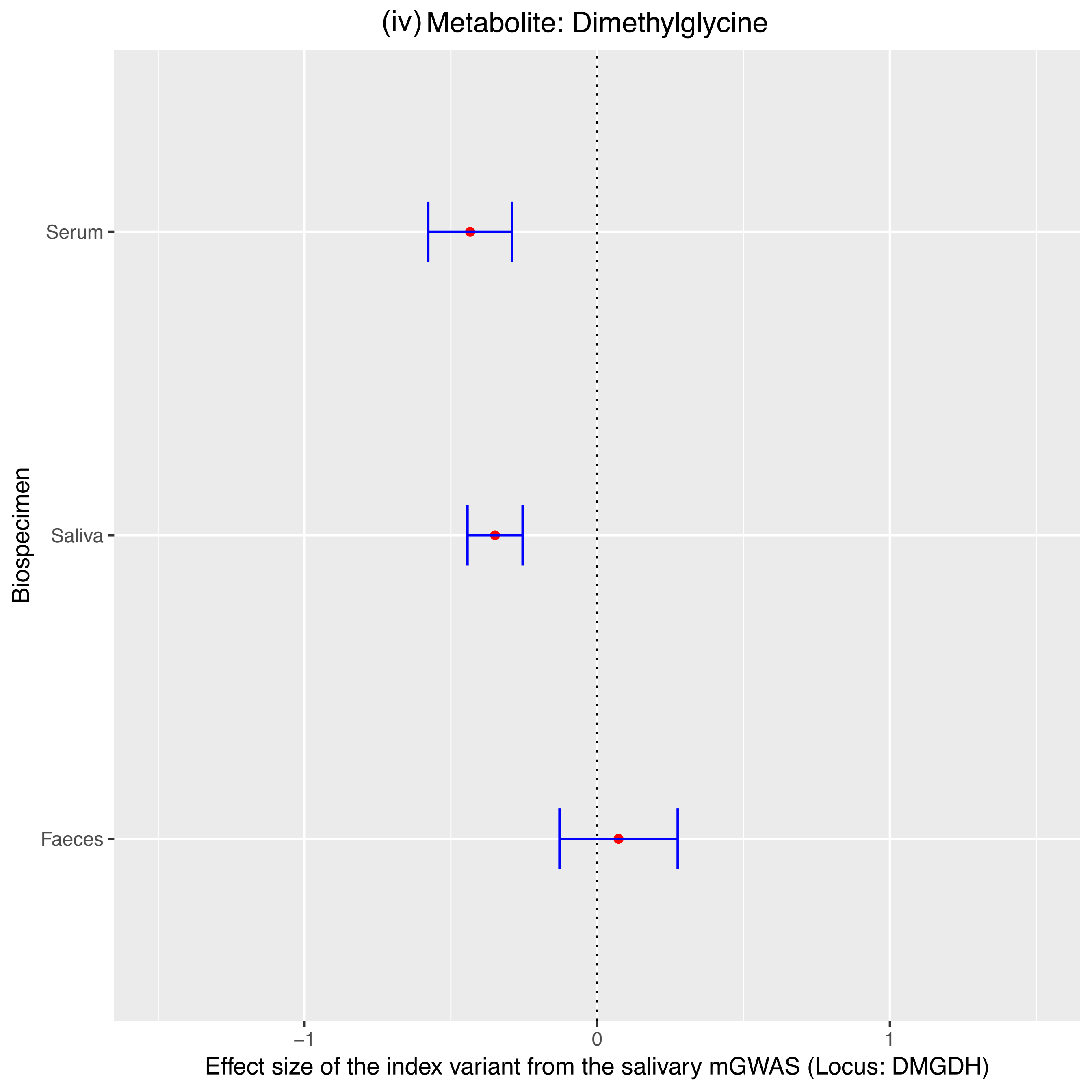

(iv) Metabolite: Dimethylglycine
Serum
Saliva
Faeces
Biospecimen
−1
0
1
Effect size of the index variant from the salivary mGWAS (Locus: DMGDH)

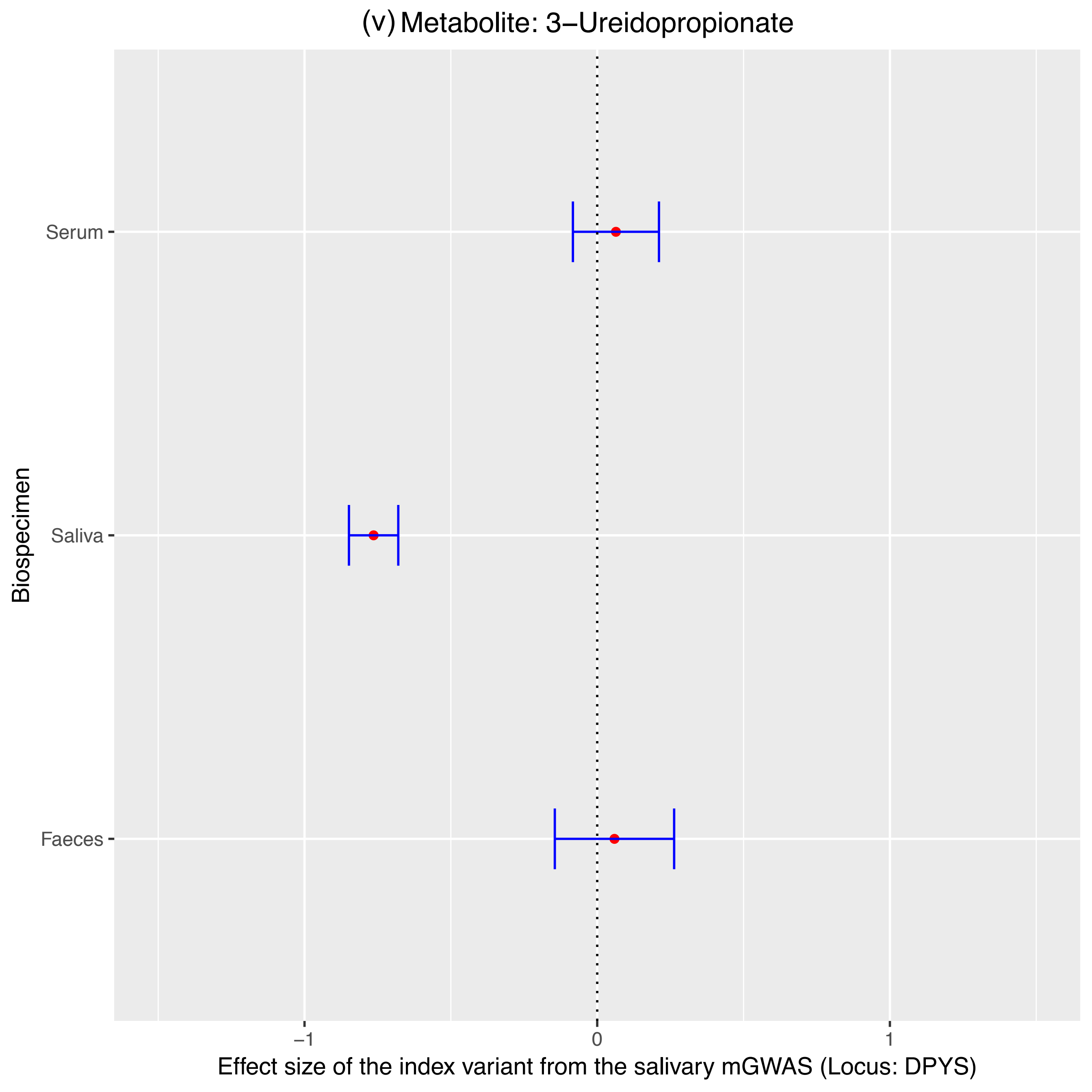
(v) Metabolite: 3−Ureidopropionate
Serum
Saliva
Faeces
Biospecimen
−1
0
1
Effect size of the index variant from the salivary mGWAS (Locus: DPYS)

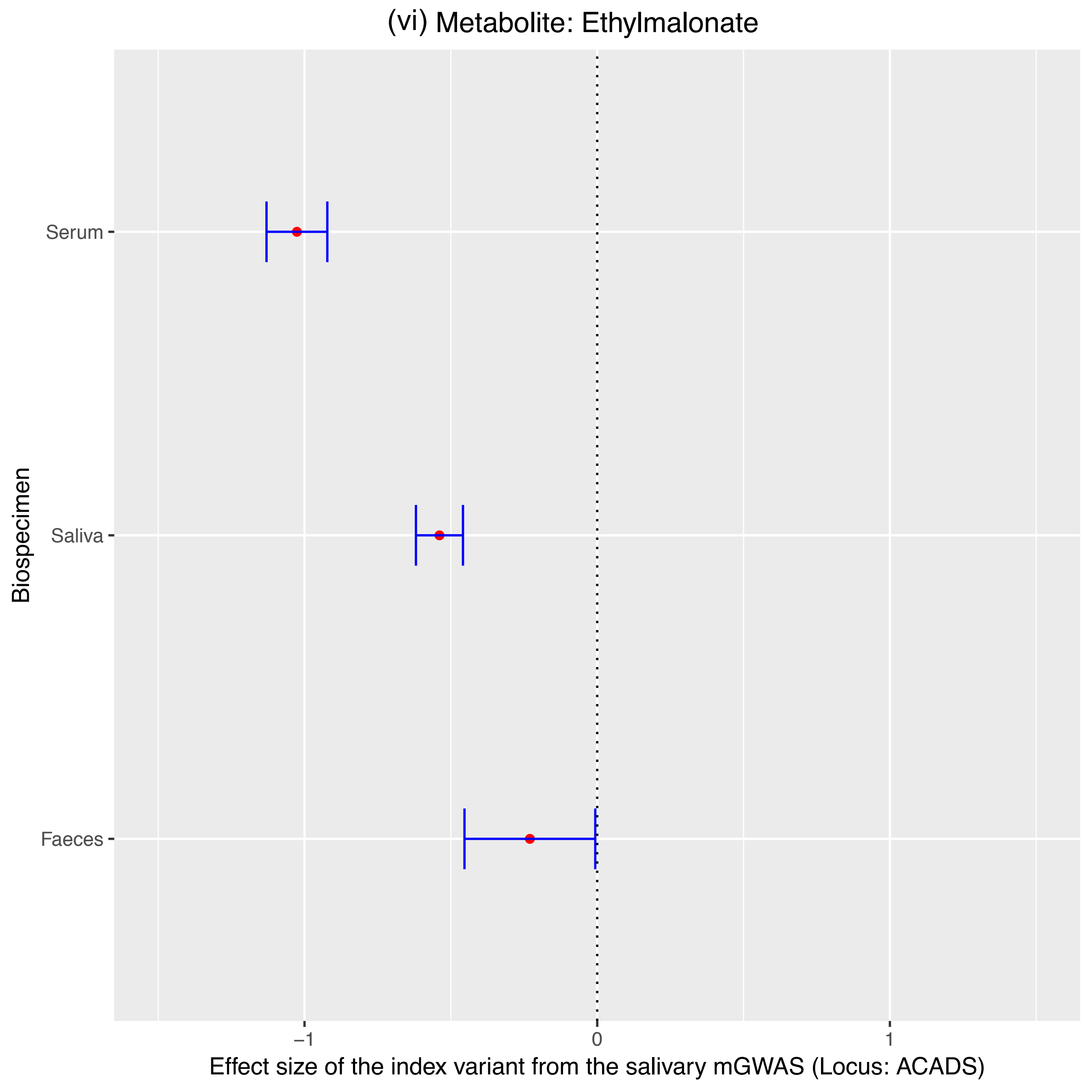

(vi) Metabolite: Ethylmalonate
Serum
Saliva
Faeces
Biospecimen
−1
0
1
Effect size of the index variant from the salivary mGWAS (Locus: ACADS)

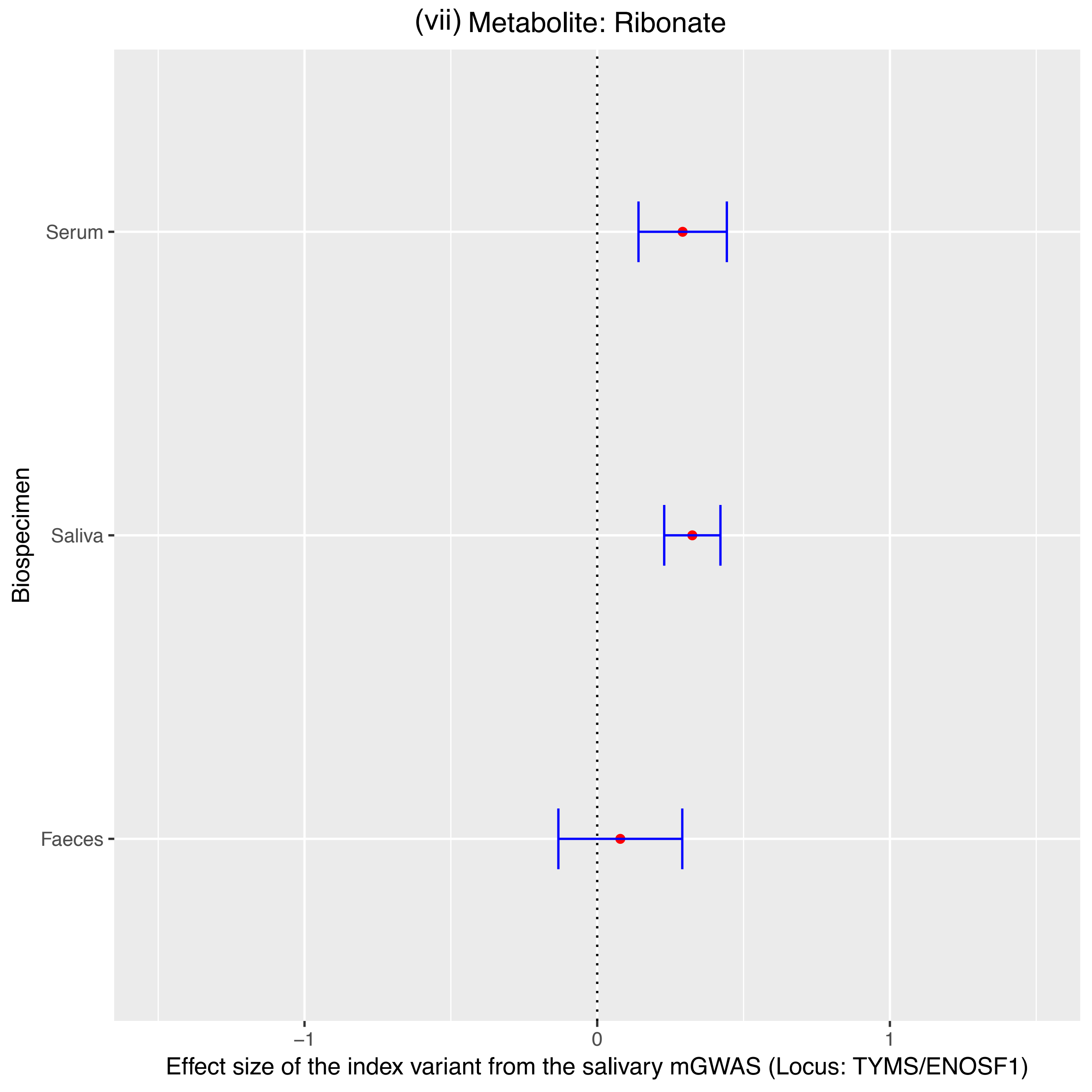

(vii) Metabolite: Ribonate
Serum
Saliva
Faeces
Biospecimen
−1
0
1
Effect size of the index variant from the salivary mGWAS (Locus: TYMS/ENOSF1)

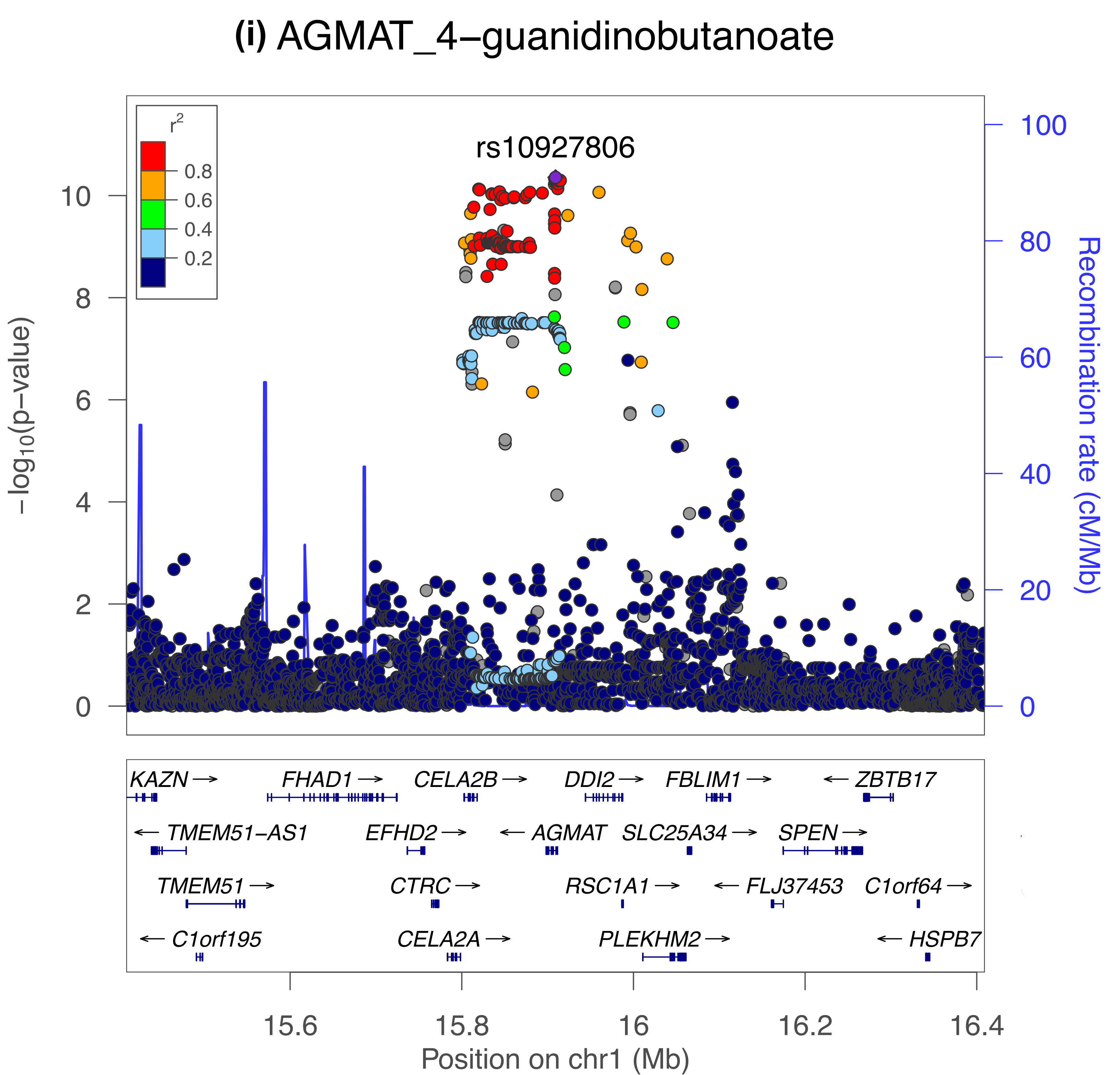

(i) AGMAT_4−guanidinobutanoate
rs10927806
r2
0.8
0.6
0.4
0.2
−log10(p−value)
Recombination rate (cM/Mb)
0
2
4
6
8
10
0
20
40
60
80
100
KAZN →
FHAD1 →
CELA2B →
DDI2 →
FBLIM1 →
← ZBTB17
← TMEM51−AS1
EFHD2 →
← AGMAT
SLC25A34 →
SPEN →
TMEM51 →
CTRC →
RSC1A1 →
← FLJ37453
C1orf64 →
← C1orf195
CELA2A →
PLEKHM2 →
← HSPB7
15.6
15.8
16
16.2
16.4
Position on chr1 (Mb)

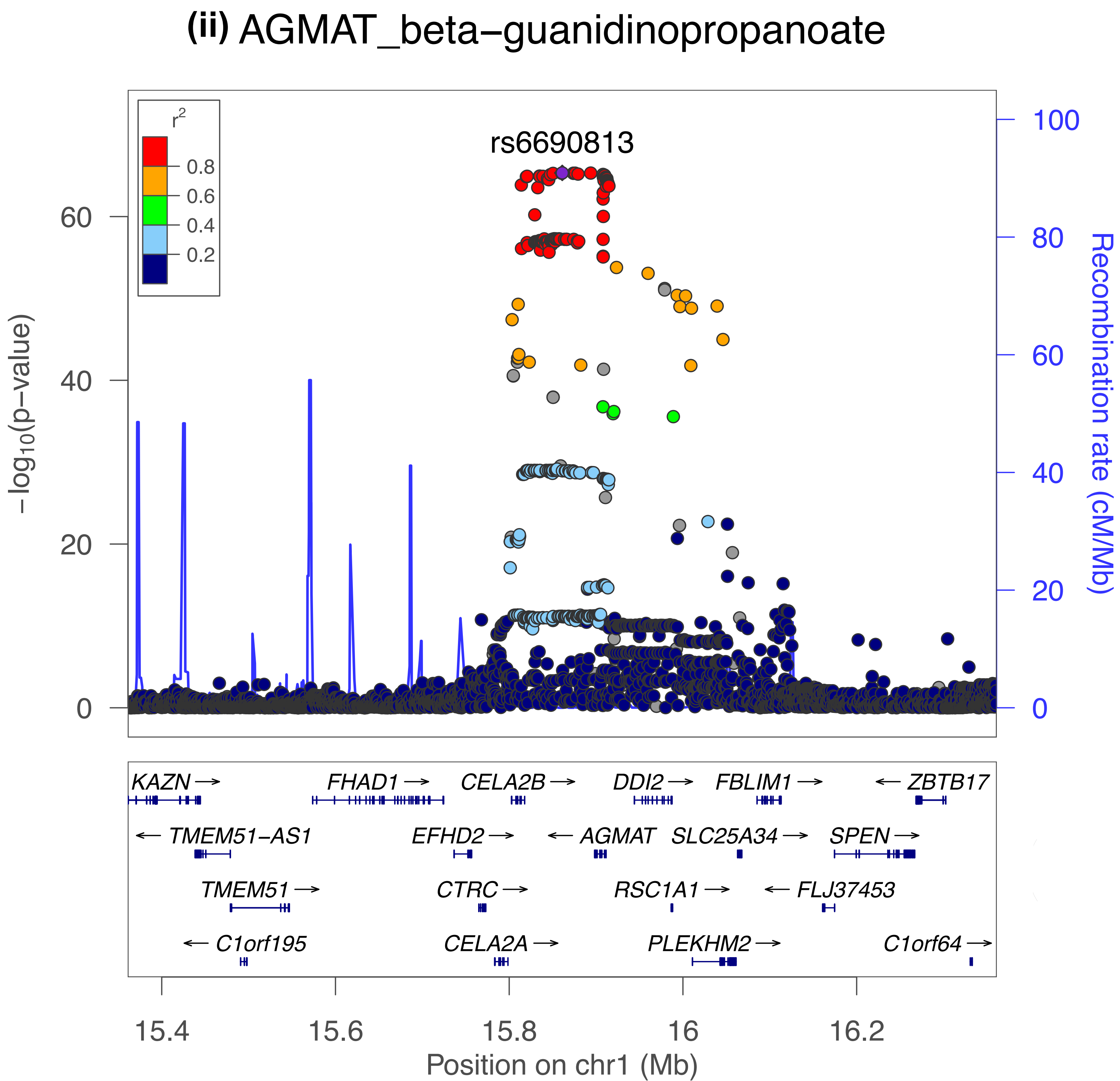

(ii) AGMAT_beta−guanidinopropanoate
r2
0.8
0.6
0.4
0.2
rs6690813
−log10(p−value)
0
20
40
60
Recombination rate (cM/Mb)
100
80
60
40
20
0
KAZN
TMEM51−AS1
TMEM51
C1orf195
FHAD1
EFHD2
CTRC
CELA2A
CELA2B
AGMAT
RSC1A1
PLEKHM2
DDI2
SLC25A34
FBLIM1
FLJ37453
SPEN
ZBTB17
C1orf64
15.4
15.6
15.8
16
16.2
Position on chr1 (Mb)

(iii) ATP13A5_creatinine

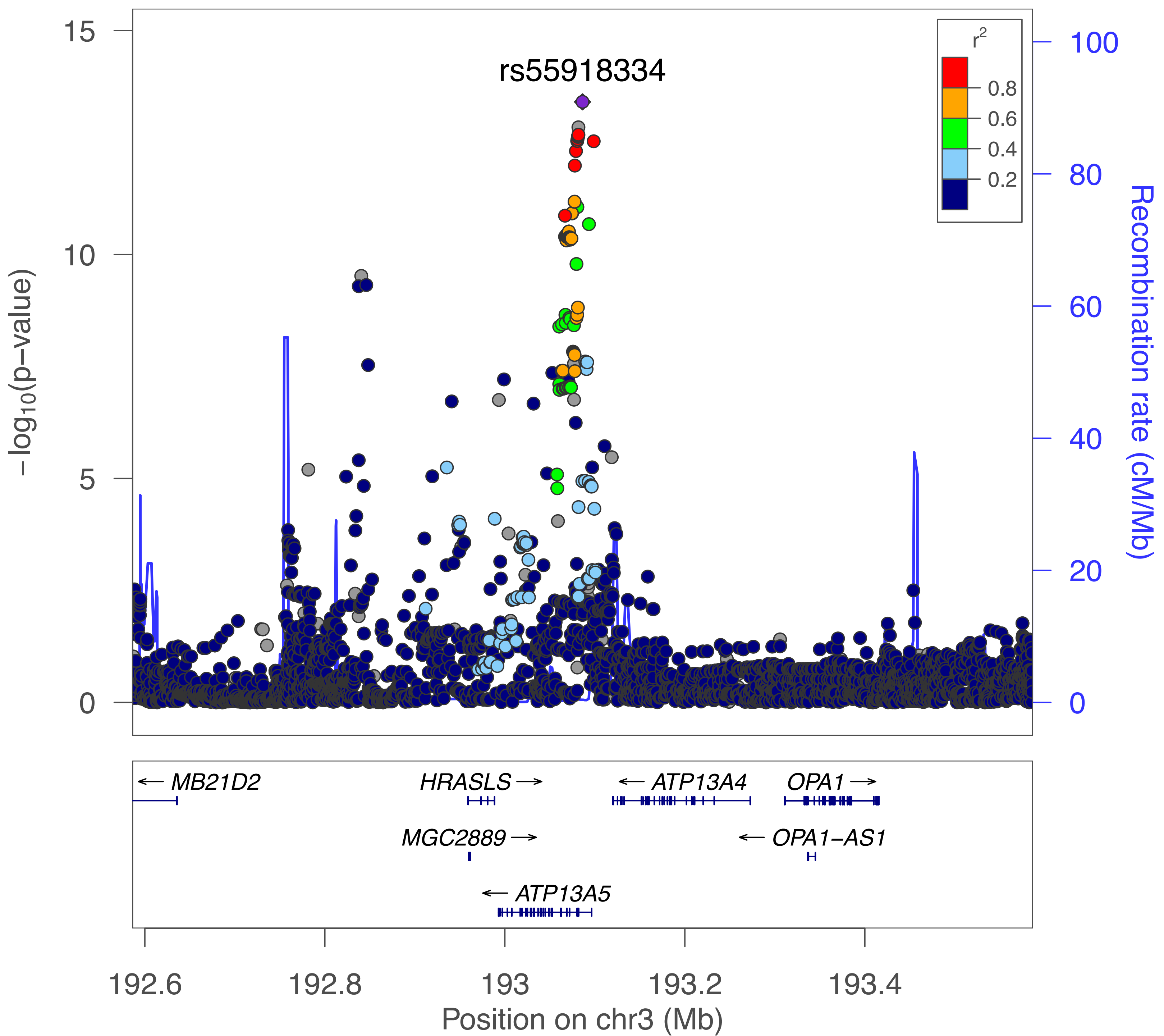

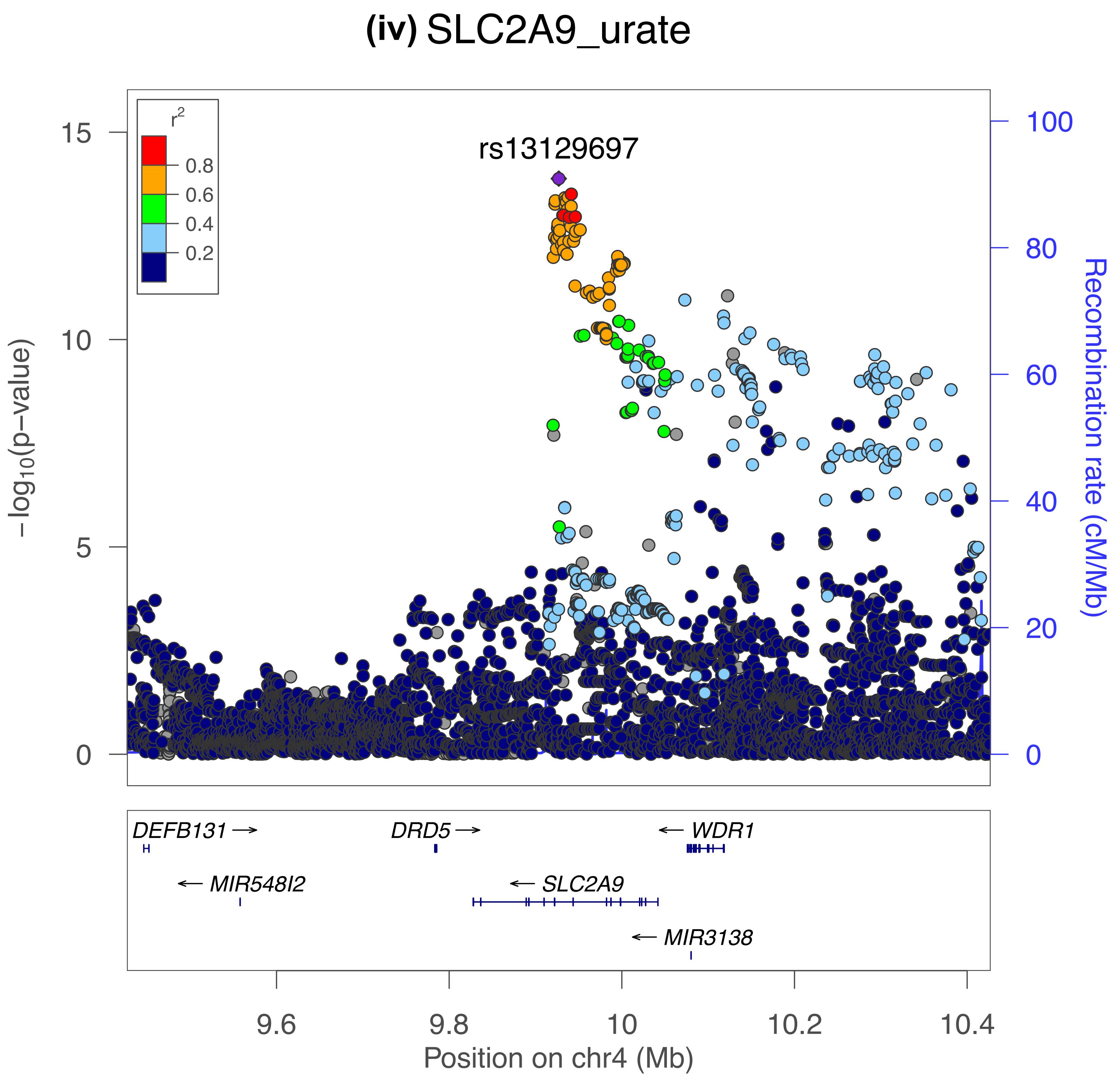

(iv) SLC2A9_urate
rs13129697
r2
0.8
0.6
0.4
0.2
15
10
5
0
-log10(p-value)
100
80
60
40
20
0
Recombination rate (cM/Mb)
DEFB131
DRD5
WDR1
MIR548I2
SLC2A9
MIR3138
9.6
9.8
10
10.2
10.4
Position on chr4 (Mb)

(v) SLC2A9_allantoin

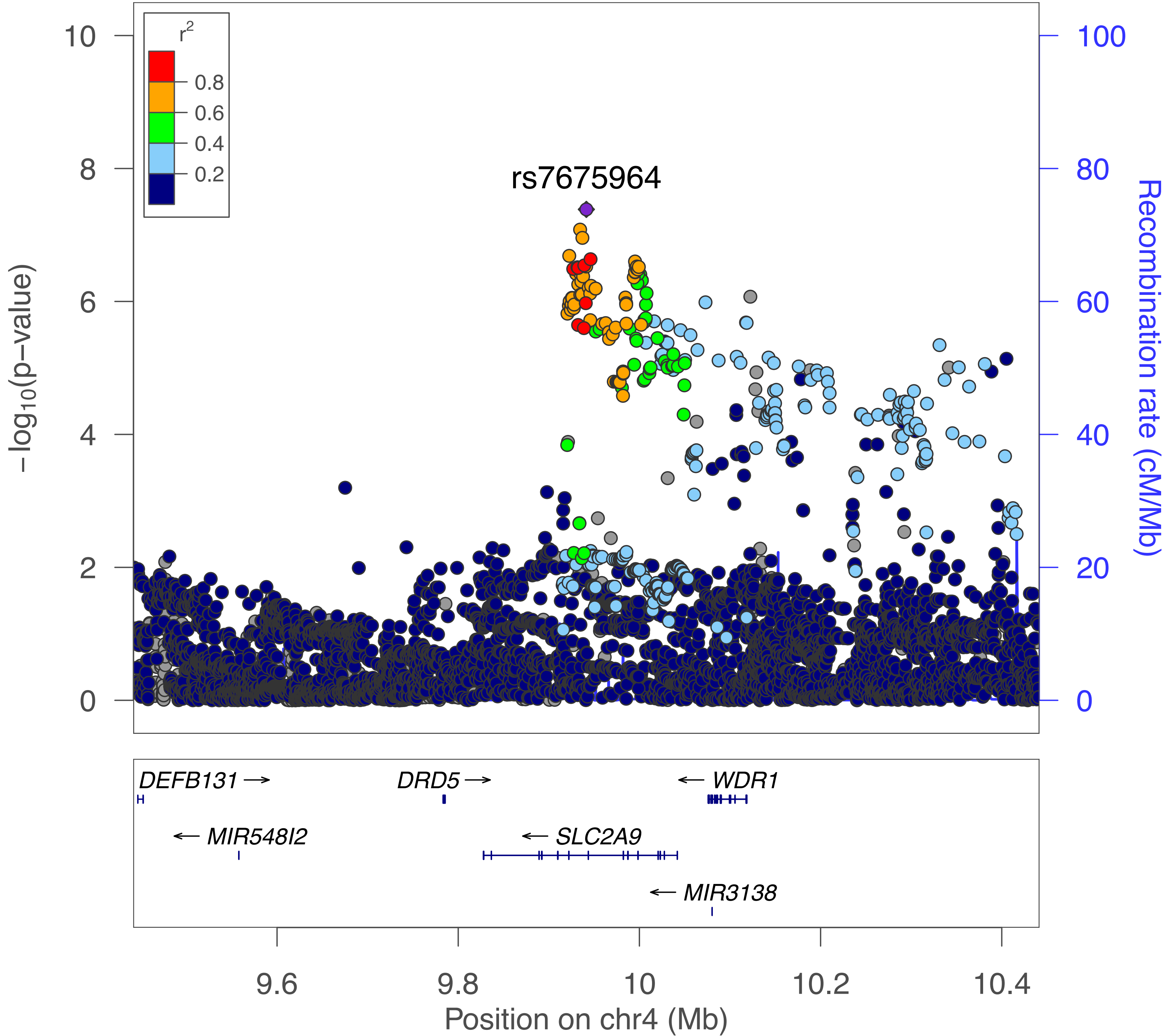

#### (vi) DMGDH_dimethylglycine

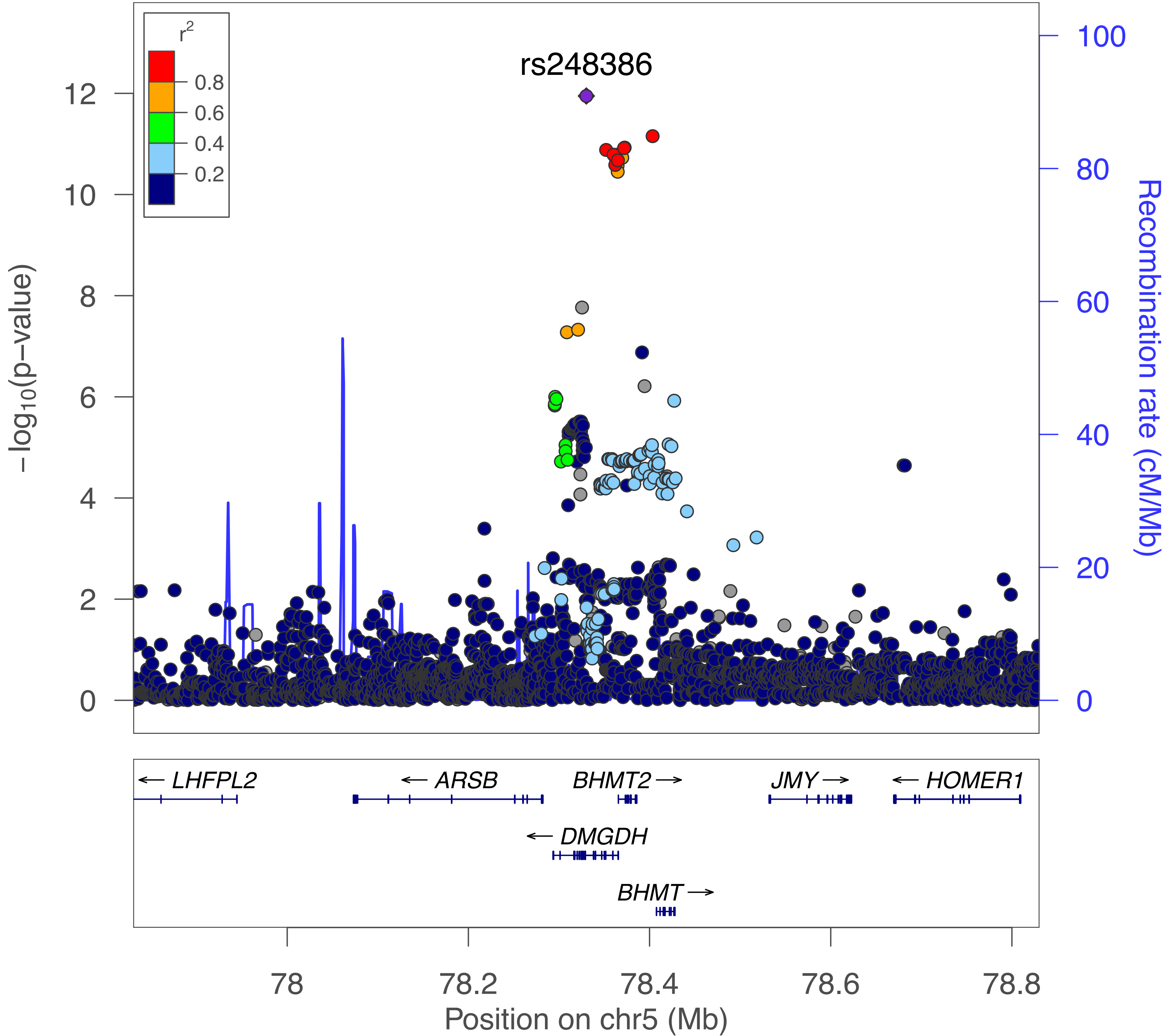

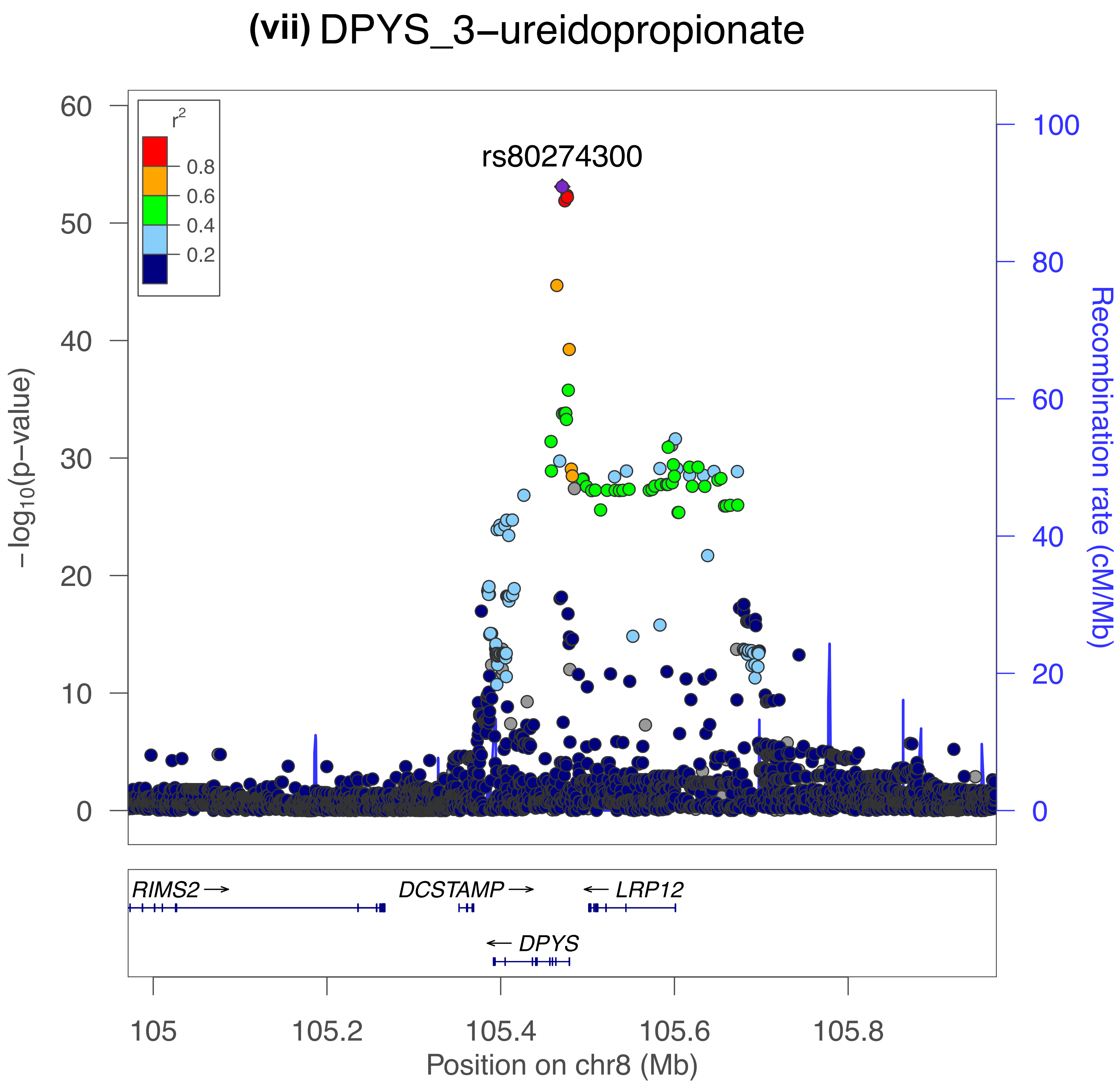
(vii) DPYS_3−ureidopropionate
rs80274300
r2
0.8
0.6
0.4
0.2
60
50
40
30
20
10
0
−log10(p−value)
100
80
60
40
20
0
Recombination rate (cM/Mb)
RIMS2 →
DCSTAMP →
← LRP12
← DPYS
105
105.2
105.4
105.6
105.8
Position on chr8 (Mb)

**(viii)** DPYS_3−ureidoisobutyrate

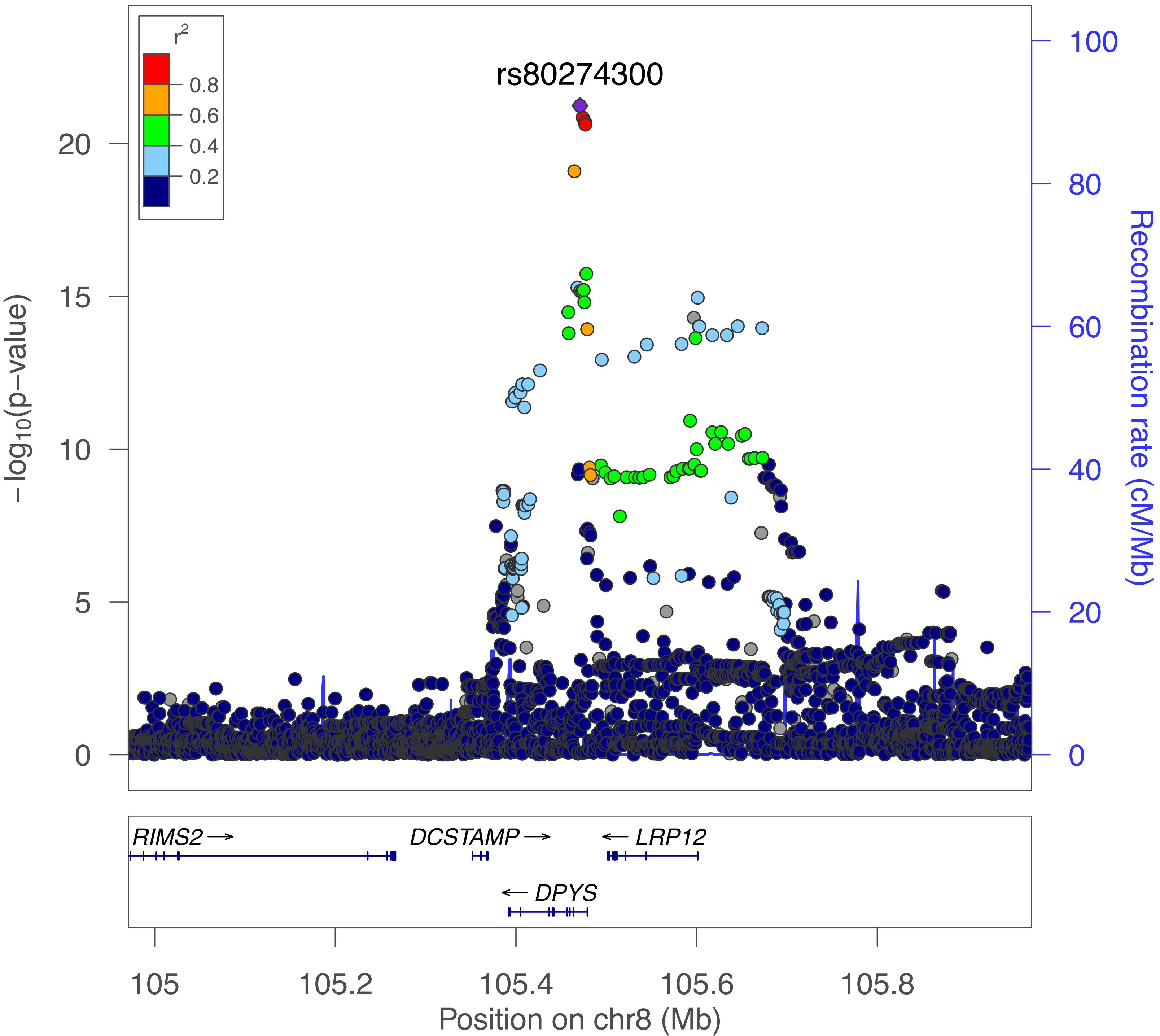

**(ix)** ABO_N-acetylglucosamine/N-acetylgalactosamine

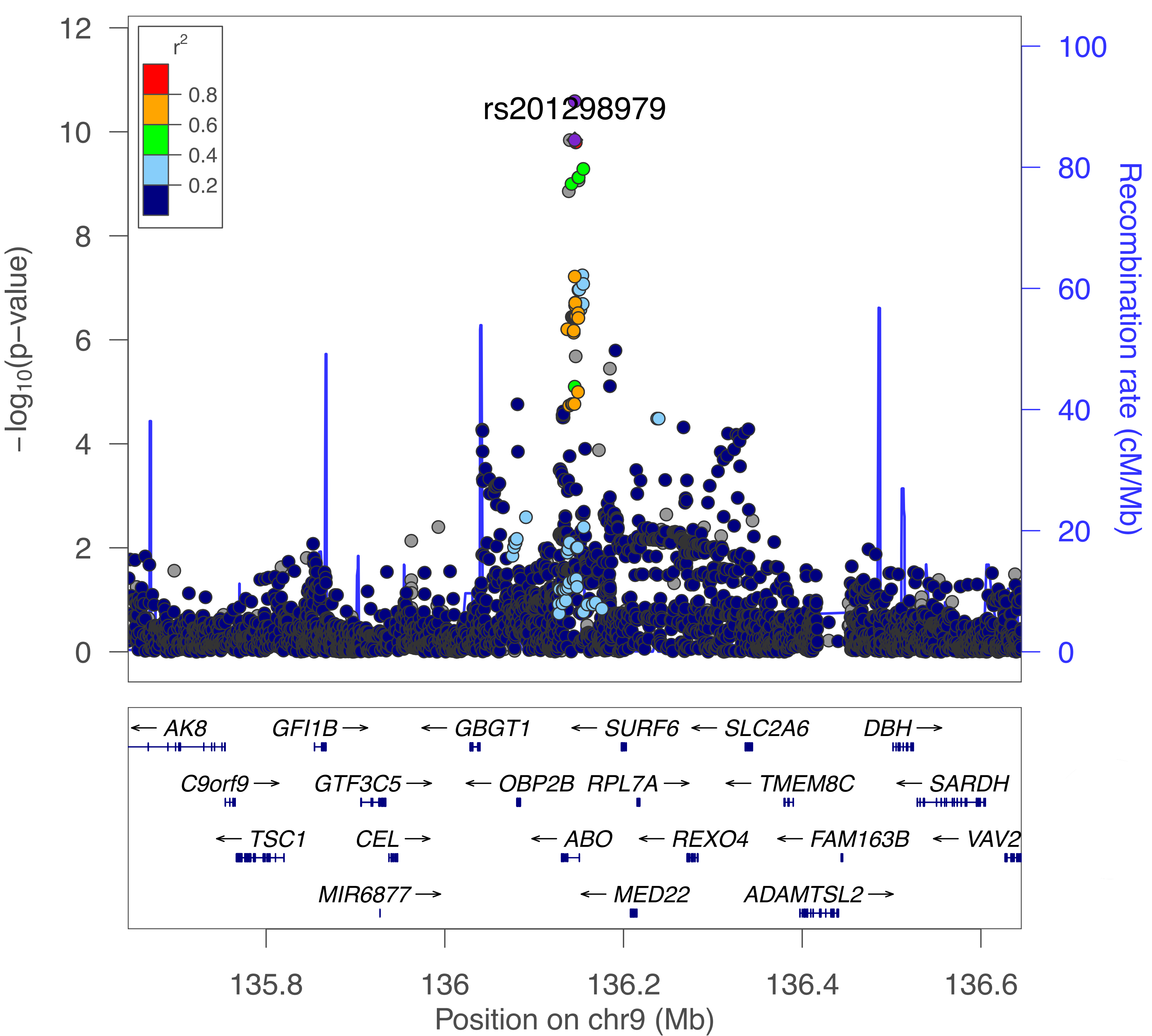

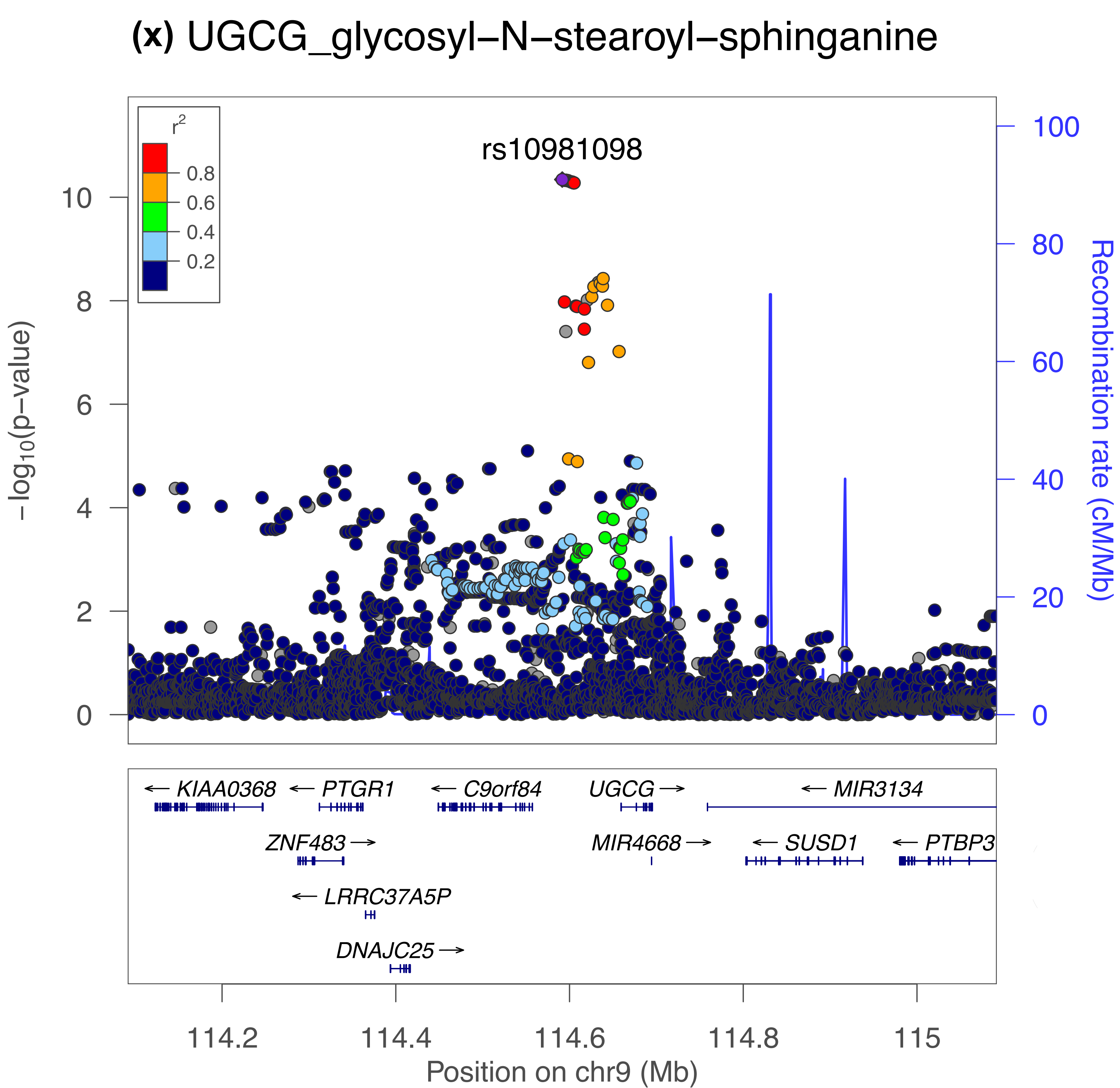

(x) UGCG_glycosyl–N–stearoyl–sphinganine
rs10981098
r2
0.8
0.6
0.4
0.2
−log10(p−value)
0
2
4
6
8
10
Recombination rate (cM/Mb)
0
20
40
60
80
100
KIAA0368
PTGR1
C9orf84
UGCG
MIR3134
ZNF483
MIR4668
SUSD1
PTBP3
LRRC37A5P
DNAJC25
114.2
114.4
114.6
114.8
115
Position on chr9 (Mb)

**(xi)** FADS2_1-(1-enyl-palmitoyl)-2-arachidonoyl-GPC

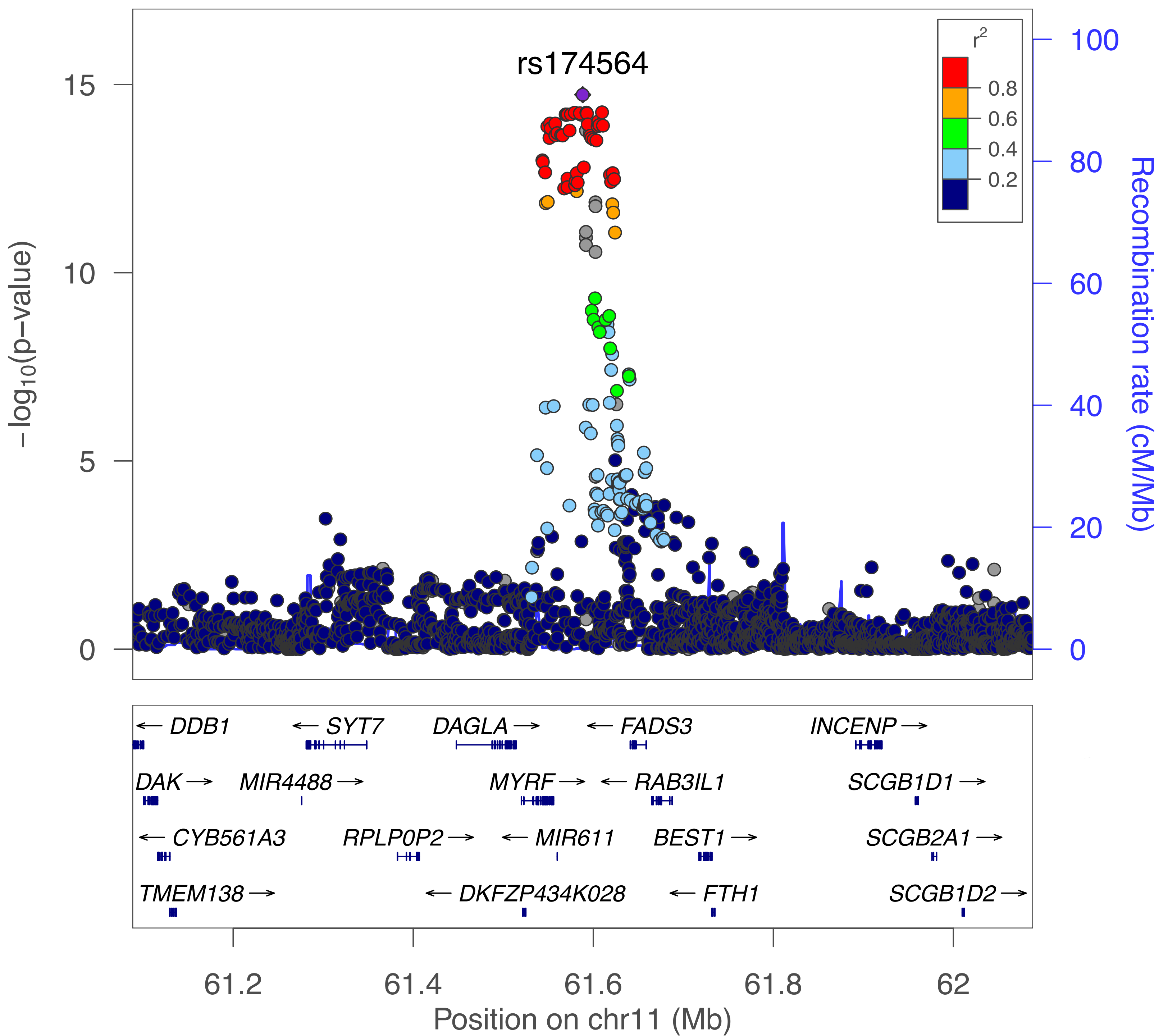

#### (xii) ACADS_ethylmalonate

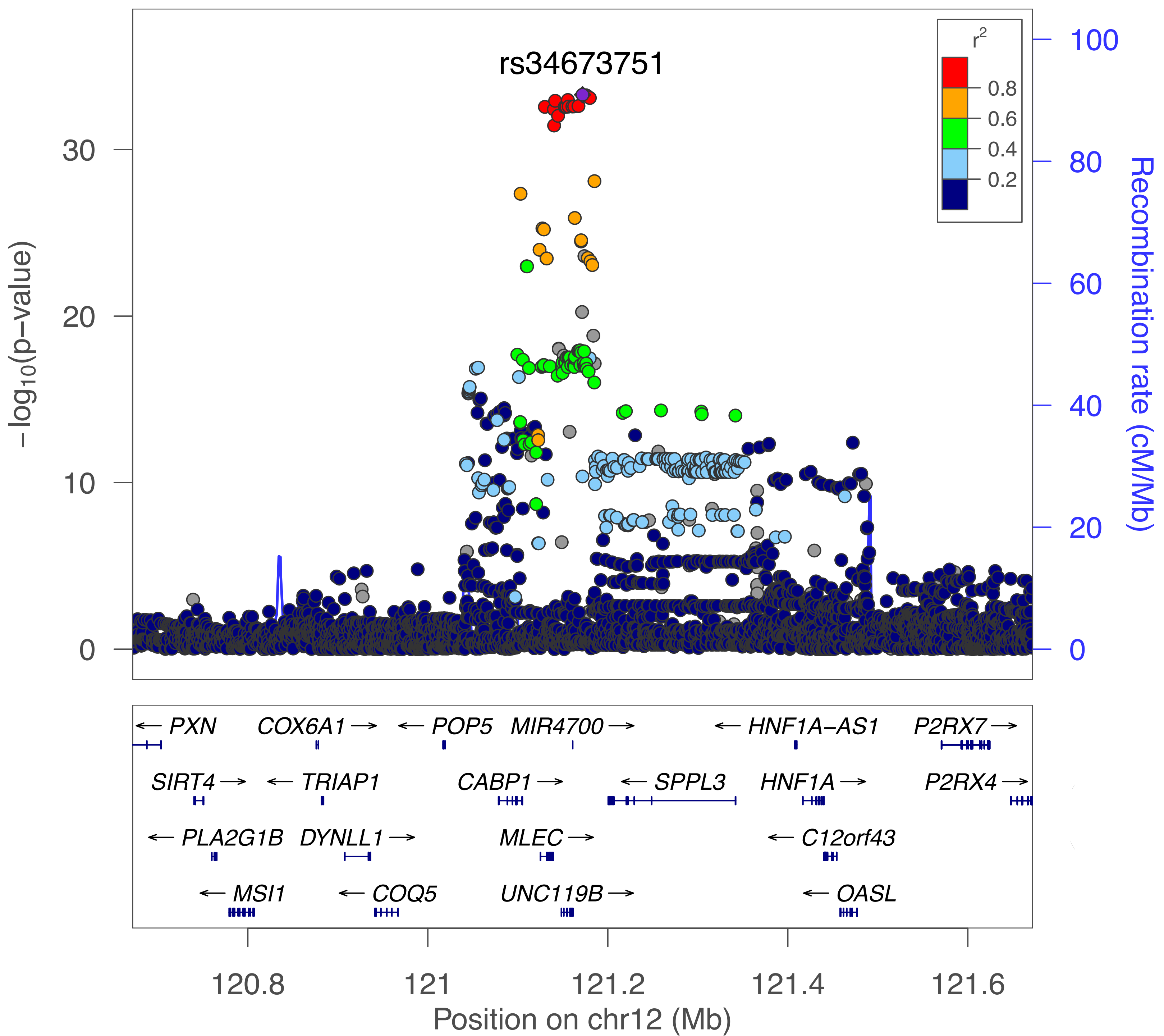

**(xiii)** TYMS/ENOSF1_ribonate

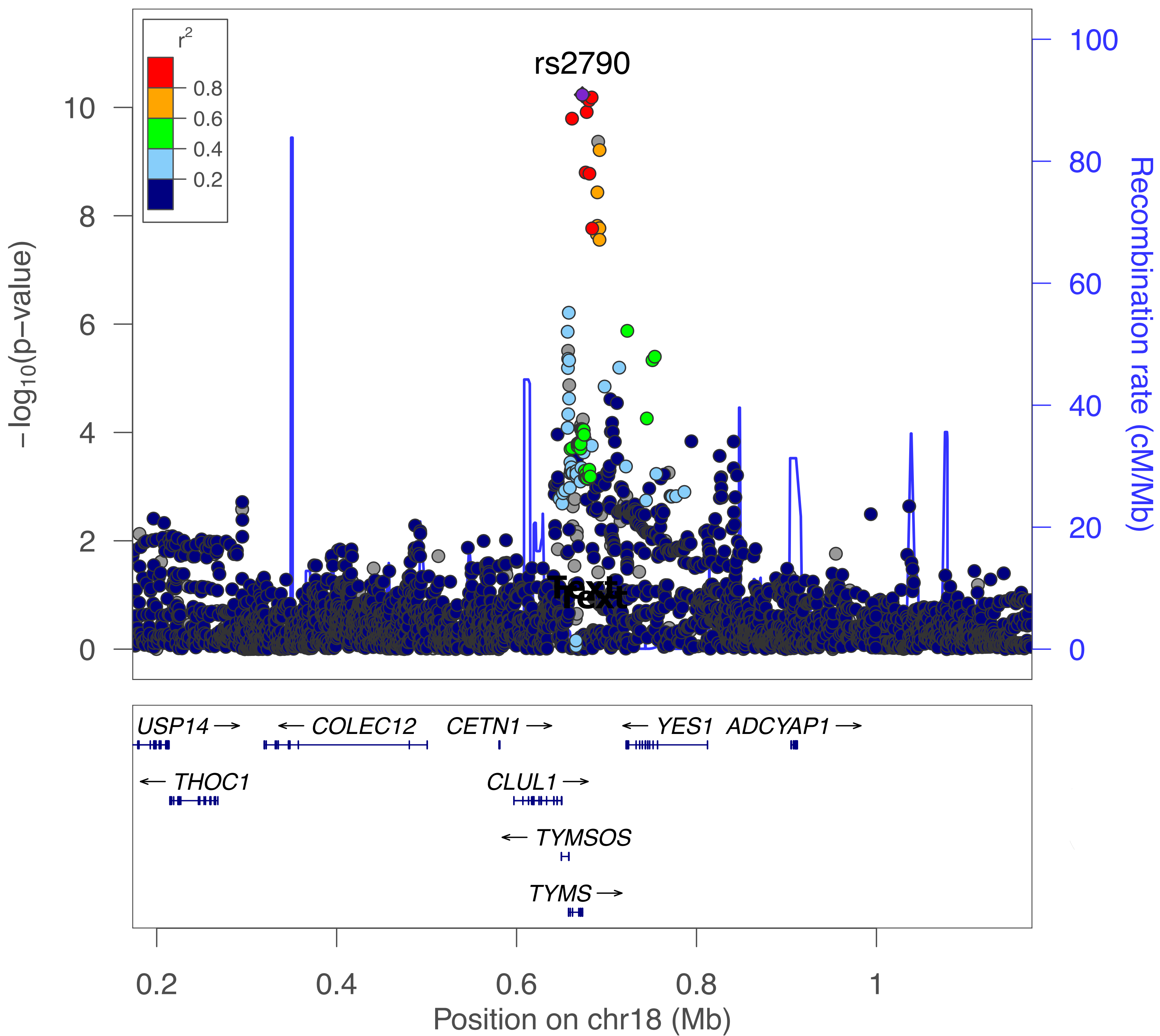

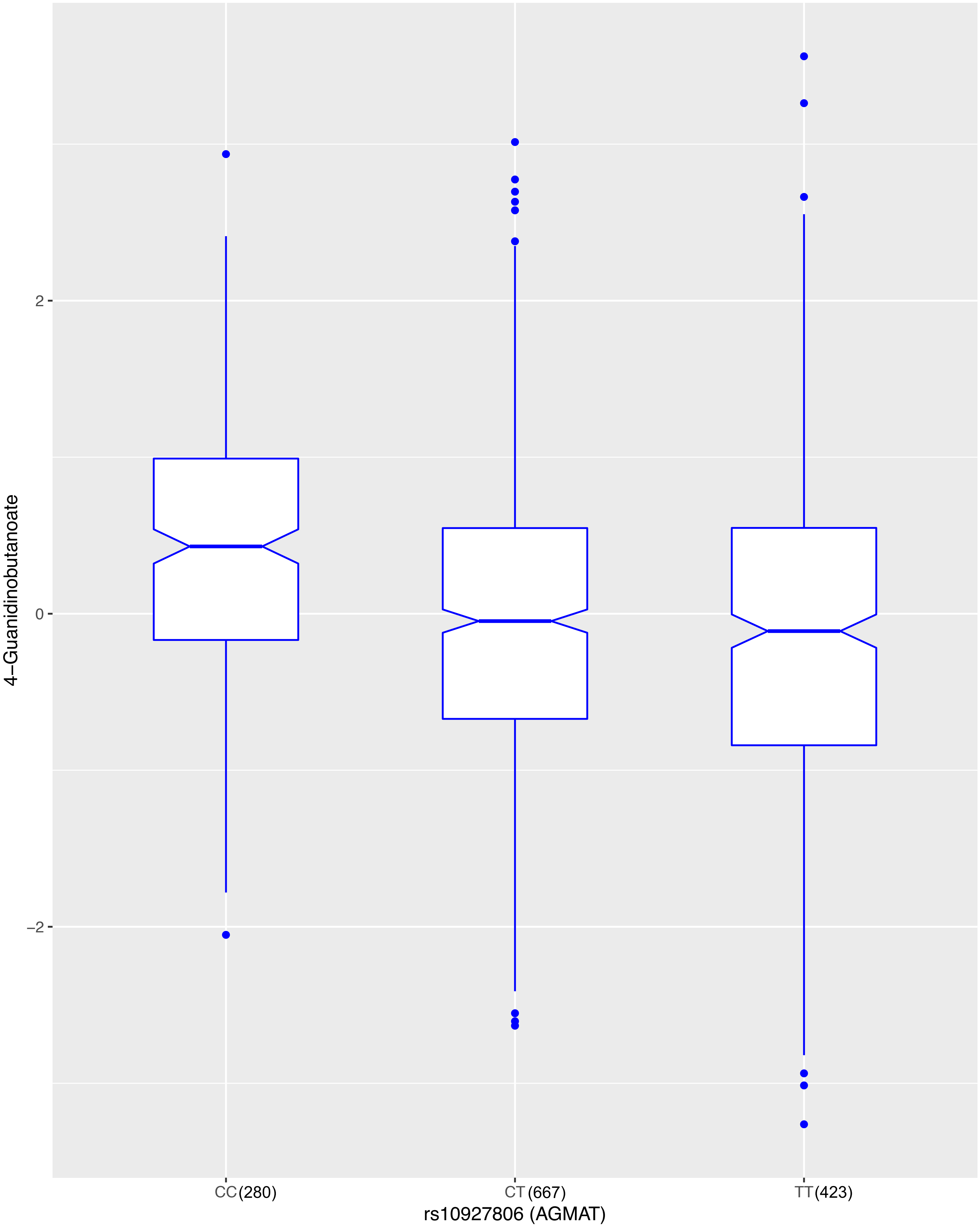

2
0
−2
4−Guanidinobutanoate
CC (280)
CT (667)
TT (423)
rs10927806 (AGMAT)

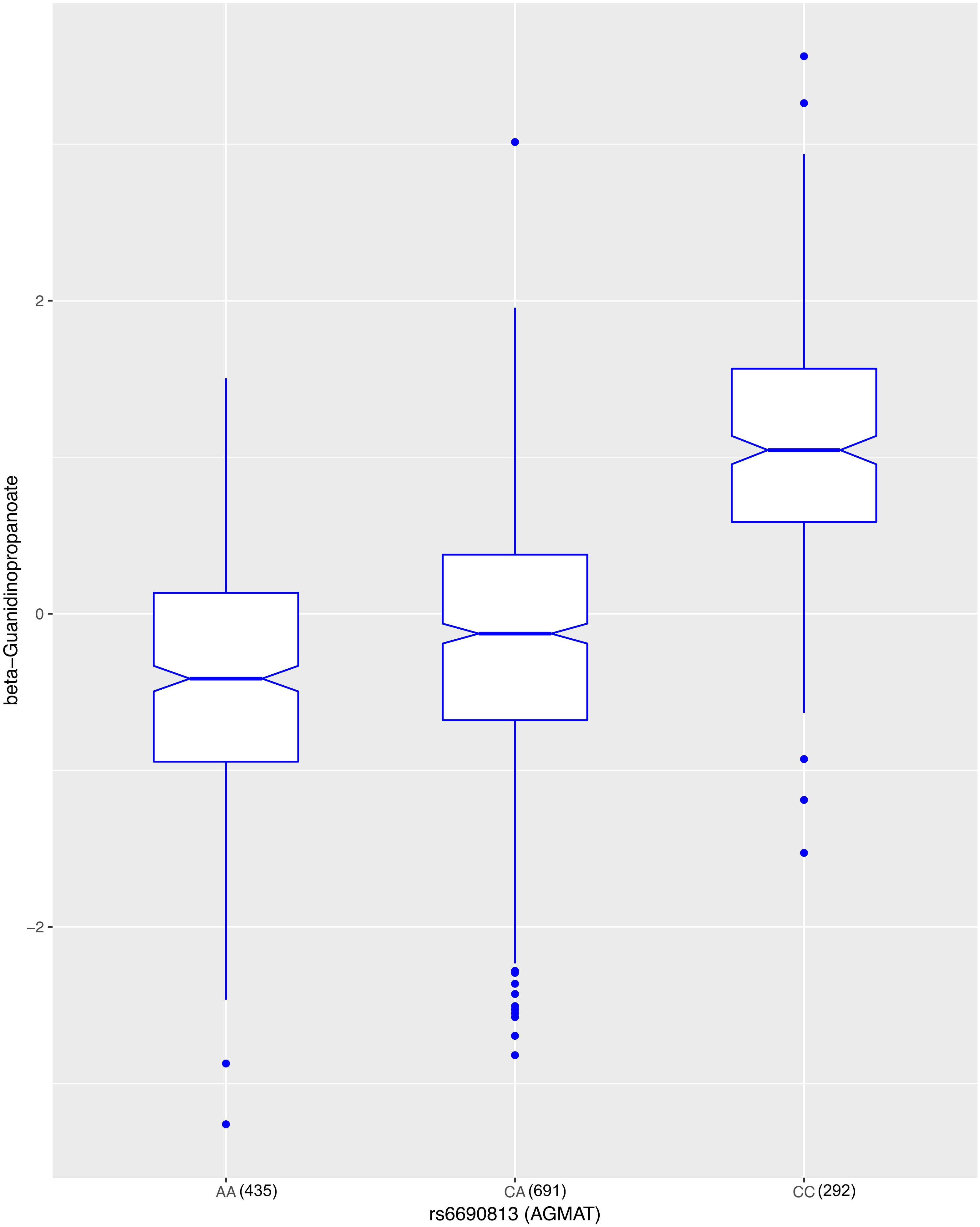
beta−Guanidinopropanoate
2
0
−2
AA (435)
CA (691)
CC (292)
rs6690813 (AGMAT)

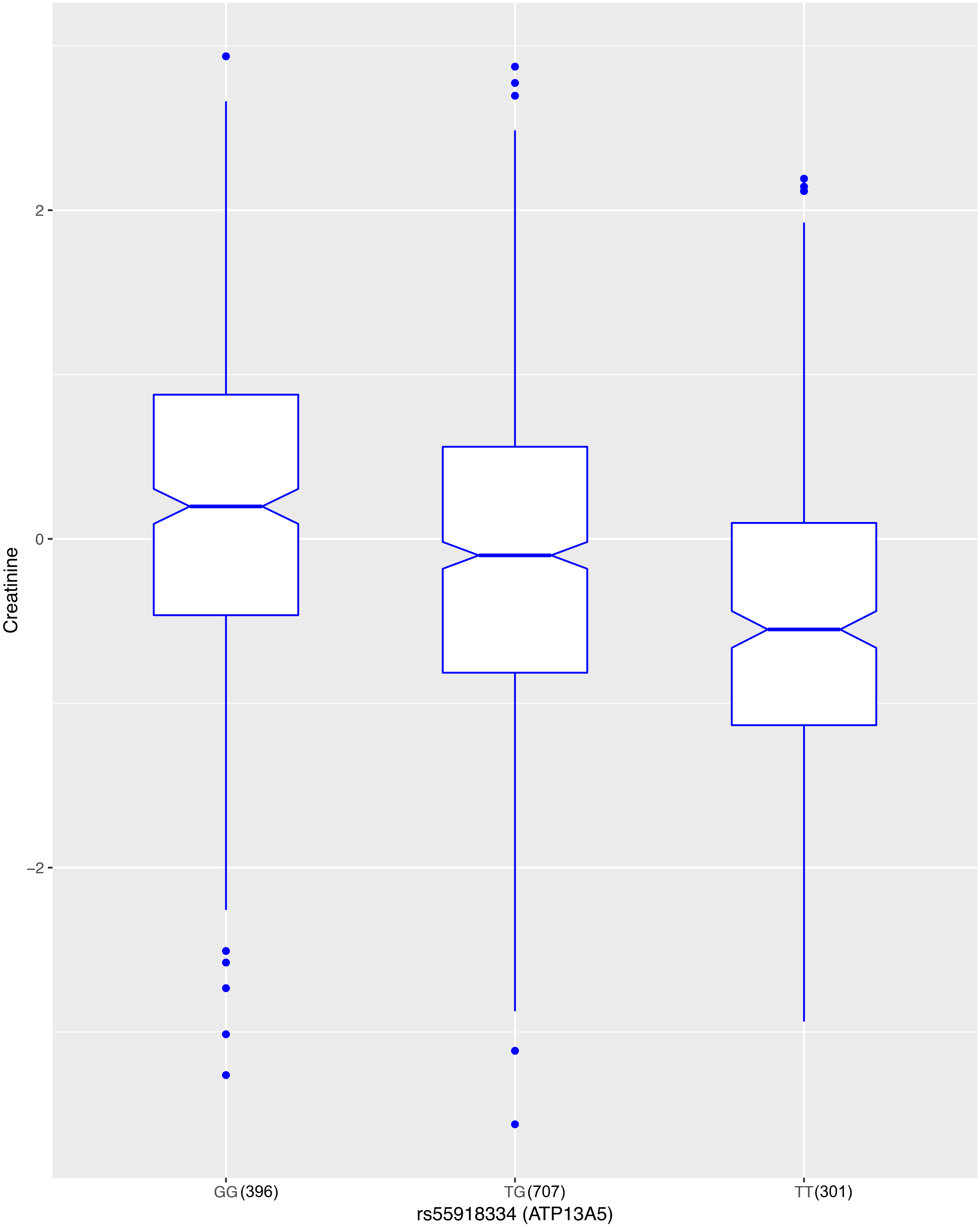

Creatinine
2
0
−2
GG(396)
TG(707)
TT(301)
rs55918334 (ATP13A5)

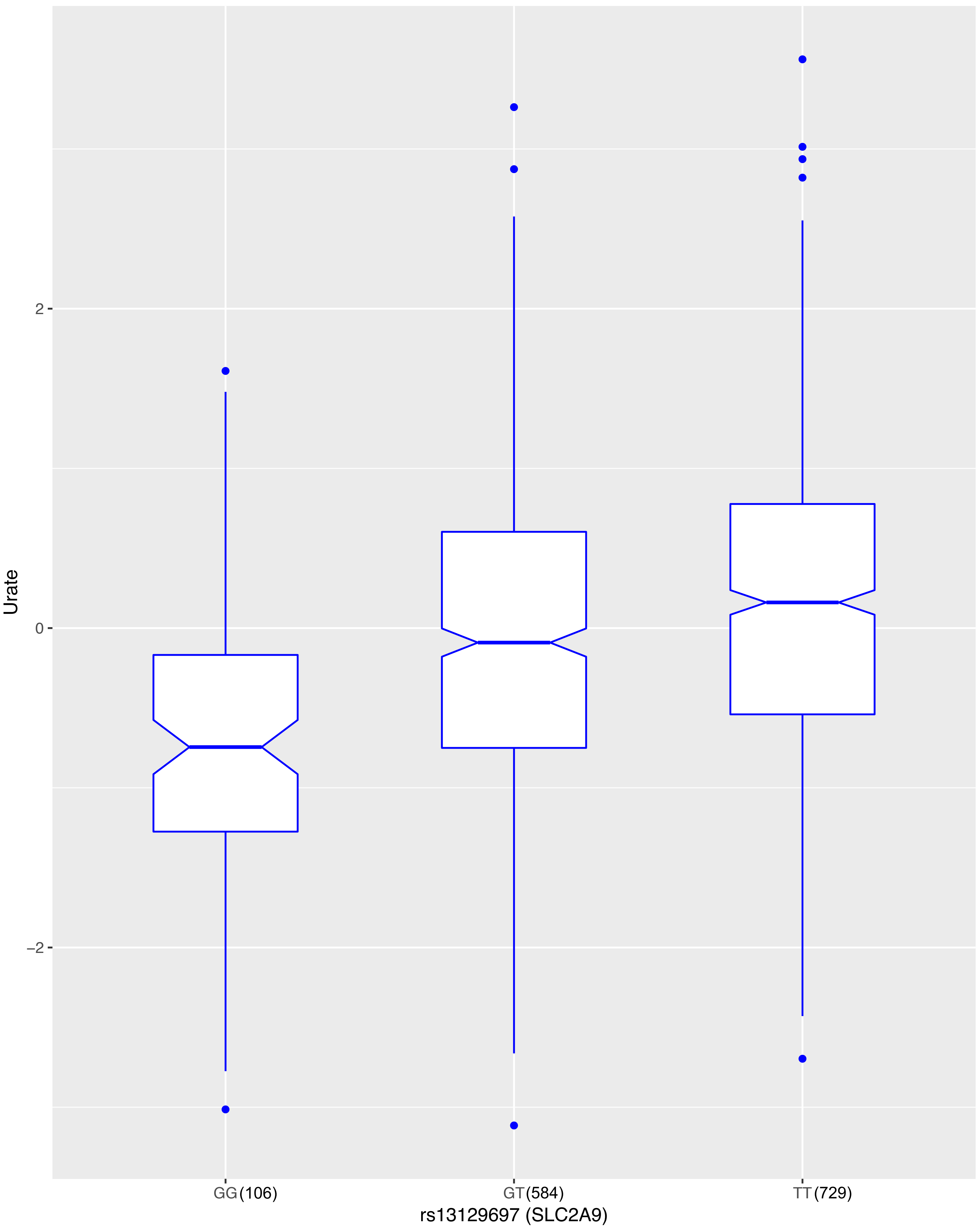
Urate
2
0
−2
GG (106)
GT (584)
TT (729)
rs13129697 (SLC2A9)

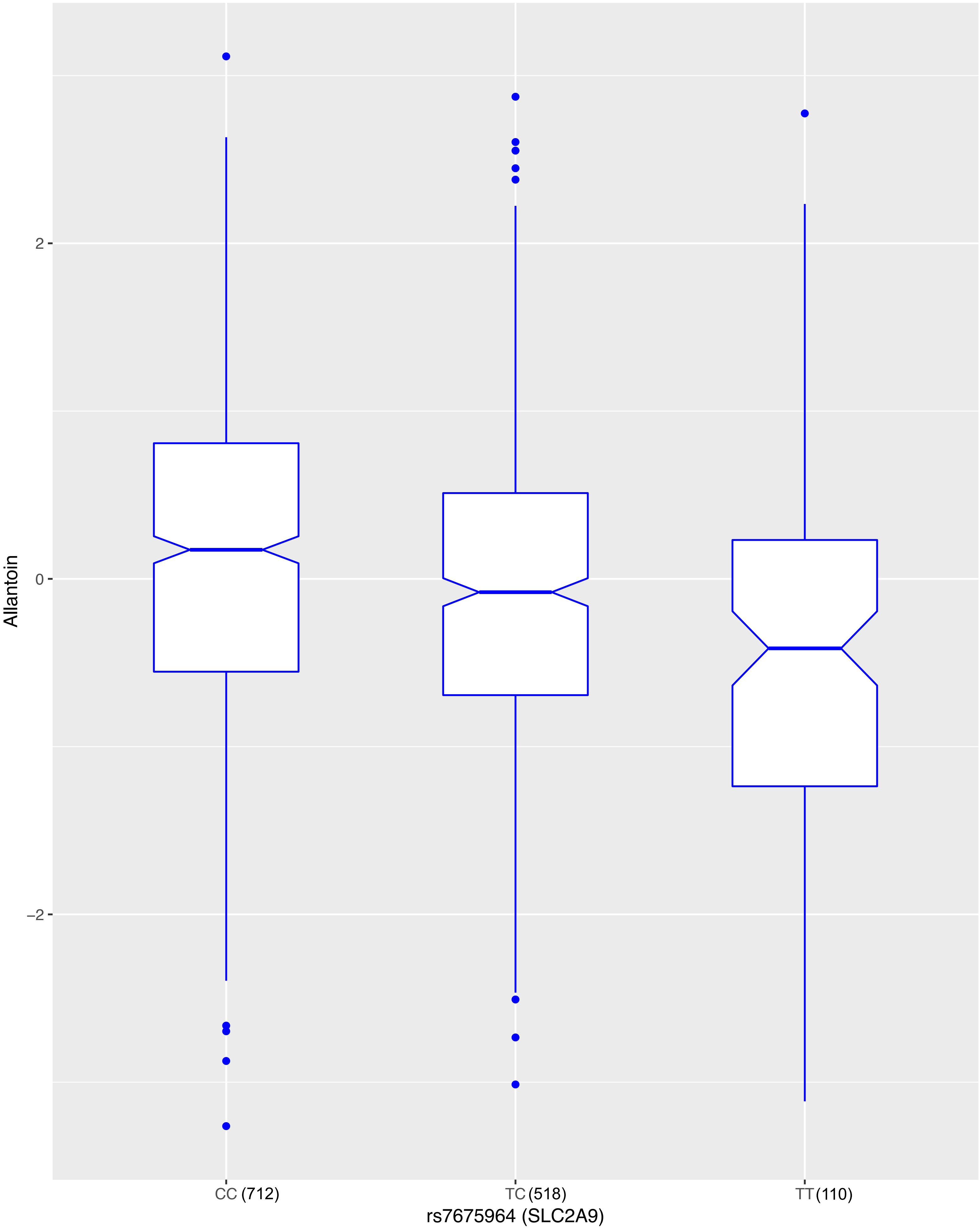
Allantoin
2
0
−2
CC (712)
TC (518)
TT (110)
rs7675964 (SLC2A9)

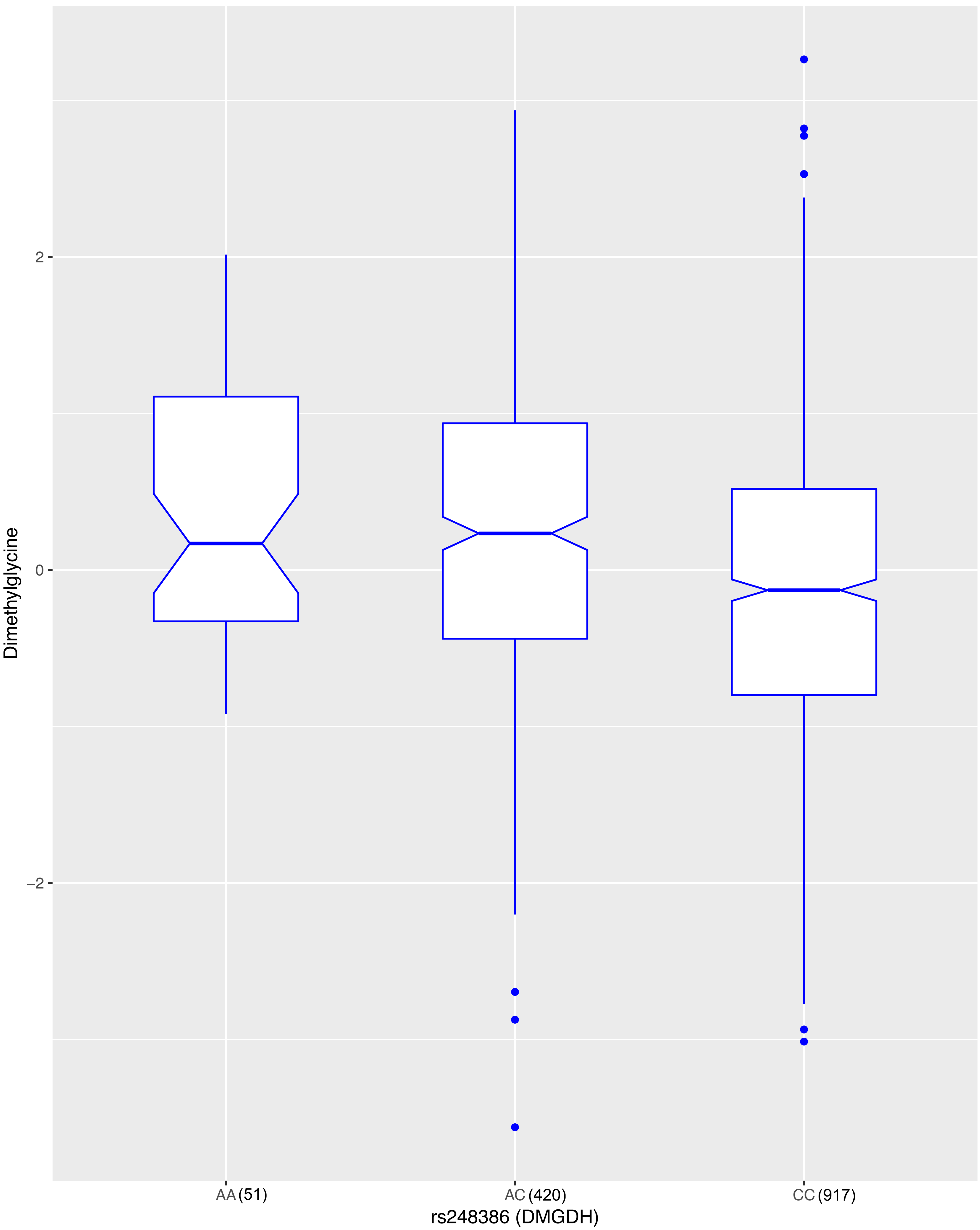

Dimethylglycine
2
0
−2
AA (51)
AC (420)
CC (917)
rs248386 (DMGDH)

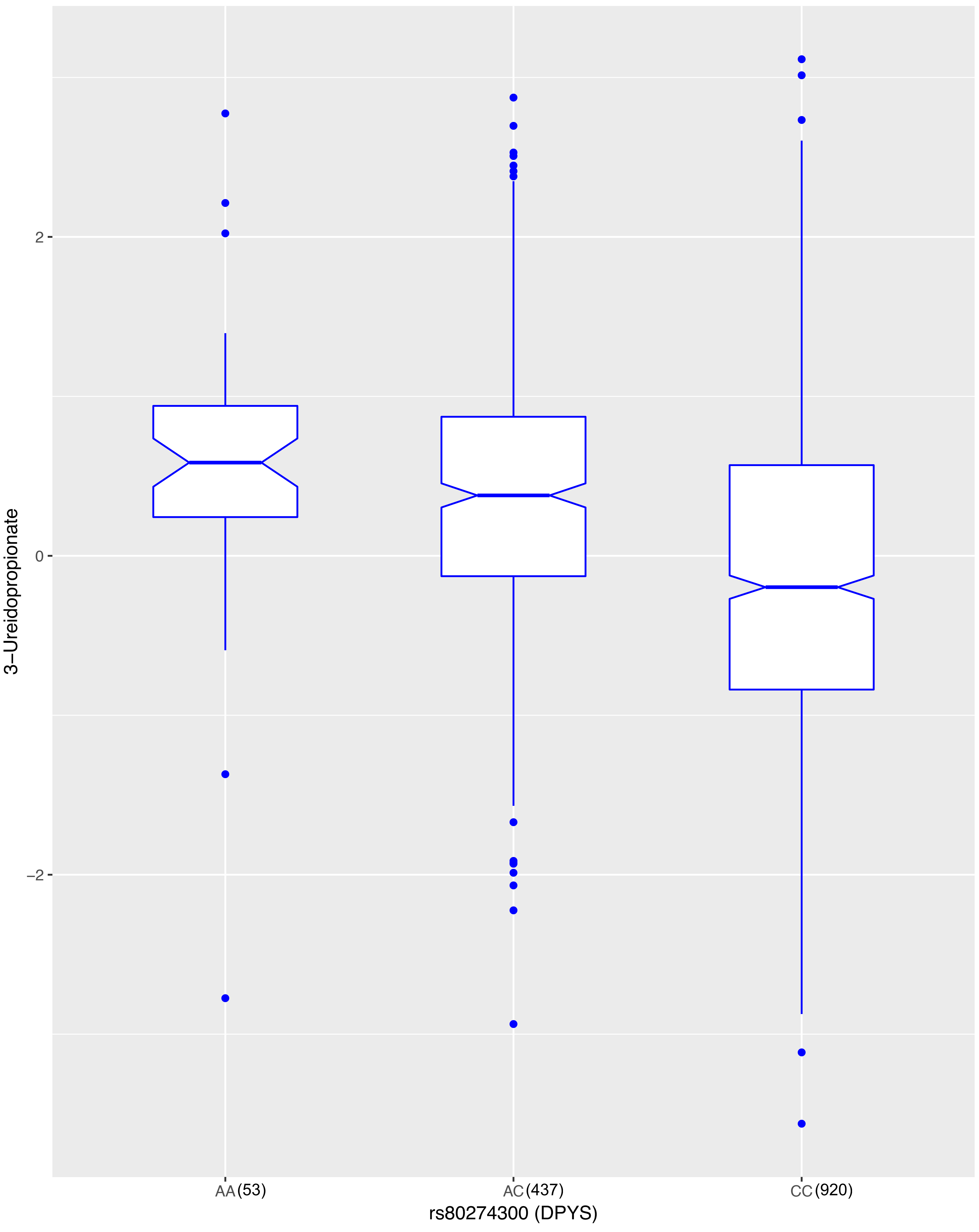

3–Ureidopropionate
2
0
–2
AA (53)
AC (437)
CC (920)
rs80274300 (DPYS)

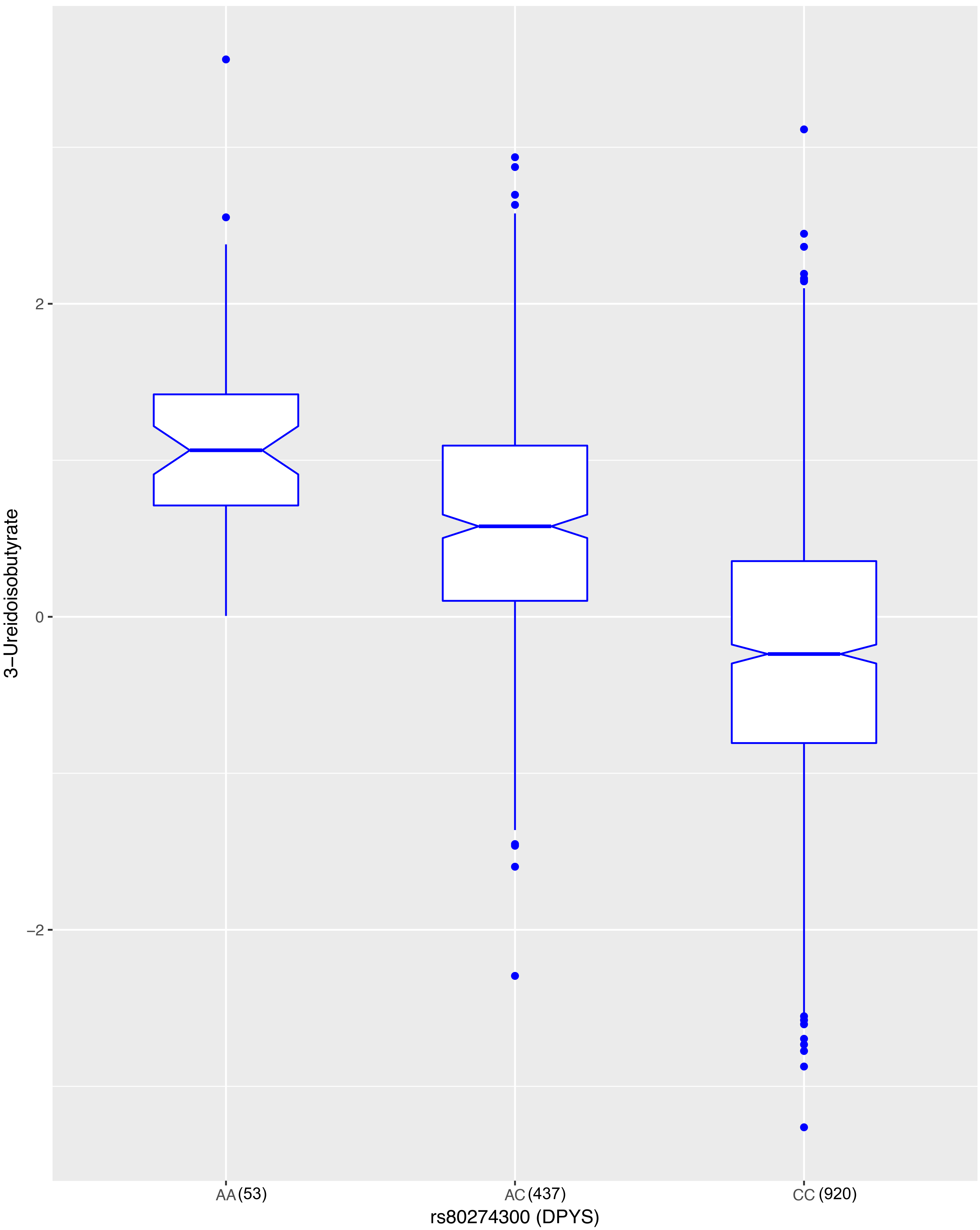
3−Ureidoisobutyrate
2
0
−2
AA (53)
AC (437)
CC (920)
rs80274300 (DPYS)

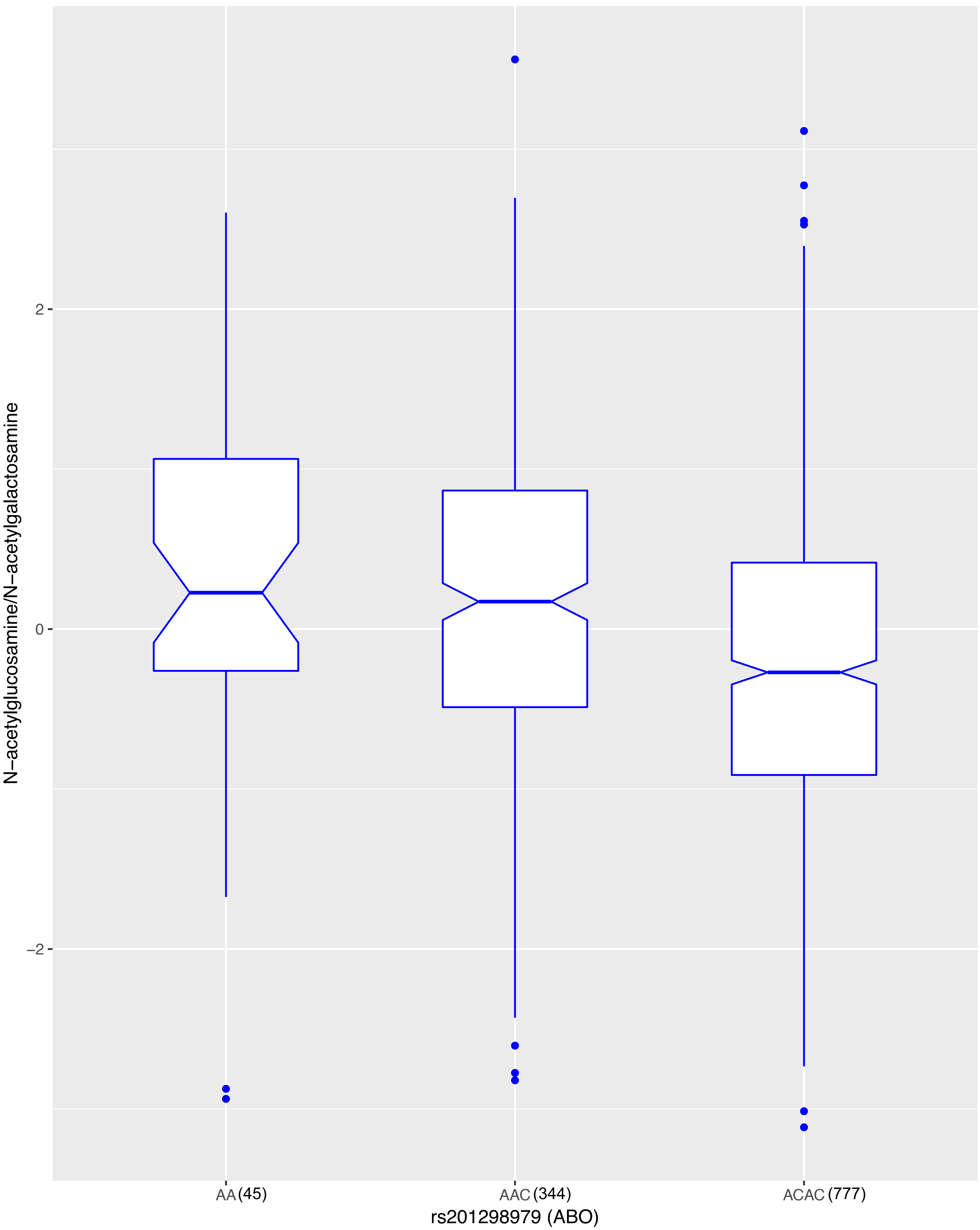

N−acetylglucosamine/N−acetylgalactosamine
2
0
−2
AA (45)
AAC (344)
ACAC (777)
rs201298979 (ABO)

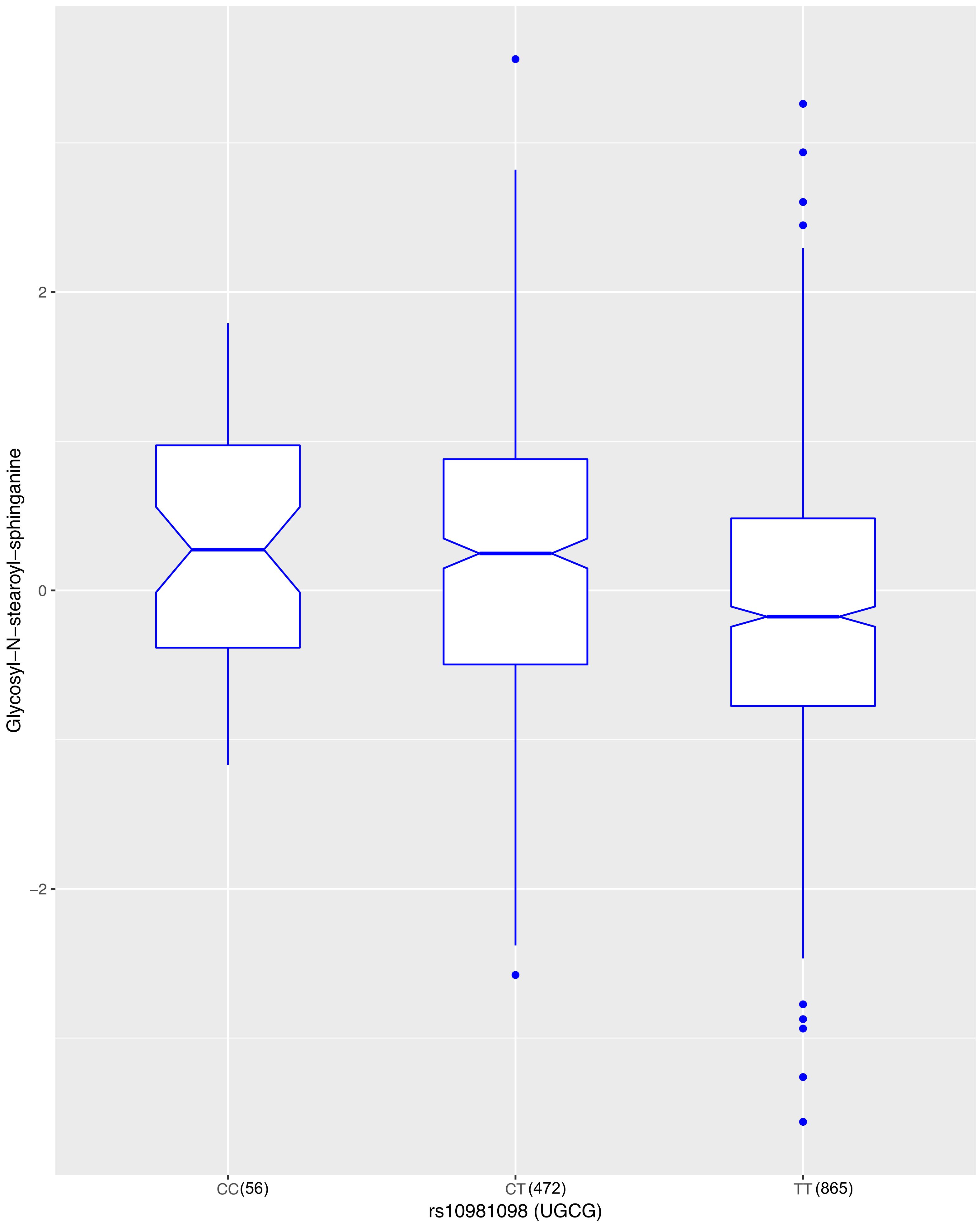

Glycosyl−N−stearoyl−sphinganine
2
0
−2
CC(56)
CT(472)
TT(865)
rs10981098 (UGCG)

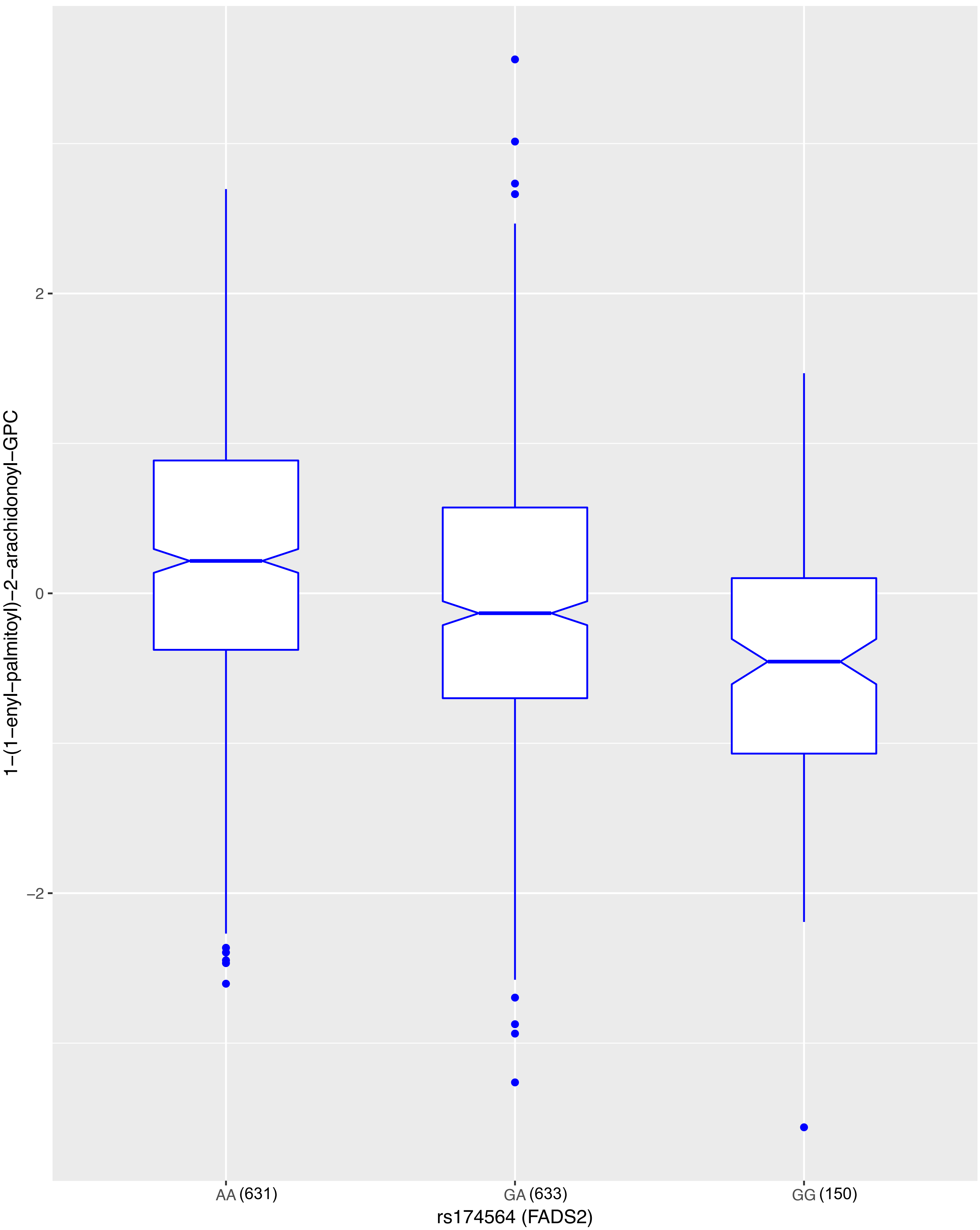

1−(1−enyl−palmitoyl)−2−arachidonoyl−GPC
2
0
−2
AA (631)
GA (633)
GG (150)
rs174564 (FADS2)

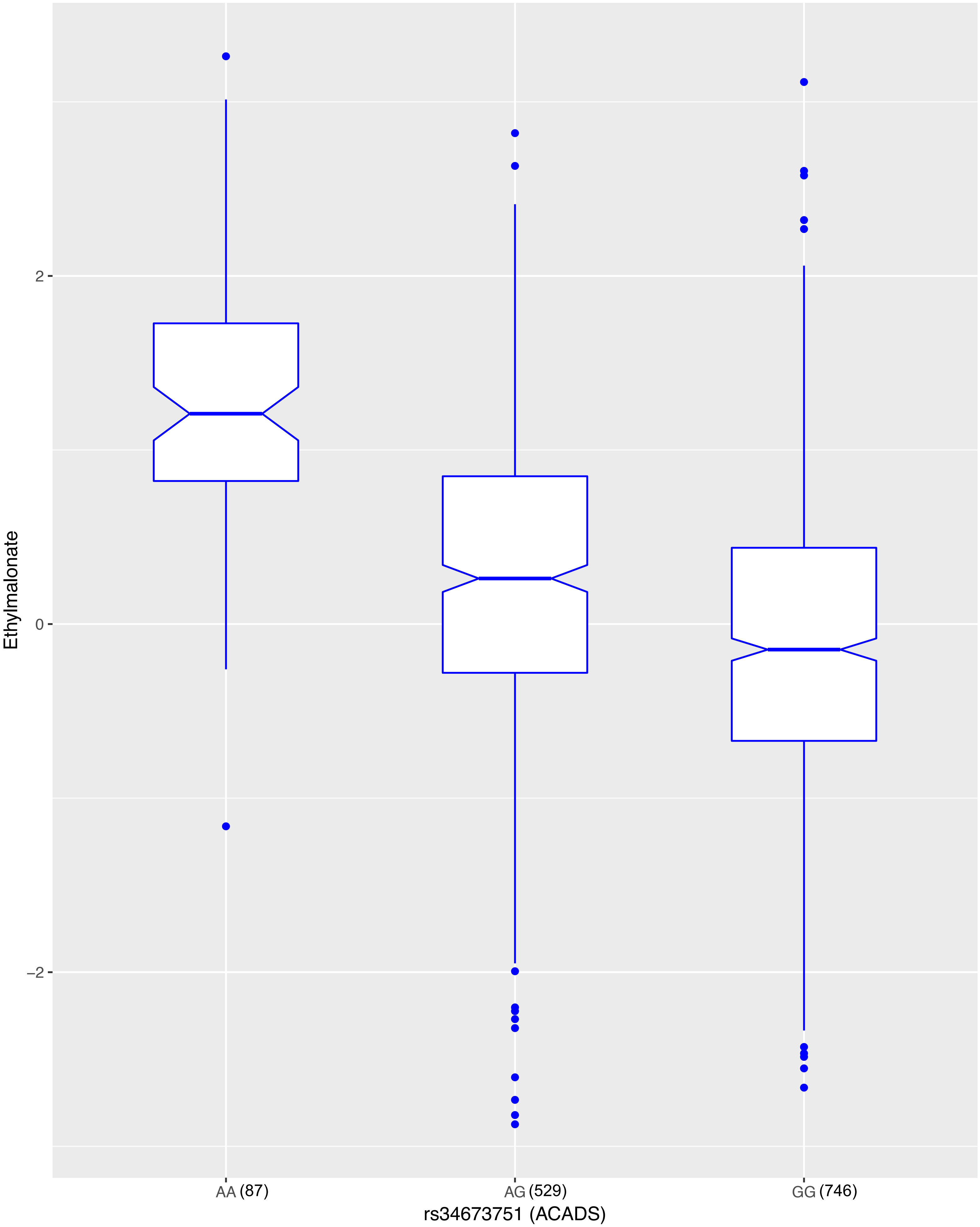
Ethylmalonate
2
0
−2
AA (87)
AG (529)
GG (746)
rs34673751 (ACADS)

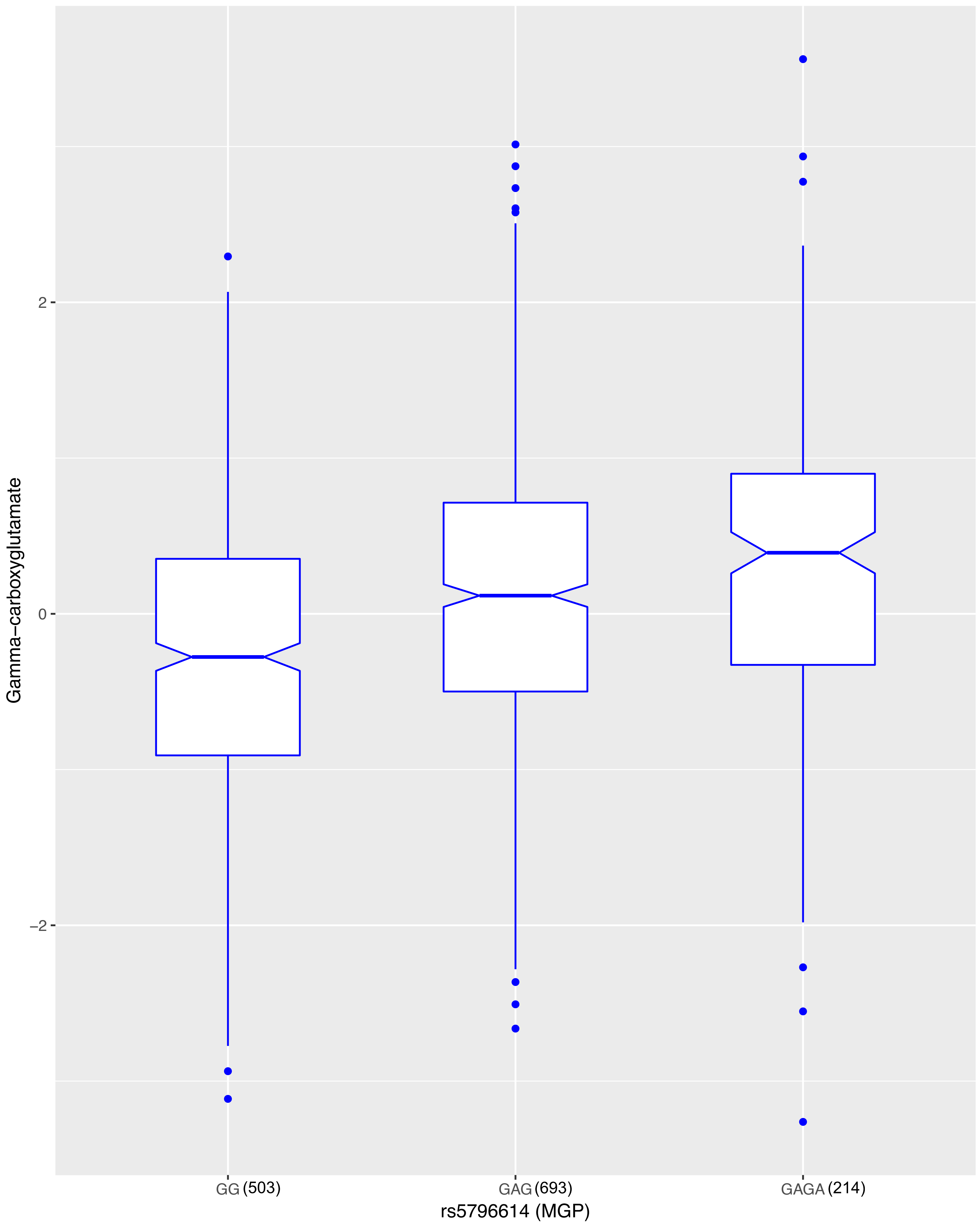
Gamma−carboxyglutamate
2
0
−2
GG (503)
GAG (693)
GAGA (214)
rs5796614 (MGP)

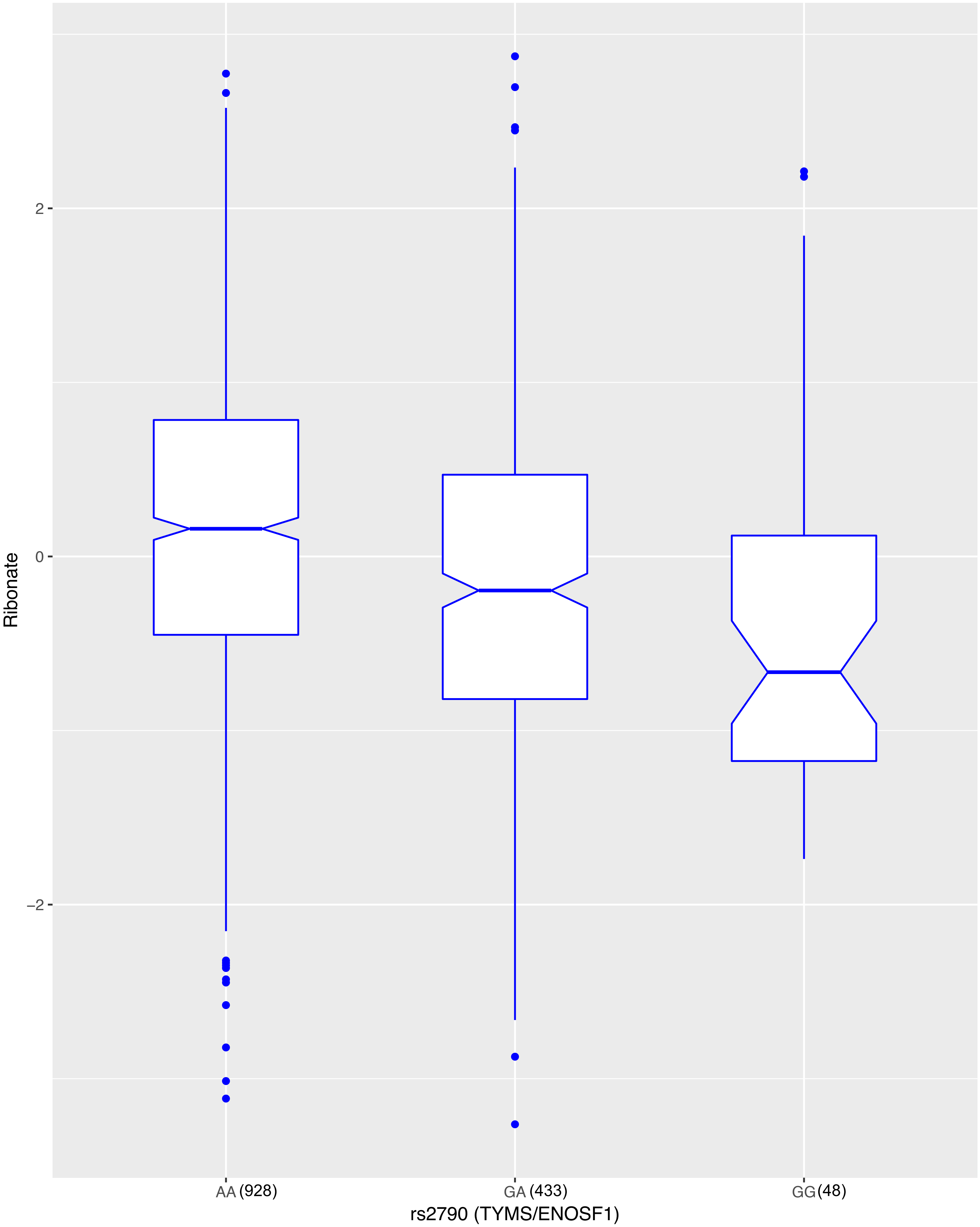

2
0
−2
Ribonate
AA (928)
GA (433)
GG (48)
rs2790 (TYMS/ENOSF1)
